# Key morphological features of human pyramidal neurons

**DOI:** 10.1101/2023.11.10.566540

**Authors:** Ruth Benavides-Piccione, Blazquez-Llorca, Asta Kastanauskaite, Isabel Fernaud, Silvia Gonzalez-Tapia, Javier DeFelipe

## Abstract

The basic building block of the cerebral cortex, the pyramidal cell, has been shown to be characterized by a markedly different dendritic structure among layers, cortical areas, and species. Functionally, differences in the structure of their dendrites and axons are critical in determining how neurons integrate information. However, within the human cortex, these neurons have not been quantified in detail. In the present work, we performed intracellular injections of Lucifer Yellow and 3D reconstructed over 200 pyramidal neurons, including apical and basal dendritic and local axonal arbors and dendritic spines, from human occipital primary visual area and associative temporal cortex. We found that human pyramidal neurons from temporal cortex were larger, displayed more complex apical and basal structural organization and had more spines compared to those in primary sensory cortex. Moreover, these human neocortical neurons displayed specific shared and distinct characteristics in comparison to previously published human hippocampal pyramidal neurons. Additionally, we identified distinct morphological features in human neurons that set them apart from mouse neurons. Lastly, we observed certain consistent organizational patterns shared across species. This study emphasizes the existing diversity within pyramidal cell structures across different cortical areas and species, suggesting substantial species-specific variations in their computational properties.

## Introduction

Exploring the biological basis of human abilities is of great significance for both fundamental and applied neuroscience. Nevertheless, the bulk of our current understanding of brain structure and behavior stems from research conducted on experimental animals. It has been shown that the human cerebral cortex presents some unique molecular, physiological and anatomical features, which highlighting the importance of directly studying the human brain (e.g., Oberheim et al., 2009; DeFelipe, 2015; Hodge et al., 2019; Berg et al., 2021; Campagnola et al., 2022; de Kock et al., 2023, Luria et al., 2023). The investigation of pyramidal neurons is particularly intriguing given that they are the most abundant neurons in the cerebral cortex. These neurons constitute the primary source of intrinsic excitatory cortical synapses. They also serve as the primary recipients of excitatory synapses and contribute to the majority of intra-areal projections, along with nearly all inter-areal and subcortical projections (reviewed in DeFelipe and Fariñas, 1992). Consequently, they play a pivotal role in the processing and transmission of cortical information within the brain. The basic structure of pyramidal neurons is shaped by a prominent apical dendrite arising from the soma, directed towards the pia mater, giving off a number of oblique collaterals that usually terminate in an apical tuft. From the base of the soma, several laterally or downward directed dendrites emerge forming the basal arbor. Pyramidal neurons are found in all cortical layers except layer I and they are commonly categorized according to their projection site (e.g., Jones, 1984; White, 1989; Nieuwenhuys, 1994). In fact, pyramidal neurons in distinct cortical layers and regions participate in different synaptic circuits, thereby segregating particular cortical functions (for review, see Barbas, 2015; D’Souza and Burkhalter, 2017; Rockland, 2019).

There are significant variations in the structure of pyramidal neurons, depending upon cortical layer, region, and species. These variations are believed to be pivotal for the functional specialization of cortical areas (e.g., Elston and Rosa, 1997; Jacobs et al., 2001; Bianchi et al., 2011; DeFelipe, 2011; Elston et al., 2011; Elston and Manger, 2014; Eyal et al., 2016; Mohan et al., 2015; Jacobs et al., 2015, 2018; Benavides-Piccione et al., 2020, 2021; Galakhova et al., 2022; Kanari et al., 2023). Indeed, the dendritic tree structure influences the biophysical and computational properties of neurons, and these differences are crucial factors contributing to the variations in the functional organization of the cerebral cortex (reviewed in Segev and London, 2000; Stuart and Spruston, 2015; Fisek and Häusser, 2020; Poirazi and Papotsi, 2020). For example, pyramidal neurons in the granular prefrontal and temporal associational cortex of primates, including humans, exhibit larger, more branched, and more spinous structure compared to those found in other sensory areas (Elston et al., 2001; Jacobs et al., 1997, 2001; Elston and Rockland, 2002; Bianchi et al., 2013). In the occipito-temporal cortex of non-human primates, a pattern emerges where dendritic trees tend to become larger, more branched, and more spinous as one moves anteriorly from the primary visual area (V1) to the secondary visual area (V2), the fourth visual area (V4, or dorsolateral DL visual area), and the inferotemporal (IT) cortex. However, the degree of these differences varies across species (Elston et al. 1998a, b, 1999; Jacobs and Scheibel, 2002; Elston et al., 2005 a, b). In the human cortex, basal dendritic arbors have been studied in V2 (Brodmann’s area 18: BA18) and associative temporal area 20 of Brodmann (BA20). These studies also reveal a tendency for neurons to exhibit increased branching (with values surpassing those of non-human primate species) as one moves from the occipital to the temporal cortex (Elston et al., 2001). Nonetheless, comprehensive comparative studies exploring the morphology of human pyramidal neurons, encompassing both apical dendritic and axonal arbors within occipito-temporal regions, remain unavailable. Thus, in the current study, we undertook an examination of the geometry of pyramidal neurons within the primary visual area (BA17) and the anterior temporal association areas (BA20 and BA21) of the human cortex. Our aim was to gain further insights into the structural intricacies of the distinct dendritic and axonal components of these neurons. Our findings reveal distinct variations in size and architectural arrangement among human pyramidal neurons across neocortical regions, encompassing their apical, basal, and axonal compartments. Specifically, larger cell and soma sizes, more complex apical and basal dendritic patterns, thicker dendrites and axons, longer distal dendritic segments and higher spine number, distinguish human temporal neurons from primary visual neurons. These neocortical neurons also exhibit distinctive features that set them apart from other previously analyzed human and rodent cortical pyramidal neurons, while also presenting some shared organizational patterns across species.

## Materials and Methods

### Tissue Preparation

Samples were obtained at autopsy (1–4 h post-mortem) from the Unidad Asociada Neuromax — Laboratorio de Neuroanatomía Humana, Facultad de Medicina, Universidad de Castilla-La Mancha, Albacete; Laboratorio Cajal de Circuitos Corticales UPM-CSIC, Madrid, Spain; and Instituto de Neuropatología Servicio de Anatomía Patológica, IDIBELL-Hospital Universitario de Bellvitge, Barcelona, Spain. The tissue was obtained following national laws and international ethical and technical guidelines on the use of human samples for biomedical research purposes. In the present study, we used a total of 7 human cases (table 1), some of which were used as controls in previous studies unrelated to the present investigation (Benavides-Piccione et al., 2013, Dominguez-Álvaro et al., 2021). Upon removal, the brains were immersed in cold 4% paraformaldehyde in 0.1 M phosphate buffer, pH 7.4 (PB) and sectioned into 1.5-cm-thick coronal slices. Small blocks from the neocortex were then transferred to a second solution of 4% paraformaldehyde in PB for 24 h at 4 °C. Vibratome sections (300 μm) of the occipital (Brodmann’s area 17; BA17) and temporal cortex (Brodmann’s area 20 and 21; BA20 and BA21; Garey, 1994) were obtained in the coronal plane. Specifically, blocks were dissected in all individuals from the lateral BA17 at the occipital pole and from the lateral BA20 and BA21 at 2-3 cm from the temporal pole. These regions were compared with previously published human CA1 pyramidal cell reconstructions at the level of the hippocampal body and from the middle of the CA1 pyramidal cell layer of the dorsal hippocampus in the mouse (see Benavides-Piccione et al., 2020 for further methodological details).

**Table 1.** Summary of human cases included in the study. F: female; M: male; L: left; R: right.

| **Case** | **Age** | **Sex** | **Side** | **Cause of death** |
| --- | --- | --- | --- | --- |
| **AB1** | 45 | M | L | Lung cancer and cardiorespiratory arrest |
| **AB2** | 53 | F | L | Pulmonary septic shock |
| **AB7** | 66 | M | L | Bladder carcinoma |
| **IF6** | 85 | M | L | Pneumonia and interstitial pneumonitis |
| **IF10** | 66 | M | L | Bronchopneumonia and cardiac failure |
| **IF12** | 52 | M | L | Disseminated carcinoma and bronchopneumonia |
| **M16** | 40 | M | L | Traffic accident |

### Intracellular injections and immunocytochemistry

Sections were prelabeled with 4,6-diamidino-2-phenylindole (DAPI; Sigma, St Louis, MO), and a continuous current was used to inject individual neurons with Lucifer yellow (8 % in 0.1; Tris buffer, pH 7.4; LY) in layer 3a of the occipital BA17 and temporal cortex BA20 and BA21 (**Figure 1**). LY was applied to each injected cell by continuous current until the distal tips of each cell fluoresced brightly, indicating that the dendrites were completely filled and ensuring that the fluorescence did not diminish at a distance from the soma. Following the intracellular injections, the sections were immunostained for LY using rabbit antisera against LY (1:400 000; generated at the Cajal Institute) diluted in stock solution (2% bovine serum albumin, 1% Triton X-100, and 5% sucrose in PB). The sections were then incubated in biotinylated donkey anti-rabbit IgG (1:100; Amersham, Buckinghamshire, United Kingdom) and streptavidin-conjugated Alexa fluor 488 (1:1000; Molecular Probes, Eugene, OR, United States of America). Finally, the sections were washed and mounted either with ProLong GoldAntifade Reagent (Invitrogen Corporation, Carlsbad, CA, USA) or with Glycerol 50% in PB. See Elston et al. (2001) and Benavides-Piccione et al. (2013) for further details of the cell injection methodology.

**Figure 1.**
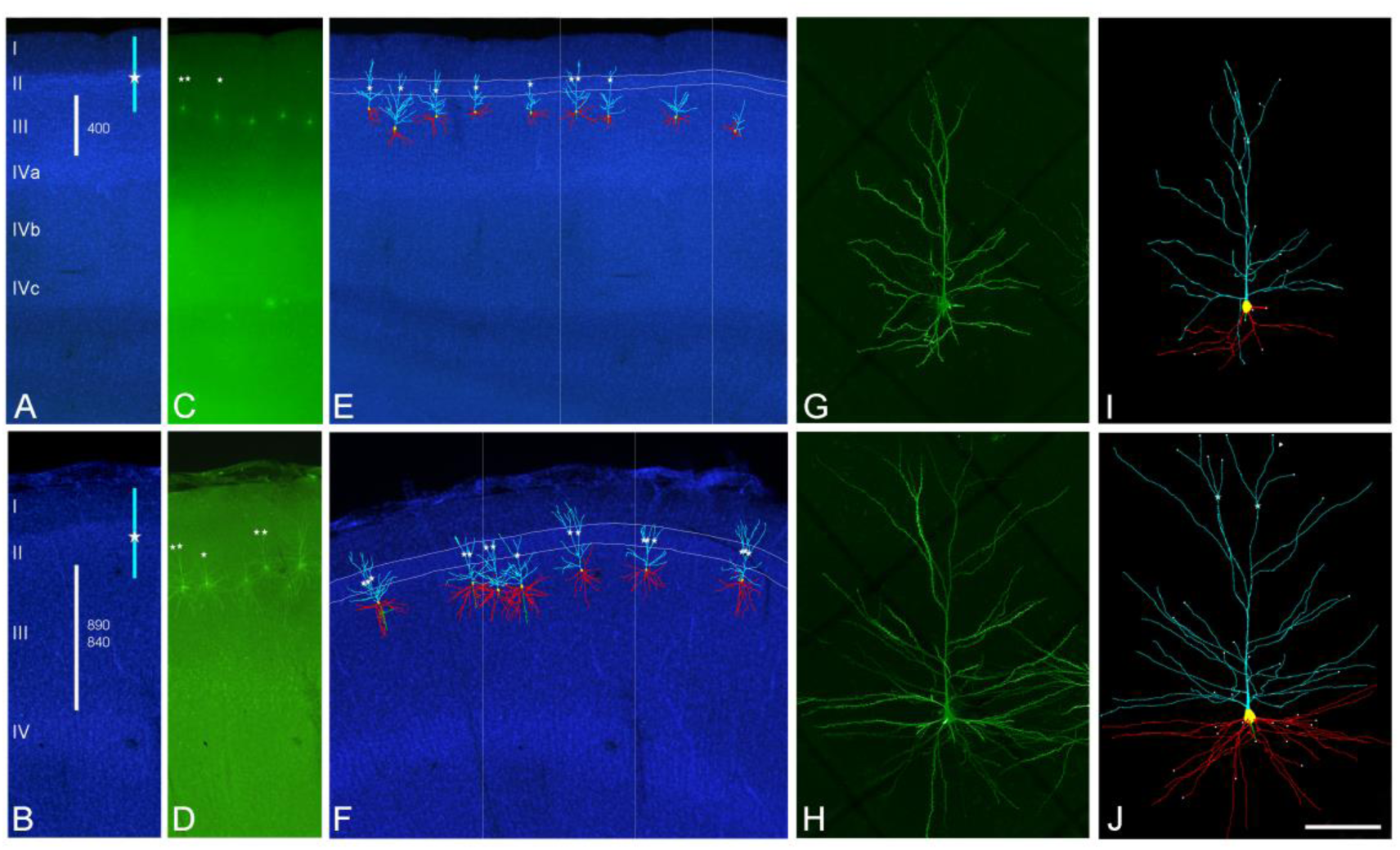
Confocal microscopy images of human neurons injected with LY in occipital BA17 cortex (top images) and temporal BA20 cortex (bottom images). (**A, B**) DAPI staining to illustrate cortical thickness (white bars) and estimated values (in microns) in occipital (A) and temporal (B; top: BA20, bottom: BA21) cortex. Blue bars indicate the approximate position of labeled neurons apical arbors. White stars represent the initiation of the apical tuft. (**C, D**) Labeled pyramidal neurons in layer 3a of the region shown in A and B. (**E, F**) 3D reconstructed neurons superimposed on the DAPI staining showing apical (light blue) and basal (red) arbors. Apical tufts, if present (white stars), were found to initiate within layer II (highlighted in white lines). Boxed area corresponds to regions shown in A–D. (**G, H**) High magnification image z projection showing an injected BA17 (G) and BA20 (H) pyramidal cell. (**I, J**) 3D reconstruction of the same neurons shown in G and H. Incomplete dendritic segments are highlighted in white. Scale bar (in J) =500 μm in A–F and 100 μm in G–J.

### Cell Reconstruction

Sections were imaged with a Leica TCS 4D confocal scanning laser attached to a Leitz DMIRB fluorescence microscope. Fluorescent labeling profiles were imaged, using an excitation wavelength of 491 nm to visualize Alexa fluor 488. Consecutive stacks of images (x63 glycerol; voxel size, 0.240 × 0.240 × 0.29 μm^3^) were acquired to capture the apical and basal dendritic arbors. Since intracellular injections of the pyramidal cell were performed in coronal sections (300 μm-thick), the part of the dendritic arbor nearest the surface of the slice from which the cell soma was injected (typically at a depth of ∼30– 50μm from the surface) was lost. Using a similar method of intracellular injection, Krimer et al. (1997) estimated that the reconstruction of neurons represented approximately two–thirds of the total dendritic arbor of pyramidal neurons. Nonetheless, it is important to mention that the percentage of the basal arbor and apical arbor included within the section may vary in each cell, depending on how parallel the main apical dendrite runs with respect to the surface of the slice. In the present study, neurons were included for apical analysis if they had a main apical dendrite of at least 200 µm in length. Furthermore, in some cases distal apical dendrites (apical tufts) were not included in the analysis. Specifically, due to technical limitations, apical tufts were included within the section in 19 out of 55, 40 out of 90, and 20 out of 50 pyramidal neurons in BA17, BA20, and BA21, respectively.

Data points of neuron morphology of each pyramidal cell included in the analysis (55 from BA17, 90 from BA20 and 50 from BA21 neurons) were extracted in 3D using Neurolucida 360 (MicroBrightfield; **Figure 2**). Briefly, the apical and basal dendrites, as well as the axon and soma, were described through 3D points, delimiting the different segments that form the cell arbor. These points have an associated diameter that provides the information of the varying thickness of the dendrite at that particular point varying along the length of the dendrite. Axons were only traced if they were included within the section over a length of at least 50 µm. The soma was defined through a set of connected points tracing the contour of the soma both in 2D and 3D.

**Figure 2.**
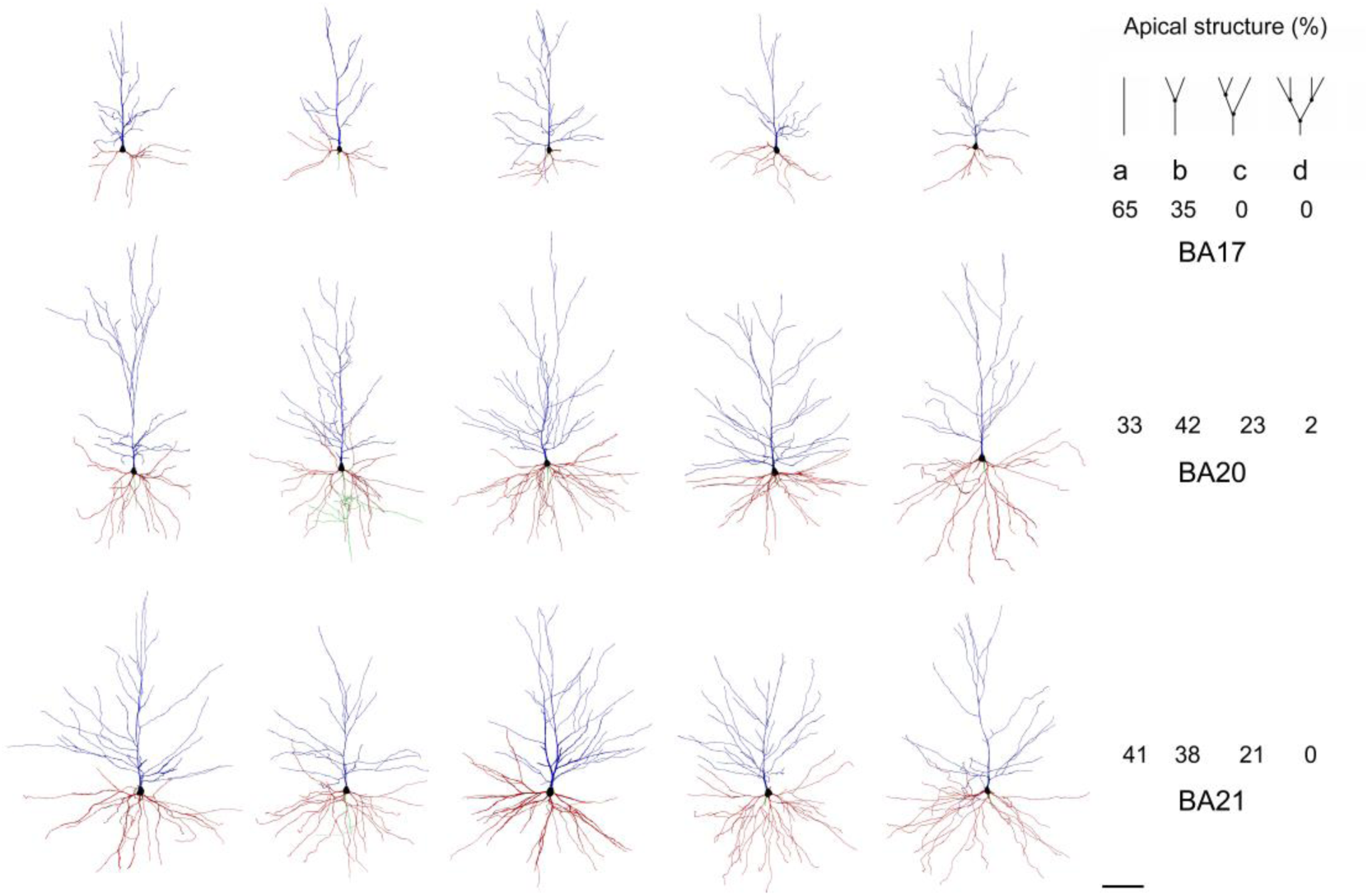
Example drawings of the apical (blue) and basal (red) dendritic arbors of human BA17 (top), BA20 (middle) and BA21 (bottom) pyramidal neurons. Axons, if present, are shown in green. On the right, the corresponding prevalence percentage of the different main apical branching patterns (top right): **a**, 0 bifurcations; **b**, 1 bifurcation; **c**, 2 bifurcations; **d**, 3 bifurcations —within the first 200 µm— is shown in each cortical region. Scale bar =100 µm.

### Quantitative Analysis

Several morphological variables were extracted using Neurolucida software. Some of the features measured did not depend on the entirety of the reconstructed cell and can thus be considered as full measurements: soma size (including 2D cross sectional area estimated by measuring the area of the maximum perimeter of the soma and 3D surface area and volume); dendritic/axonal average segment diameter; segment length; segment surface area; and segment volume; as well as axonal varicosity density (defined as a swelling of the axon exceeding the typical variation in diameter of the adjacent axonal shafts per axonal length) and intervaricosity distance (defined as the distance between two adjacent axonal varicosities). Other morphological variables do depend on the entirety of the cell, and, thus, may only partially describe the cell and can be considered ‘non-full’ measurements: area and volume of the dendritic arbor (2D and 3D convex hull); total number of dendrites; total number of nodes; total dendritic length; total dendritic surface area; and total dendritic surface volume.

Dendritic spine density was also analyzed. Spines were counted on 10 horizontally projecting basal dendrites, randomly taken from different neurons (x63; voxel size, 0.075 × 0.075 × 0.29 μm^3^), from 2 individuals per BA area. Spine density was calculated every 10 μm from the soma to the distal tip of the dendrites using Imaris software. Spine density was also calculated as a mean according to the number of spines found on the dendrite divided by the corresponding dendritic length. An estimate of the total number of spines found in the basal dendritic tree of the pyramidal neurons was calculated by multiplying the mean number of spines per 10 μm of dendrite with the mean number of branches for the corresponding part of the dendritic tree, from the cell body to the distal tips of the dendrites. These numbers of dendritic intersections were previously corrected assuming that the basal arbors in coronal sections represent two thirds of the total basal arbor (see *Cell Reconstruction* for further information). Thereafter, the summing of these partial values accounts for the estimation of the total number of spines in the basal arbor (for further details on quantitative analysis, see Elston, 2001 and Benavides-Piccione et al., 2020).

Values were expressed as total numbers, as a function of the distance from the soma (Sholl analysis) and per dendritic segment branch order. Only dendritic segments that were completely reconstructed were included in the branch order analysis (see also **Figure 1**). All statistical analyses were performed using GraphPad Prism version 9.3.1 for Windows (GraphPad Software, San Diego, CA, United States of America). When morphological parameters were presented as mean values, the Kruskal–Wallis test was used to compare between the groups. Measurements reported as a function of the distance from the soma and per branch order were analyzed using repeated-measures ANOVA (Mixed-effects analysis was performed at the furthest distances, in the cases that missing values were present due to the fact that the distal tips of the different dendrites do not reach the same length). Correlation analysis between the parameters quantified was performed with non-parametric Spearman analysis since most parameters did not exhibit a normal distribution. Significant correlations were classified as weak (Spearman rho (r) value lower than 0.40), moderate (0.4 < r < 0.7) and strong (r > 0.7). Differences were considered to be significant when P < 0.05. Measurements are reported as mean ± SEM, unless otherwise indicated.

## Results

### Comparison between occipital and temporal regions

#### Pyramidal cell and soma size are larger in the temporal cortex

Layer 3a pyramidal neurons were analyzed in the human occipital cortical primary visual area BA17 and temporal associative areas BA20 and BA21 (**Figure 1 and 2**). The mean layer 3 thickness in which neurons were injected was 402.31 ±11.05 μm, 891.04 ±11.42 μm and 837.48 ±12.05 μm in A17, A20 and A21, respectively. Layer 3a soma and cell sizes were similar to each other in temporal areas and significantly larger than those in BA17 (**Figure 3A; Supplementary Figure 1A, B and 10; Supplementary Table 1a**).

**Figure 3.**
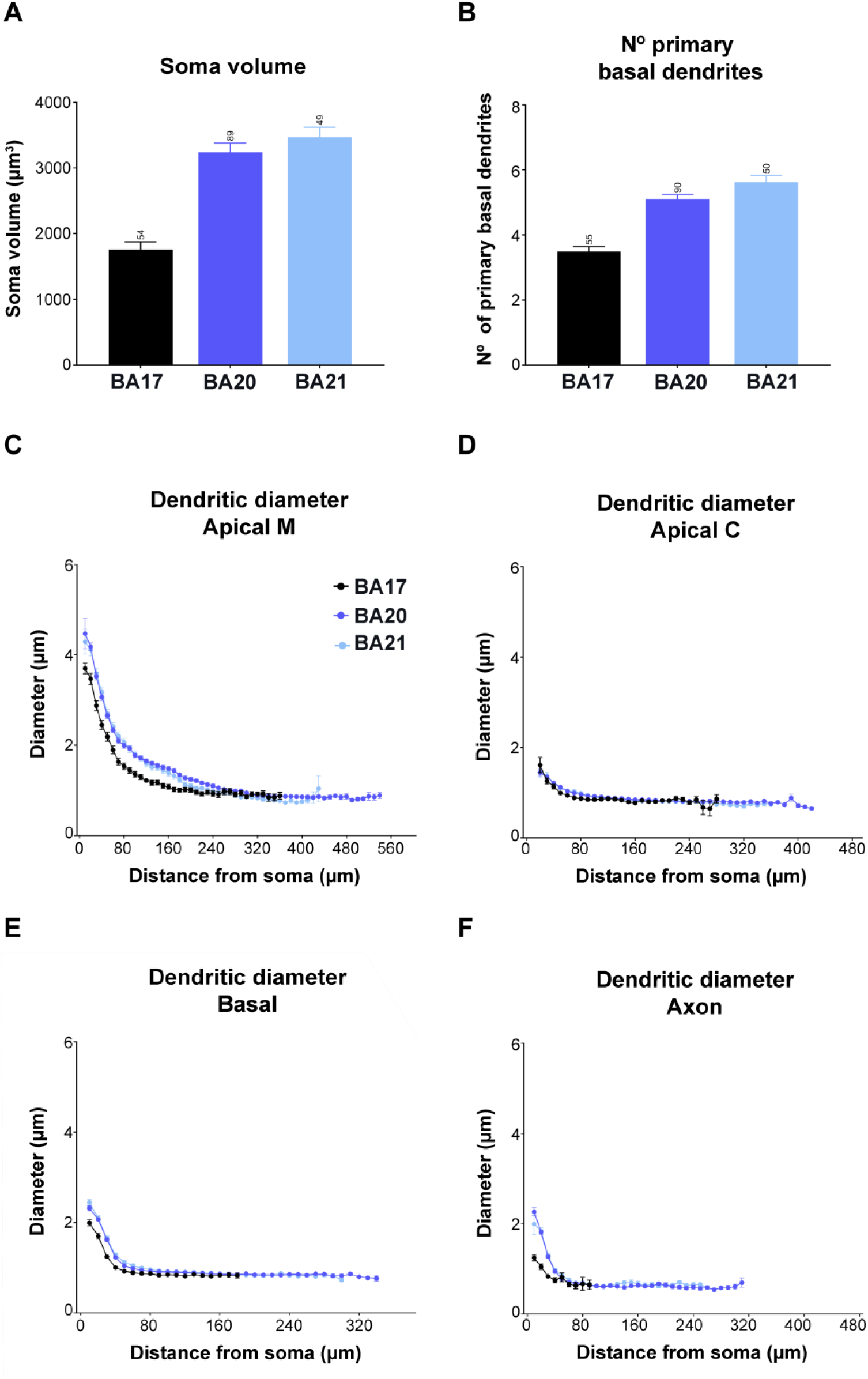
Graphs showing soma volume (**A**), the number of primary basal dendrites (**B**) and diameter distribution as a function of the distance from soma for the main apical dendrites (apical M; **C**), apical collateral dendrites (apical C; **D**), basal dendrites (**E**) and axon (**F**) from human BA17 (black), BA20 (dark blue) and BA21 (light blue) cortical areas. Measurements are reported as mean ± SEM. Additional graphs showing cell body cross-sectional area and surface area are shown in Supplementary Figure 1. The statistical significance of the differences is shown in Supplementary Tables 1 and 2.

#### Temporal apical and basal arbors are more complex than those in occipital cortex

Apical arbors emerging from the soma showed a reach of ∼500 μm in V1 and ∼630 μm in temporal areas (**Figure 1**). Sometimes the apical tufts were included within the section and were, thus, also analyzed. Tufts appeared when the apical arbors reached layer II (**Figure 1**). The length of the main apical dendrite (for simplicity, apical M) from the soma to the beginning of the tuft was 200.95 ±13.57 μm, 254.82 ±11.22 μm and 216.33 ±11.47 μm in BA17, BA20 and BA21, respectively. The estimation of the tuft extent was ∼300 μm in V1 and ∼380 μm in temporal areas (**Figure 1**).

The apical M branching structure was analyzed in dendrites of at least 200 µm in length, as an arbitrary measure (**Figure 2**). These neurons either showed an ascending course without branching (65%, 33%, 41% in BA17, BA20 and BA21, respectively), bifurcated once (35%, 42%, 38%), twice (0%, 23%, 21%), or three or more times (0%, 2%, 0%). **Supplementary Figures 2 and 3** show examples of these apical M dendrograms. Regarding apical collateral dendrites (for simplicity, apical C), these processes emerged mainly oblique to the apical M dendrite and were more numerous in temporal areas than in the occipital area (**Figure 1**). The basal arbors were composed of a number of primary basal dendrites that emerged from the soma, which were similar in both temporal areas and significantly higher than in the occipital cortex (**Figure 3B**, **Table 2 and Supplementary Table 1a**).

**Table 2.**
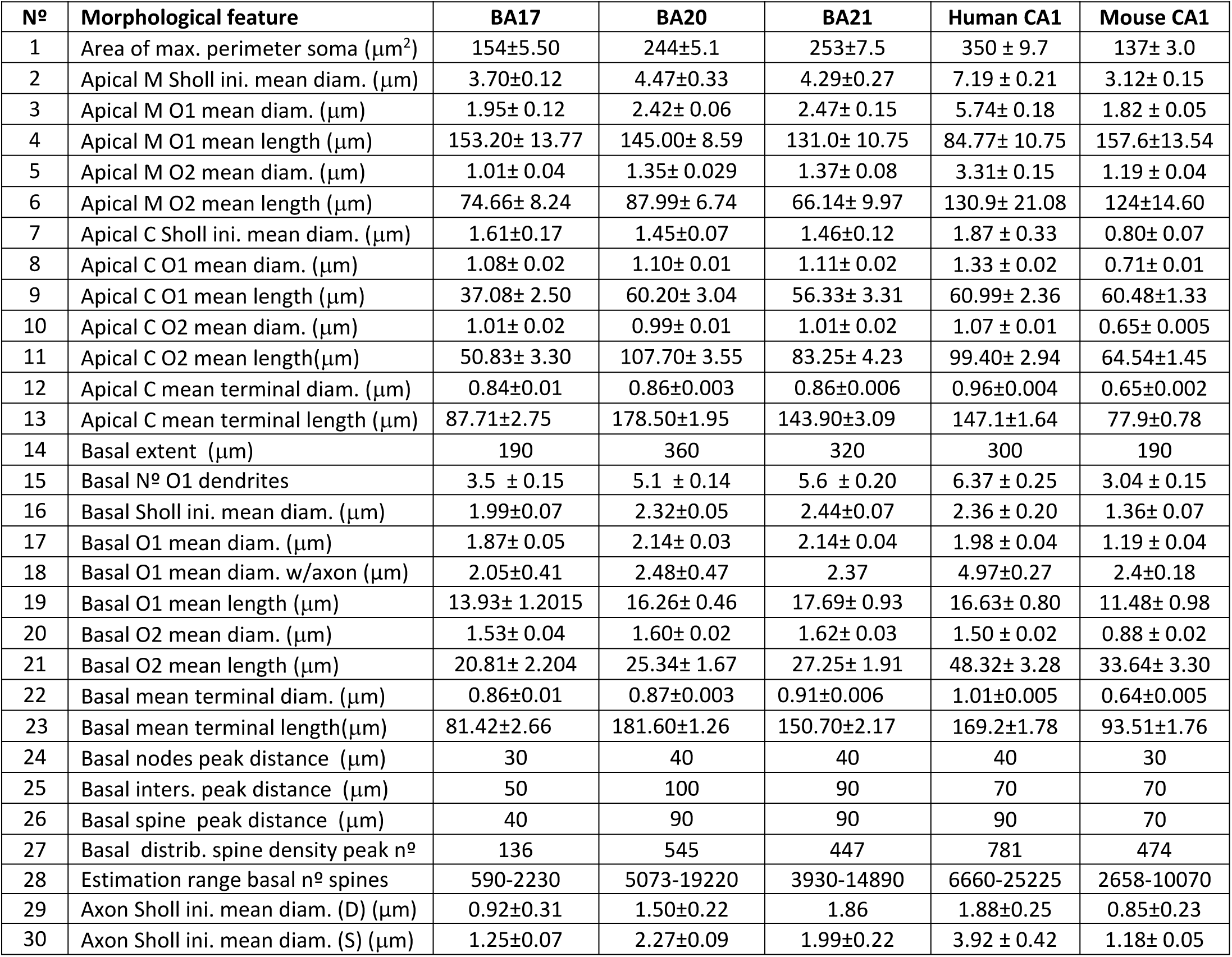
Summary of some measurements (mean±SEM) including soma size (n° 1), apical main (apical M; n° 2-6), apical collateral (apical C; n° 7-13) and basal (n° 14-28) dendritic diameters and lengths analyzed per Sholl analysis, at 10 μm initial (ini) Sholl distance, and per branch order analysis: first- and second-order (O1, O2) dendritic segments and axonal diameters (n° 29-30) emerging from dendrite (D) and soma (S), for human primary visual BA17, human associative temporal BA20/21 and human (HCA1) and mouse (MCA1) hippocampal CA1 regions. See also schematic Figure 11.

#### Apical M and basal dendrites are thicker in temporal cortex

The thickness of the apical M dendrite that emerged from the soma was similar in both temporal areas and thicker than that in the occipital cortex (**Figure 3C**). In all cases, the diameter gradually decreased from the soma along the length of the apical M dendrite. Temporal cortex values were significantly higher, from the beginning up to the first ∼250 microns, compared to V1 cortex (see **Supplementary Table 2** for statistical comparisons). From this distance onwards, the diameter values were similar (∼0.85 microns) in all cortical regions. The diameter of apical C dendrite was relatively similar in the 3 cortical regions and decreased along the first 50 μm from the soma (thickness from ∼1.5 microns to ∼0.8 microns, **Figure 3D**). However, at some points (between 50– 110 microns from the soma), significant differences were observed (**Supplementary Table 2**). The diameter of basal dendrites (**Figure 3E**) —from soma to distal tip— was similar between temporal cortical regions and significantly thicker than the occipital cortex (**Supplementary Table 2**), particularly for the first 150 μm, and then gradually decreased to reach a similar diameter (∼0.85μm) at a distance of ∼50 μm from the soma, and for the remaining distances, in all cortical regions.

Then, the structure sorted per dendritic segment branch order was analyzed (only dendritic segments that were completely reconstructed, thus excluding incomplete endings, were included in this analysis; see material and methods and **Figure 1I, J** for details). When taken together, mean segment diameters from apical M and C compartments were relatively similar between regions (**Supplementary Figure 1C; Supplementary Table 1b**). However, when sorted per branch order, temporal apical M dendrites showed significantly larger values for branch order 1 (O1) and 2 (O2) compared to those in the corresponding occipital cortex (branch orders 3 and 4 did not show this clear difference; **Figure 4A, Supplementary Table 3**). In all regions, the diameters decreased, as the branch order increased, with branch O1 being significantly thicker compared to the corresponding remaining orders. Apical C dendritic segment diameters were relatively similar between neocortical areas, both when taken together (**Supplementary Figure 1C; Supplementary Table 1b**) and when sorted per branch order (**Figure 4A**). In all cortical regions, values decreased as branch order increased — a trend which was found to be significant in some cases (**Supplementary Table 3)**.

**Figure 4.**
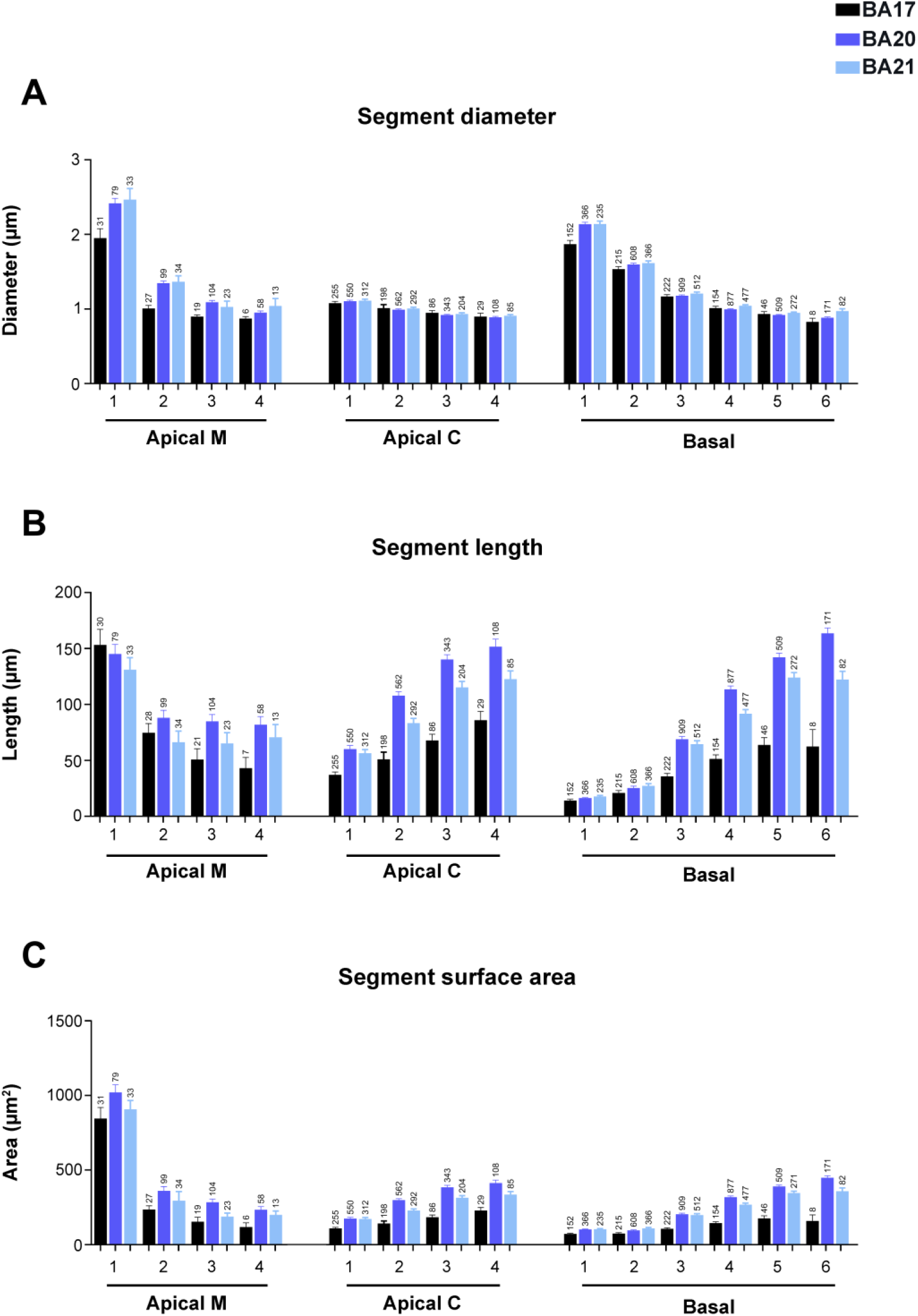
Graphs showing dendritic segment average diameter (**A**), length (**B**) and surface area (**C**), expressed per branch order (1, 2, 3, etc.) and per dendritic compartment: main apical dendrite (Apical M), apical collateral dendrites (Apical C), and basal arbor (Basal) from human BA17 (black), BA20 (dark blue) and BA21 (light blue) cortical areas. Measurements are reported as mean ± SEM. Only dendritic segments that were complete, and thus excluding incomplete endings, were included in this analysis. An additional graph showing segment volume is shown in Supplementary Figure 5A. The statistical significance of the differences is shown in Supplementary Table 3.

When basal dendrites were analyzed per dendritic segment (**Figure 4A and Supplementary Figures 1 C–F** and **5A**), the diameter of basal O1 dendrites was found to be significantly larger in both temporal areas compared to occipital cortex (**Supplementary Table 3**) and similar for the remaining orders. In addition, branch O1 segments had the thickest diameters and progressively decreased in all cortical areas. Significant differences were found mainly between branch orders 1–3 in all cortical regions (**Supplementary Table 3**).

#### Distal dendritic segments are longer in the temporal Apical C and basal dendrites

Regarding the length of the apical M segments, the values were similar between cortical regions, both when taken together (**Supplementary Figure 1D**) and per branch order (**Figure 4B**). In all regions, branch O1 segments were of similar length (∼150 microns). At higher branch orders, the apical M length decreased as the branch order increased, and although values were slightly larger in the temporal cortex compared to the occipital cortex, most branch orders showed no significant differences between regions (**Supplementary Table 3**). Thus, the branch O1 segments showed the highest values in all regions compared to the remaining orders (significant differences were found in all cortical regions, (**Figure 4B and Supplementary Table 3**).

The apical C dendritic segment length values were significantly larger in the temporal cortex, at all branch orders (particularly in area 20), compared to the occipital cortex (**Figure 4B and Supplementary Figure 1D**). The length values of the collateral dendritic segments increased as the branch order increased and these differences were found to be significant in most cases and branch orders (**Figure 4B and Supplementary Table 3**). The length of the basal dendritic segments (**Figure 4B**) from the temporal cortex were significantly longer (particularly area 20) than the occipital cortex at most branch orders

#### Dendritic surface area and volume are larger in temporal neurons

The surface area (**Figure 4C and Supplementary Figure 1E**) of apical M dendritic branch O1 segments were similar (∼900 square microns) in all 3 regions (**Supplementary Table 3**). At higher branch orders, the values were larger in the temporal cortex compared to the occipital cortex, at most branch orders, although most differences were not significant (**Supplementary Table 3**). Branch O1 showed significantly higher values compared to the remaining orders in all cortical regions. Decreasing values were observed as the branch order increased — with such differences found to be significant between most branch orders in the three cortical regions. Apical C dendritic segment surface area values (**Figure 4C and Supplementary Figure 1E**) were significantly higher in the BA20/21 cortex than in the BA17 cortex, both when taken together and at most branch orders. Values showed an increasing trend as the branch order increased; this trend that was found to be significant in most cases and branch orders (**Figure 4C and Supplementary Table 3**). Regarding the surface area of the basal dendritic segments, neurons in the temporal cortex presented significantly higher values than in the occipital cortex at most branch orders (**Figure 4C**; **Supplementary Table 3**). Values increased in all regions as the branch order increased, but to a higher extent in the temporal areas (**Supplementary Table 3**).

The volume of the apical M dendritic segments (**Supplementary Figures 1F and 5A**) was also larger in the temporal cortex (particularly area 20) than in the occipital cortex in most cases (**Supplementary Table 3**). Values decreased as the branch order increased and were significantly larger in branch O1 compared to the remaining orders (**Supplementary Table 3**). Regarding the volume of apical C and basal segments, there was a trend of increasing values with increasing branch orders in the temporal cortex (see Supplementary Figure 5A and Supplementary Table 3). However, in the occipital cortex, these values did not show this increase as the branch order increased.

#### Intermediate segments are thicker and shorter than terminal segments, whereas terminal segments are of similar diameter but variable length

Dendritic segments were then further analyzed according to their position in the dendritic arbor: intermediate (meaning a segment that is between nodes and, thus, bifurcates) and terminal segments (meaning a segment that no longer bifurcates and, thus, ends). **Supplementary Figure 4** shows mean values from both positions (intermediate and terminal dendritic segments) and both compartments (apical C and basal dendrites). Intermediate segments were thicker, shorter and had less surface area and volume than terminal segments, both in apical C and basal compartments from all areas (**Supplementary Figures 4 and 5B, C and Supplementary Table 4** for statistical comparisons). In all cortical regions, intermediate basal segments were of similar width and thicker than apical C segments (which were also of similar width between regions). Terminal apical C and basal segments were of similar width in all regions. However, temporal (particularly BA20) intermediate and terminal segments were longer and had larger surface area and volume than occipital terminal segments.

We then further analyzed the intermediate and terminal segments according to their branch order. Apical C intermediate segments showed quite similar diameters (∼1.2 μm) between branch orders and cortical regions (**Figure 5A**; see **Supplementary Table 5** for statistical comparisons). Basal intermediate segments showed a similar decreasing pattern as branch order increased, with similar values between cortical regions, except branch O1 segments, which showed significantly thicker values in temporal cortical areas (**Figure 5A, Supplementary Table 5**). Terminal segments were thinner than intermediate segments and had a similar value (∼0.8 μm) regardless of the branch order, dendritic compartment or cortical region (**Figure 5B, Supplementary Table 5**).

**Figure 5.**
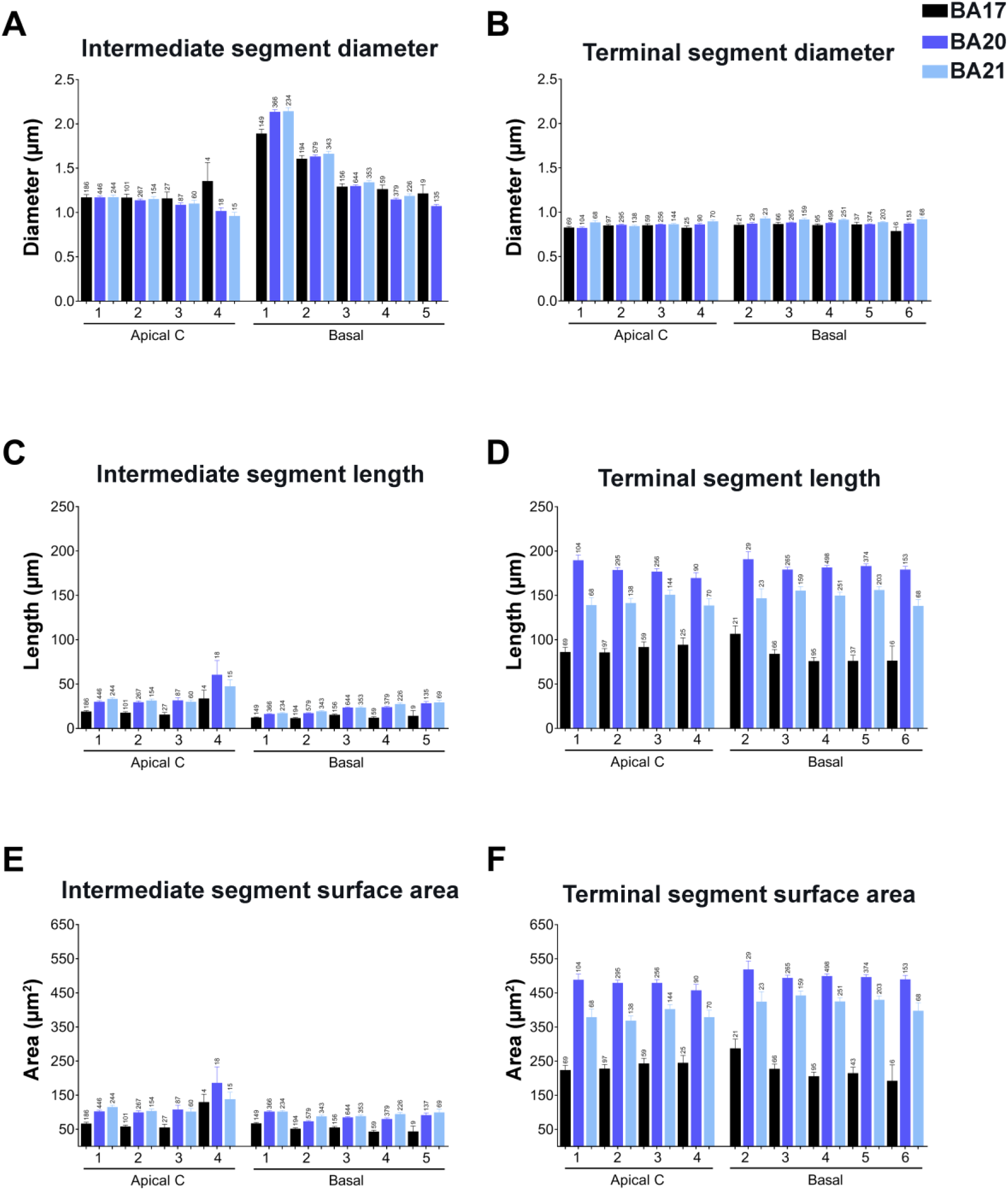
Graphs showing —per branch order— intermediate (**A, C, E**) versus terminal (**D, D, F**) segment diameters (A, B), lengths (C, D) and surface area (E, F) for apical collateral (apical C) and basal dendrites from human BA17 (black), BA20 (dark blue) and BA21 (light blue) cortical areas. Measurements are reported as mean ± SEM. Only dendritic segments that were complete, and thus excluding incomplete endings, were included in this analysis. Additional graphs showing intermediate and terminal segment volumes are shown in Supplementary Figure 5D, E. The statistical significance of the differences is shown in Supplementary Table 5. (**Supplementary Table 3**). The length of the dendritic segments that composed the basal arbors increased in all regions as the branch order increased. Differences were found mainly between branch orders 1–5 in all cortical regions.

Regarding dendritic length, in BA17,—both in apical C and basal intermediate segments— were shorter at most branch orders than those of the temporal areas (**Figure 5C, Supplementary Table 5**). Basal intermediate segments were slightly shorter than apical C intermediate segments in all cortical regions. Terminal segments lengths were longer in the temporal areas than in BA17, and of similar length between branch orders and compartments in each of the cortical regions (**Figure 5D, Supplementary Table 5**).

The values for surface area were slightly higher in apical C intermediate segments compared to basal intermediate segments in both regions, although these values were higher in the temporal cortex (**Figure 5E, Supplementary Table 5**). The surface area of terminal segments was quite similar between branch orders and compartments but showed higher values in the temporal cortex (**Figure 5F, Supplementary Table 5**). The dendritic volume of intermediate segments was higher in the temporal cortex compared to the BA17 region in the different branch orders and compartments (**Supplementary Figure 5D**). The volume of terminal segments was quite similar between branch orders and compartments but showed higher values in the temporal cortex (**Supplementary Figure 5E, Supplementary Table 5**).

#### Axons are initially thicker in the temporal cortex

The axons from all cortical regions analyzed emerged mainly from the soma (98%), with only 2% emerging from the initial portion of a basal dendrite. The axonal diameter was initially thicker in both temporal cortical areas compared to occipital cortex, up to the first 50 μm, and then gradually decreased to reach ∼0.6 μm at a distance of ∼70 μm from the soma and for the remaining distances in all regions (**Figure 3F and Supplementary Table 2**). In the cases in which the axon emerged from the dendrite, its diameter was thinner (∼1.8 μm, 2.35 μm and 1.76 μm for BA17, BA20 and BA21, respectively) and the distance from the soma to the initiation of the axon was ∼3μm in all cortical regions.

In 6, 25 and 15 pyramidal neurons from the BA17, BA20 and BA21, respectively, the axons were of sufficient length to include 1–6 axonal collaterals (**Figure 6A–C**). BA17 main axonal shaft gave off their first collateral at a significantly shorter distance than BA20 and BA21, whereas no significant differences were found for the remaining collaterals (**Figure 6D**). Axonal collaterals showed much greater density than main axonal shafts in all cortical regions (**Figure 6E**). The total mean varicosity density (including the main axonal shaft and collaterals) was 1.15 ±0.16 μm, 1.26 ±0.07μm and 1.21 ±0.12 μm in BA17, BA20 and BA21, respectively. The axonal varicosity density found in the 1–6 collaterals was not significantly different between regions (**Figure 6F**); neither was varicosity interdistance per 10 μm in the 1–6 collaterals (figure not shown). Accordingly, intervaricosity distance was much higher in main axonal shafts than in axonal collaterals (**Figure 6G**). The total mean intervaricosity distance (including the main axonal shaft and collaterals) was 8.78 ±2.86 μm, 11.92 ±2.07 μm and 14.98 ±3.67 μm in BA17, BA20 and BA21, respectively.

**Figure 6.**
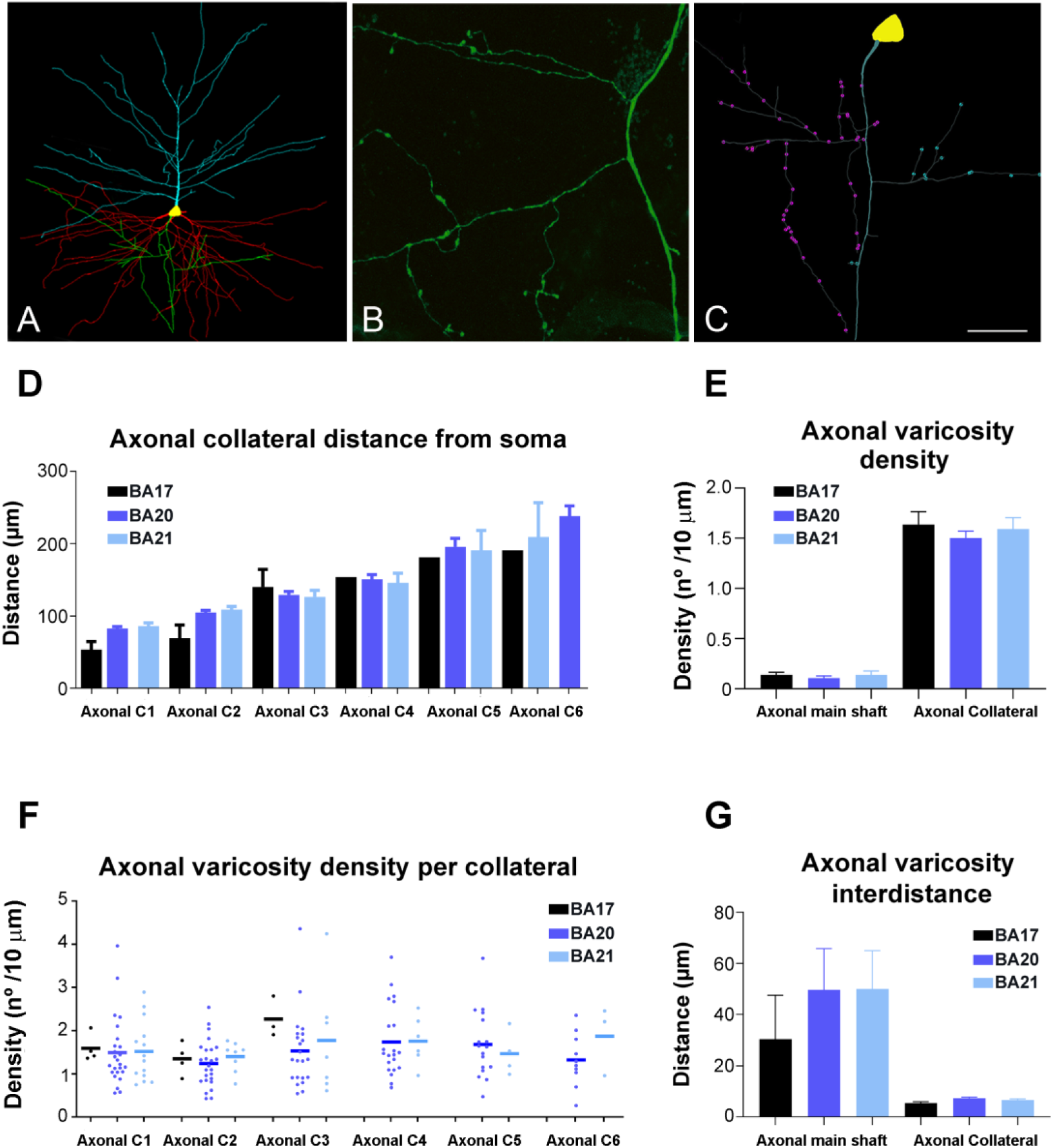
(**A**) 3D pyramidal cell reconstruction (apical: blue; basal: red; axon: green) from BA20. (**B**) High magnification confocal microscopy image from BA20 to illustrate axonal boutons. (**C**) 3D reconstructed axon from the cell shown in A illustrating the main axonal shaft and axonal varicosities from collateral 1 (purple) and collateral 2 (blue). (**D–G**) Graphs showing the mean distance from soma of the axonal collaterals (D), mean axonal varicosity density (E), mean axonal varicosity density per collateral (F) and mean axonal varicosity interdistance (G) in the main axonal shaft and in the axonal collaterals (C1,C2…etc) from human BA17 (black), BA20 (dark blue) and BA21 (light blue) cortical areas. Scale bar (in C) =100 μm in A, 17μm in B and 40μm in C.

### Comparison between neocortical and hippocampal regions

In order to further understand pyramidal cell architecture, we compared the present human neocortical results with those of our previous study in the CA1 hippocampal region. **Figures 7**–**9** display a selection of previously presented graphs including these comparisons (the remaining graphs are shown in **Supplementary Figures 6–9**).

**Figure 7.**
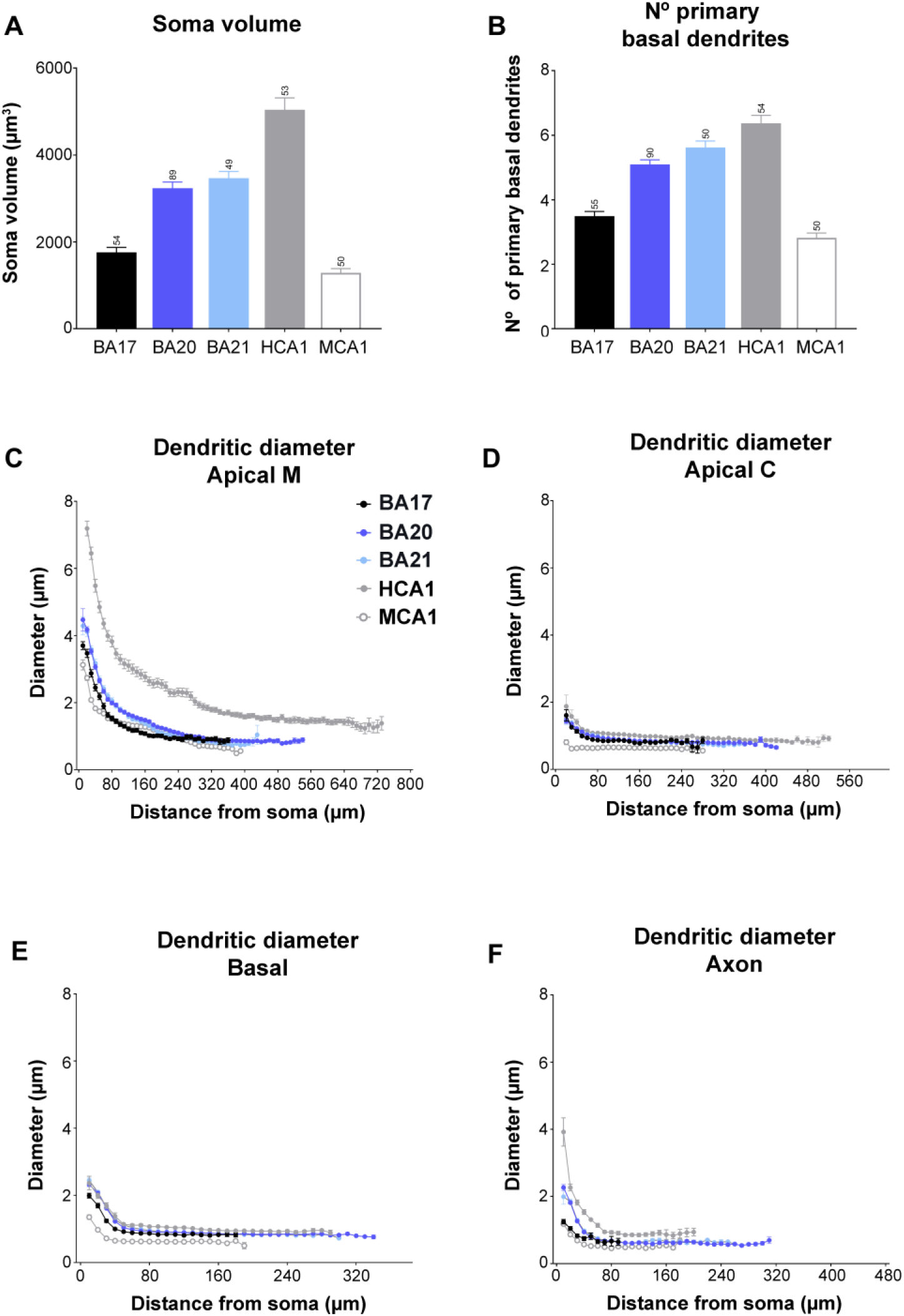
Graphs showing soma volume (**A**), the number of primary basal dendrites (**B**) and diameter distribution as a function of the distance from soma for the main apical dendrites (apical M; **C**), apical collateral dendrites (apical C; **D**), basal dendrites (**E**) and axon (**F**) from human BA17 (black), BA20 (dark blue), BA21 (light blue) cortical areas and human CA1 (grey) and mouse CA1 regions (white). Measurements are reported as mean ± SEM. Additional graphs showing cell body cross-sectional area and surface area are shown in Supplementary Figure 6A, B. The statistical significance of the differences is shown in Supplementary Tables 6, 7.

**Figure 8.**
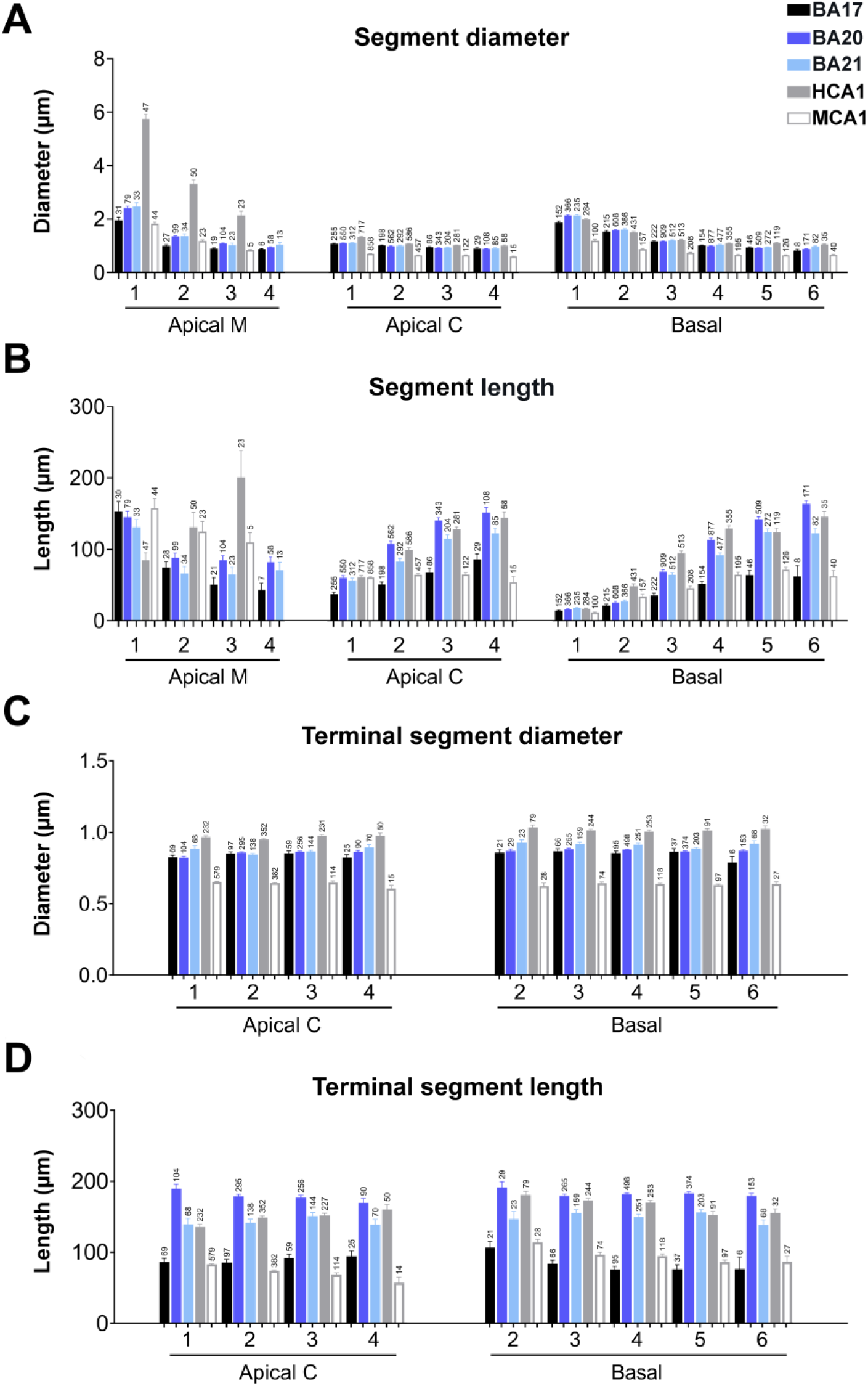
(**A, B**) Graphs showing —per branch order— dendritic segment average diameter (A) and length (B) per dendritic compartment: main apical dendrite (Apical M), apical collateral dendrites (Apical C), and basal arbor (Basal) from human BA17 (black), BA20 (dark blue), BA21 (light blue) cortical areas and human CA1 (grey) and mouse CA1 (white) regions. (**C, D**) Graphs showing —per branch order— terminal segment diameter (C) and length (D) for apical C and basal dendrites from the same regions as in A and B. Measurements are reported as mean ± SEM. Only dendritic segments that were complete, and thus excluding incomplete endings, were included in this analysis. Additional related graphs are shown in Supplementary Figures 7 and 9. The statistical significance of the differences is shown in Supplementary Tables 8, 9.

**Figure 9.**
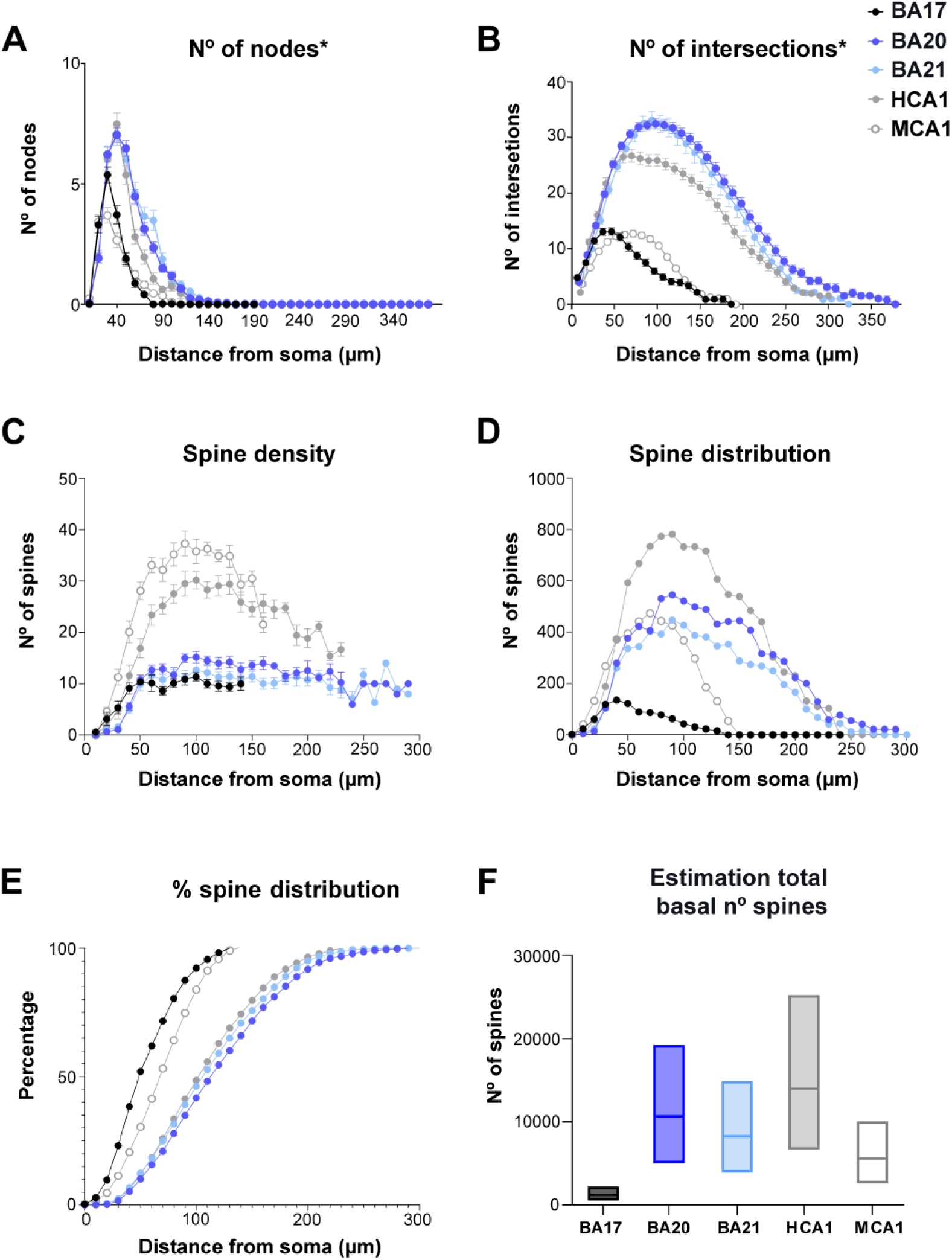
Estimation of total number of spines in basal arbor. Graphs showing the number of basal nodes* (**A**), number of corrected intersections* (**B**), spine density distribution (**C**), estimated number of spines distribution (**D**), cumulative percentage of the estimated number of spines distribution (**E**), range estimation of total basal number of spines (**F**) for the different groups. Note that the percentage of spines distribution is remarkably similar between human temporal and hippocampal regions (E), despite the differences in branching complexity and spine density (B, C). Measurements are reported as mean ± SEM. The statistical significance of the differences is shown in Supplementary Table 10. *Numbers were corrected assuming reconstructed neurons represent 2/3 of the total size (see materials and methods for further details).

#### Neocortical pyramidal neurons have smaller somata, but similar cell size than hippocampal pyramidal neurons

These comparisons showed the size of the pyramidal somata from neocortical areas to be significantly smaller than those of the CA1 human region (5041 ±273.1 μm^3^; **Figure 7A**). Interestingly, no significant differences were found between the soma size of human visual BA17 and mouse CA1 neurons (1288 ±97.52 μm^3^; see **Supplementary Tables 6–9** for statistical comparisons).

#### Neocortical apical and basal dendritic arbors are less complex than those from hippocampal neurons

The structure of the human hippocampal apical M dendrites was more complex than any neocortical region, with a distribution of 22%, 24%, 30%, and 24% for 0 bifurcations, 1 bifurcation, 2 bifurcations, and 3 bifurcations, respectively. CA1 human region (5041 ±273.1 μm^3^; **Figure 7A**). Similar percentages were found between the human BA17 and mouse CA1 neurons. The number of primary basal dendrites (**Figure 7B**) from CA1 pyramidal neurons (6.37±0.25) was the largest (statistically significant when compared to BA17 and BA20; **Supplementary Table 6**). No significant differences were found in the number of primary basal dendrites between human visual area 17 and mouse CA1 neurons (3.04 ±0.15). Additional morphological non-full measurements, showed a proportional distance of 23% of the peak intersection value relative to the extent of the basal arbor in human CA1 and 37% in mouse CA1, whereas in the neocortical regions this percentage was 26-28%.

#### Neocortical apical M dendrites are thinner than hippocampal apical M dendrites

The apical M dendrites of neocortical areas were thinner than those of human CA1 pyramidal neurons at all distances from the soma (**Figure 7C and Supplementary Table 7**). CA1 human apical M diameter stabilized at a higher value (∼1.5 microns) and at a longer distance from the soma (∼400 microns) than neocortical regions. The occipital cortex showed statistically higher values than mouse CA1 neurons from the soma up to 70 microns. The diameter of the apical C dendrites was slightly higher at some distances in human CA1 region (decreasing from ∼1.8 microns to ∼1 microns; **Figure 7D Supplementary Table 7**). The diameter of basal dendrites from temporal cortical areas, per distance from soma (**Figure 7E**), was quite similar to that of human hippocampal neurons, although slightly higher values (∼1 micron) were observed from ∼60 μm from the soma onwards. Mouse CA1 neurons showed smaller values than any human cell, particularly at the distances nearest to the soma (**Supplementary Table 7**).

When dendrites were analyzed per branch order (**Figure 8A**), all neocortical regions showed significantly lower diameter values than human CA1 apical M dendrite at all branch orders. The O1 apical M segments were longer in all neocortical regions compared to the CA1 region (**Figure 8B and Supplementary Table 8**) — and similar to the length of mouse CA1 O1 apical M segments. At higher branch orders, the trend observed in neocortical regions was different from that observed in human hippocampal pyramidal neurons (which increased as branch order increased). In apical M CA1 mouse pyramidal neurons, the trend was similar to that observed in human neocortical regions (length values were similar in branch O1 and larger than human neocortical neurons at higher branch orders (**Supplementary Table 8**). Apical M human CA1 surface area and volume had the greatest values (**Supplementary Figure 7**). Mouse CA1 O1 apical M surface area and volume yielded similar values to human neocortical neurons. At higher branch orders, the trend observed in neocortical regions was different from that observed in human hippocampal pyramidal neurons. In CA1 mouse pyramidal neurons, the trend was similar to that observed in human neocortical regions (values were similar in branch order 1; and larger or similar at higher branch orders in the mouse compared to human neocortical neurons; significant differences are shown in **Supplementary Table 8**).

Apical C dendrites of the CA1 branch order 1 (O1) segment were significantly thicker than in neocortical regions, whereas mouse had thinner segments than any human region. Apical C dendritic length values in the temporal cortex were closer to those of human CA1 apical C dendrites, whereas human occipital cortex dendrites yielded values that were closer to those of mouse CA1 (see **Figure 8B and Supplementary Table 8** for statistical comparisons). Apical C dendritic segment surface area values in the temporal cortex were similar to those in the human CA1 region (particularly those of area 20) and all human cortical areas had larger values than in mouse neurons (see **Supplementary Figure 7 and Supplementary Table 8** for statistical comparisons).

The values for neocortical basal dendritic diameters per branch order (**Figure 8A**) were similar to those in human neocortical pyramidal neurons. Mouse CA1 basal dendrite values were lower in all cases (See **Supplementary Table 8** for statistical comparisons).

Temporal basal dendrites were shorter than those in the hippocampal region (**Figure 8B**) in branch O2–4, whereas there were fewer differences between the remaining orders (see **Supplementary Table 8**). Human occipital cortex dendrites had similar lengths to mouse CA1 basal dendrites. CA1 basal dendrites had higher surface area and volume values (**Supplementary Figure 7**) than temporal cortex from branch O2 and the remaining orders.

Human occipital cortex dendrites had values that were either similar to or higher than those of the mouse CA1 neurons (see **Supplementary Table 8** for statistical comparisons).

When dendrites were further analyzed per intermediate/terminal segments, human temporal and occipital cortex yielded relatively similar diameter values to those of human CA1 neurons, with larger diameters than mouse neurons, both in intermediate and terminal segments (**Figure 8C and Supplementary Figure 9**). However, segment lengths were shorter in BA17 and similar to mouse CA1 neurons (**Figure 8C, D** and **Supplementary Figure 9**). The surface area and volume of intermediate and terminal segments were also lower in mouse CA1 neurons (**Supplementary Figure 9**). That is, all intermediate and terminal segments were thicker and had larger surface areas and volumes in human neurons than in mouse neurons.

#### Neocortical axons are thinner than hippocampal axons

Neocortical regions showed a similar decreasing trend to that of CA1 neuron axonal diameters, although the CA1 neurons were thickest at all distances (from 3.92 ±0.42 μm to ∼0.9 μm; **Figure 7F**). The distance from where the diameter showed similar values was similar to that of neocortical regions (∼70 microns from soma). Mouse CA1 neurons were slightly thinner than BA17 human neurons, but no significant differences were found.

The distances from where the collaterals emerged were longer in CA1 human neurons. The axonal varicosity density was slightly higher in CA1 neurons (1.74 ±0.23, 1.06 ±0.02, 1.39 ±0.08, and 1.31 ±0.03 varicosities per 10 μm in the first, second, third, and fourth collaterals, respectively), compared to neocortical neurons. Consequently, the human CA1 axonal intervaricosity distance was smaller (6.53 ±1.26, 3.45 ±0.06, 3.75 ±0.17, and 3.77 ±0.68 μm in the first, second, third, and fourth collaterals, respectively) than in neocortical regions.

### Additional morphological features

Additional morphological non-full measurements, such as convex hull 2D/3D, total number of nodes, intersections, length, etc., are displayed in **Figure 9A, B and Supplementary Figures 10–12**. Although not full numbers, these measurements showed, for example, that cells were smaller in occipital areas compared to temporal areas and that the peak number of nodes and intersections in the BA17 basal arbors were also lower (**Figure 9A, B; Supplementary Figure 12A, B**). Also, BA17 showed peak intersection values closer to the soma compared to BA20 and 21 but at a quite similar proportional distance relative to the extent of the basal arbor (with distances from the soma of 26%, 28% and 28% for BA17, BA20 and BA21, respectively; see also **Figure 9A, B and Table 2**). These estimations were 23% in human CA1 and 37% in mouse CA1.

### Dendritic spines

In order to estimate the total number of spines that a basal arbor would contain, the numbers of dendritic intersections were corrected (assuming that the basal arbors in coronal sections represent two thirds of the total basal arbor (Krimer et al., 1997). **Figure 9A, B** shows these corrected basal number of nodes and intersections. The spine density analysis (**Figure 9C**) showed that hippocampal neurons (particularly in the mouse) have the highest spine density values, whereas human BA17 neurons had the lowest values (see **Supplementary Table 10** for statistical comparisons). The spine density distribution (**Figure 9D**), as an estimation of the number of spines, combining the results from Sholl analysis (**Figure 9B**) and those of spine density (**Figure 9C**) showed large differences between the groups. Human CA1 neurons showed the greatest number of spines, whereas BA17 neurons had the lowest values. Interestingly, hippocampal neurons in both species showed a steeper descending curve than neocortical neurons. Moreover, the normalized cumulative distribution (**Figure 9E**) showed a similar distribution in human temporal and CA1 region, whereas visual and mouse CA1 neurons followed different distributions. **Figure 9F** shows the estimated total number of spines contained within the basal arbor. These estimations showed that human CA1 neurons present the highest values, followed by human temporal neurons, and then mouse CA1 neurons, with BA17 neurons displaying the lowest values.

### Correlations of variables

We then examined the potential correlation between variables by performing several correlations (**Figure 10 and Supplementary Figures 13–17**; see **Supplementary Table 11** for statistical comparisons). We found strong correlations between M apical diameter-soma size-axonal diameter (**Figure 10**) such that the larger the apical M diameter, the larger the axonal diameter and the soma size. Also, the size and complexity of dendrites were related to the thickness of the layer in which they lie and the distance to the pia (see **Supplementary Figure 17 and Supplementary Table 11** for statistical comparisons).

**Figure 10.**
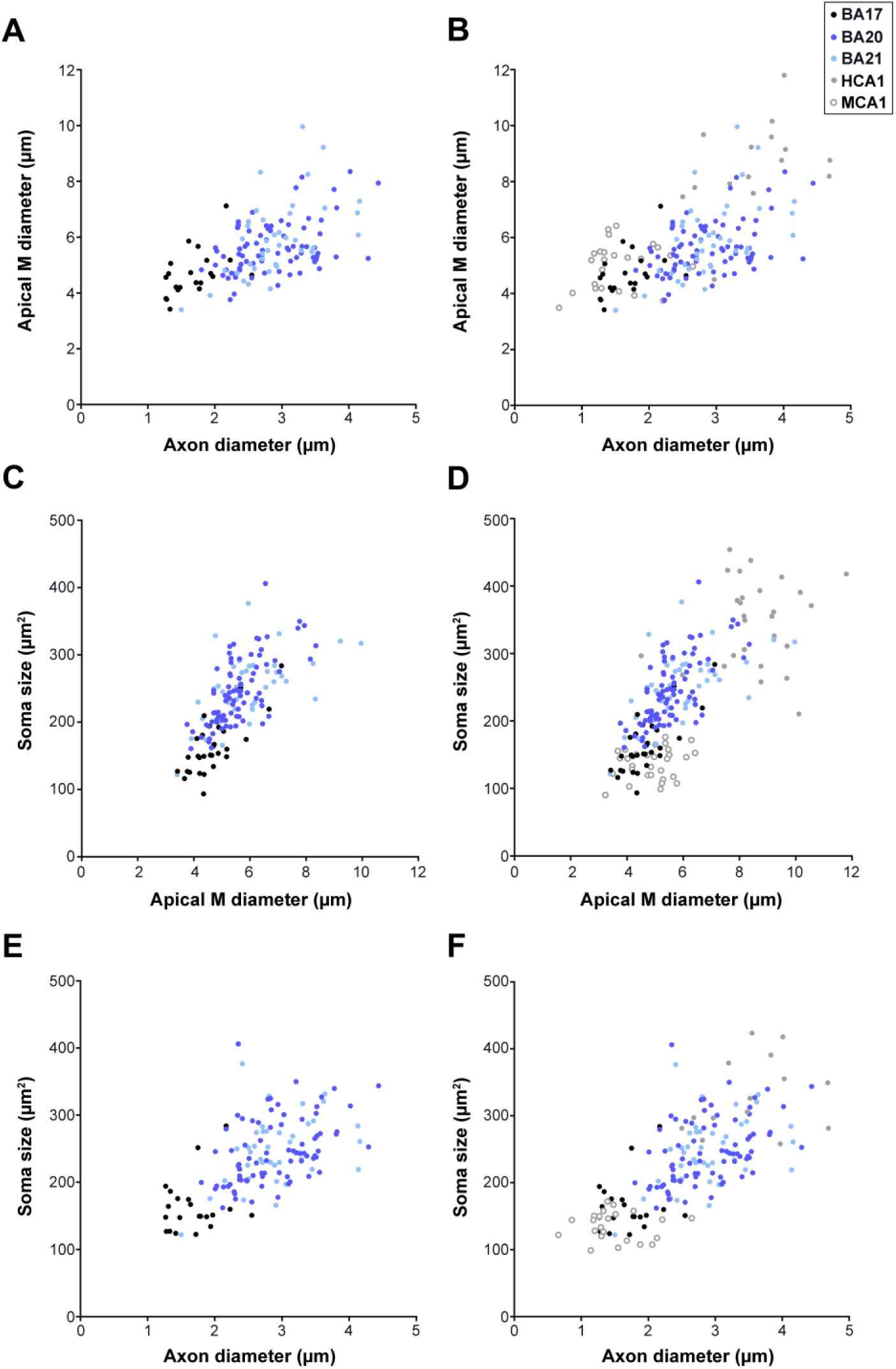
Correlation of variables. Correlation analyses between various morphological parameters analyzed. Each point represents the values obtained in one cell from human BA17 (black), BA20 (dark blue), BA21 (light blue) cortical areas (left column). Additional correlations, including human CA1 (grey) and mouse CA1 (open circles) regions, are shown (right column). Significant correlations were classified as weak [Spearman’s rho (r) value lower than 0.40], moderate (0.4<r<0.7), and strong (r>0.7). The statistical significance of the differences is shown in Supplementary Table 11. See also Supplementary Figures 13–17 for further morphological feature relations.

## Discussion

The primary finding in the current study is that the structure of distinct dendritic compartments of pyramidal neurons within human occipital and temporal cortex reveals specific shared and distinct morphological characteristics across various areas and species. These insights offer fundamental understanding regarding their morphological specializations, highlighting significant variations in the processing of information within the cortex. Specifically, the observations were as follows: 1) Human pyramidal neurons from temporal association cortex exhibit larger sizes and more complex apical and basal dendritic patterns. They also possess a greater number of spines and thicker axons in comparison to those in the primary visual cortex; 2) Dendritic and axonal thickness, as well as the lengths of dendritic segments, stand out as crucial, previously unrelated morphological characteristics that distinguish human temporal neurons from primary visual neurons; 3) The size and dendritic complexity of pyramidal neurons correlate with the thickness and depth of the layer in which they are found. Notably, greater apical M diameters correspond to larger axonal diameters and soma sizes; 4) A prominent distinction between temporal association cortex neurons and human hippocampal CA1 pyramidal neurons lies in the thickness and complexity of apical dendrites; 5) Human temporal neurons share more similarities with CA1 human neurons than with neurons from the human BA17 cortex. Conversely, the relatively smaller and simpler human BA17 pyramidal neurons exhibit more similarities with CA1 mouse pyramidal neurons than with neurons from the human temporal cortex; 6) The increased thickness of dendrites emerges as a consistent morphological trait among all dendritic compartments in the human neurons analyzed, setting them apart from mouse neurons; and, finally, 7) Certain shared patterns of organization exist among different regions and species.

### Methodological Considerations

Due to technical limitations, 3D reconstructions of neurons do not include complete basal and apical arbors (see methods for further details). Indeed these limitations may not apply equally to the areas: the larger cells in temporal cortex may be more likely truncated, or be truncated to a larger extent, in comparison with occipital cells. Similarly, the spatial extent on axonal arborization is limited since only the proximal axonal extent were included within the sections. Therefore, we focused on morphological variables that do not depend on the entirety of the reconstructed cell (soma area and volume, segment diameter, segment length, segment surface area and segment volume). These measurements can be consistently compared. However, some results from variables that do depend on the entirety of the cell —primarily presented in the supplementary material (2D and 3D convex hull, total number of dendrites, nodes, intersections, total dendritic length, surface area and volume)— should be interpreted with these technical limitations in mind.

Furthermore, while we managed to reconstruct a relatively large number of neurons and individuals (n=7), it is important to note that not all cases included every region, and the number of neurons examined varied across different regions. This variability can be attributed to technical limitations and challenges encountered in obtaining human brain tissue with the requisite quality of fixation for these experiments. Moreover, it is well-established that the structure of the dendritic arbor varies with the age of the individuals examined (for example, see Petanjek et al., 2019). In the present study, the age range of the individuals examined spans from 40 to 85 years. Nonetheless, the age of six out of the seven cases studied falls within middle adulthood (ranging from 40 to 66 years old). In spite of these limitations, the present study constitutes a significant advancement in the characterization of human brain microorganization. However, it remains essential to validate these findings with a larger cohort of individuals and a broader range of cortical regions.

### Main differences between human occipital and temporal neurons

#### Increased potential for compartmentalization in temporal neurons

Neurons from temporal cortex (BA20 and BA21) exhibited larger cell size and more complex branching complexity patterns compared to those in human primary visual cortex (BA17). Similar results had previously been observed in basal dendrites of human pyramidal neurons from the secondary visual cortex (BA18) when compared to basal dendrites of pyramidal neurons from the BA20 temporal association cortex (Elston et al., 2001). In the present study, we expanded upon these previous findings by examining not only the basal arbor but also the apical arbor (including the apical M and C dendrites), the soma, the local axonal arbors, and additional detailed features such as dendritic diameter and dendritic segment lengths. **Figure 11 and Table 2** provides a summary of the main distinctive features between human occipital and temporal neurons.

**Figure 11.**
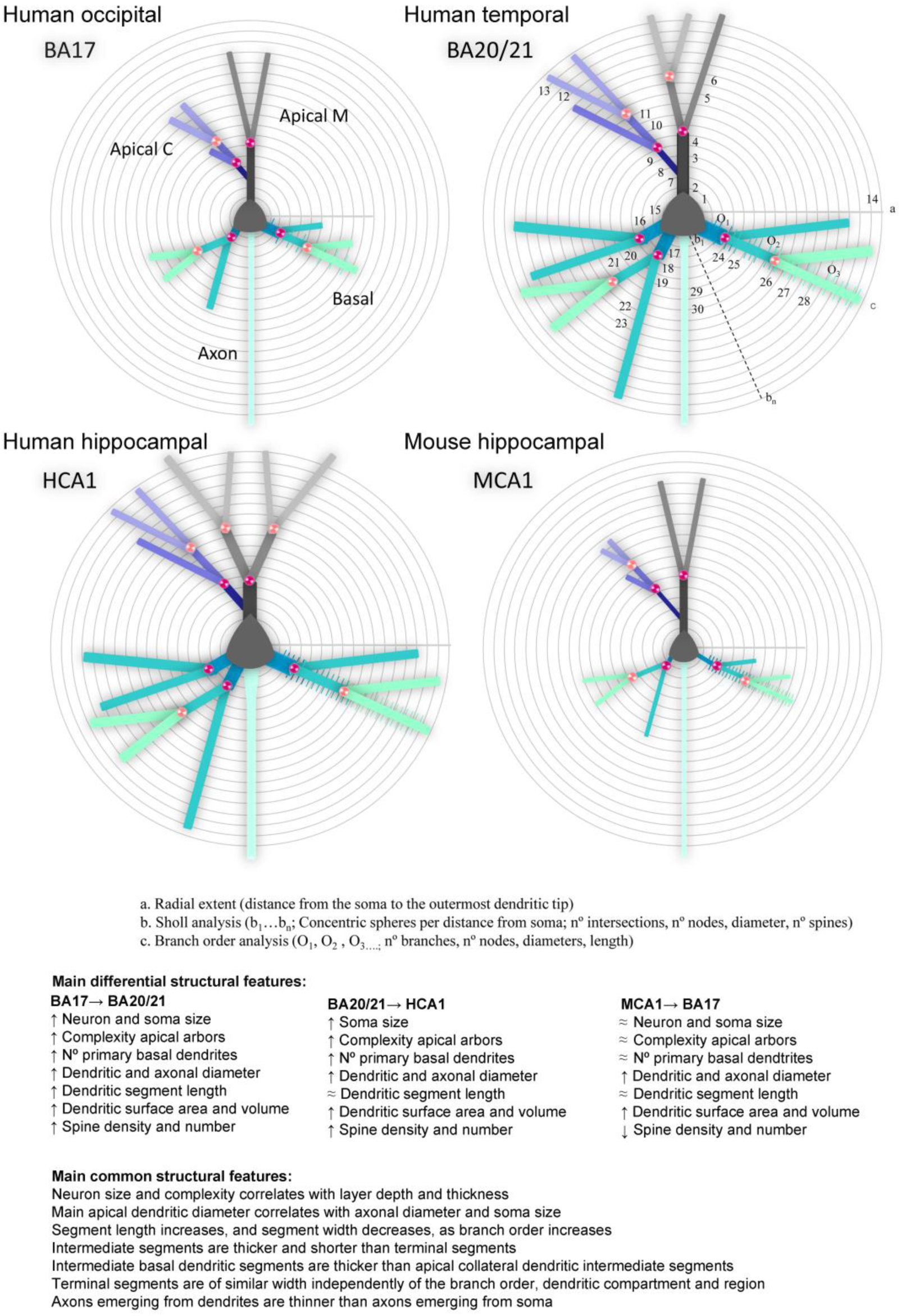
Schematic drawing of the main morphological differences shown in Table 2 (variables n° 1-30), including (a), radial extent; (b), Sholl analysis; and (c), branch order analysis variables, in apical main (M), apical collateral (C), basal and axonal compartments, from human primary visual (BA17), human associative temporal (BA20/21) and human (HCA1) and mouse (MCA1) hippocampal CA1 neurons. Symbols: X→Y: from X to Y; ↑: increase; ↓: decrease; **≈**: similar.

The apical arbor was also found to be larger, displaying a more complex and diverse pattern of arborization in association temporal areas compared to BA17. Additionally, soma size was noticeably larger in temporal regions. The diameters of the apical M and basal dendrites were greater than those in the visual cortex, particularly at the nearest distances from the soma (corresponding to branch orders O1 and O2). The dendritic segments, both in apical C and basal dendrites, were longer at greater distances from the soma. Consequently, dendritic surface area and volume were greater in temporal neurons compared to BA17 neurons across most compartments and dendritic orders. Functionally, these variations in size and branching complexity patterns are linked to the sampling strategies and integration of inputs (e.g London and Hausser, 2005), which contribute to distinct forms of processing within the dendritic tree before input potentials arrive at the soma. Consistent with previous studies (e.g., Lund et al., 1993; Malach 1994; Elston et al., 1999; Jacobs et al., 2001; Elston, 2003), these findings emphasize the increased potential for compartmentalization in human temporal areas when compared to the human primary visual region. This higher apical branching complexity in temporal regions, relative to BA17, further amplifies the extent to which integration of inputs are significantly compartmentalized within the dendritic arbors. As a result, this study reveals that both apical and basal compartments consist of highly branched structures and highlights that dendritic diameters and segment lengths are pivotal features for distinguishing human neurons across various cortical regions.

The larger size and relatively high branching complexity of temporal pyramidal neurons, coupled with a notably high dendritic spine density, result in this region having a much higher number of spines compared to the relatively smaller BA17 neurons, which exhibit low branching complexity and lower dendritic spine density. Consequently, both pyramidal neurons’ apical and basal arbors in the temporal cortex can sample a larger number of inputs, compartmentalize such inputs to a greater degree, and integrate these inputs differently from those in V1 cortex. These results are in line with previous comparative studies on mouse and monkey cortex (e.g., Gilman et al., 2017; Luebke, 2017).

Regarding the axon, neurons in the temporal cortex exhibit a greater diameter, particularly at shorter distances from the soma. Additionally, axonal collaterals were observed to emerge at greater distances from the soma than in the occipital cortex, potentially influencing both excitability and the propagation of action potentials.

#### Correlation between macroscale and cell complexity

Correlation analyses indicated that 1) the size and complexity of pyramidal neurons correlate with layer depth and thickness and 2) the apical M diameter correlates with axonal diameter and soma size. Regarding the size and complexity of dendrites they were correlated with the distance of the soma to the pia, which is in line with previous findings (Deitcher et al., 2017). Additionally, the thicker layer 3 of BA20 and BA21, in comparison to BA17, was linked to larger cell size and greater complexity. This finding is consistent with previous research on the temporal cortex, which demonstrated increased thickness accompanied by larger dendrites and cell body size in layers 2/3 of pyramidal neurons (Heyer et al., 2021). As a result, it is likely that association temporal areas have greater connectivity and display a distinct organization compared to the visual cortex. These results are in line with reports suggesting a correlation between macroscale connectivity and the complexity of layer 3 pyramidal dendrites across various human cortical regions (e.g., van den Heuvel et al. 2015, 2016). Moreover, positive correlations were found between the soma size and the apical M diameter as well as the axonal diameter. Consequently, these results highlight the greater size and complexity of temporal pyramidal neurons in comparison to occipital neurons, including both dendrites (apical and basal) and axons. These characteristics are correlated with the thickness and depth of the layer in which the neurons are located; as the apical M diameter increases, so does the axonal diameter and the soma size. These observations carry distinct computational implications for different cortical regions.

### Main differences and similarities between neocortical and hippocampal regions

#### Apical dendrite, soma and axonal thickness and complexity distinguish human temporal neurons from human hippocampal CA1 neurons

The primary distinctions between human neocortical and hippocampal regions were the larger apical M CA1 dendritic diameter, more complex and varied patterns of apical dendritic branching, and a greater soma size and axonal diameter in human hippocampal neurons compared to neocortical neurons. Additionally, the peak basal dendritic branching complexity was relatively closer to the soma in human CA1, unlike in neocortical areas where it was relatively farther away (in the mouse CA1, it was relatively more distant). Another important difference between human temporal and hippocampal CA1 pyramidal cells is regarding the origin of their axons, as the site of origin has a significant impact on the electrical properties of neurons (for a review see Kole et al., 2018). Axons originating from neocortical regions mainly emerged from the soma (98%), which is in line with previous studies on the origin of axons of pyramidal neurons in monkeys and humans (Whale et al., 2022). However, in CA1, they emerged either from the soma (∼70%) or from the initial segment of a basal dendrite (∼30%). Furthermore, axonal collaterals emerged at greater distances from the soma, and the density of axonal varicosities was slightly higher compared to neocortical regions (which were similar to each other). These axonal distinctions underscore that the structure of pyramidal neurons varies between regions and species — not only in terms of size, but also with regard to the architectural design of their cellular components.

#### Human temporal neurons share more similarities with CA1 human neurons than with neurons from the primary visual cortex

These key attributes are outlined in **Figure 11**, highlighting characteristics like large dendritic trees, relatively complex dendritic branching, and a significant number of spines. Conversely, the smaller and relatively simpler BA17 neurons shared more similarities with CA1 mouse neurons than with neurons from the human temporal cortex. Notably, these consisted of smaller dendritic trees and comparatively simpler patterns of dendritic branching complexity.

The comparison of estimations for the total number of basal spines revealed that human CA1 large neurons, characterized by relatively high branching complexity (albeit lower than neocortical neurons) and very high dendritic spine density, led to this region having the highest number of spines. Similarly, mouse CA1 field also exhibited very high spine density— but lower branching complexity than in neocortex (Ballesteros-Yañez et al., 2010), resulting in a relatively high estimated number of spines within this species.

It is interesting to note that mouse CA1 neurons resulted in higher estimations of dendritic spines within their basal arbors compared to human primary visual neurons. Consequently, human CA1 pyramidal neurons could potentially sample the largest number of inputs, compartmentalize them to a greater degree, and integrate these inputs differently from neurons analyzed to date in neocortical regions.

#### Human dendrites are thicker than mouse dendrites

It was observed that both neocortical (occipital and temporal) and hippocampal human neurons exhibited specific shared structural patterns that were not found in mice. For instance, the thickness of dendritic segments was consistently greater in all dendritic compartments of the human neurons analyzed in comparison with mouse neurons. This finding is in line with previous research on the mouse somatosensory cortex (e.g., Alpar et al., 2006; Merino-Serrais et al., 2023), which also reported smaller dendritic diameters compared to the human pyramidal neurons analyzed in the present study.

Additionally, our findings revealed that intermediate basal segments were thicker than collateral segments in all of the human neurons analyzed, whereas in mice, basal segments were thinner and more akin to collateral dendrites. Similarly, terminal segments in mice were thinner than those found in any human cell. This suggests that dendritic diameter likely plays a crucial role in distinguishing between human and mouse neurons in this analysis.

### Common dendritic organizational patterns across regions and species

Several morphological variables exhibited similar characteristics and/or values in both the occipital and temporal regions: 1) Across all cortical areas and compartments dendritic diameter values decreased as the branch order increased. 2) apical C diameters displayed similar measurements in all three areas analyzed. 3) the length of dendritic segments, in all compartments, increased with higher branch orders; 4) the surface area of initial dendritic segments was comparable between the regions; 5) intermediate segments (both in apical C and basal dendrites) were thicker and shorter compared to terminal segments; and intermediate basal segments were thicker than collateral segments — while terminal segments (both apical C and basal) exhibited similar widths. These features were also found in previously published CA1 human and mouse neurons. Thus, through detailed analyses of these features, it is revealed that they represent specific morphological parameters conserved across regions and species. They likely reflect a general trend in the structural organization and design of pyramidal neurons.

### Conclusions

The present results unveil new detailed and significant morphological features, highlighting distinct regional and species-specific specializations that contribute to varied computational implications. The principal differences between human occipital and temporal regions include the larger size, dendritic complexity, length of dendritic distal segments, and axonal diameter of human temporal neurons compared to those in the occipital region. Likewise, the key disparities between human neocortical and hippocampal regions include the thicker apical CA1 arbors, more intricate apical complexity patterns, and larger axonal diameter in human hippocampal neurons relative to neocortical neurons. Moreover, a notable distinction between human and mouse neurons lies in the larger dendritic diameter of human neurons in comparison with mouse neurons. In light of these findings, we propose that the increase in pyramidal cell complexity during cortical expansion can be understood as occurring through the following steps: (1) an increase in dendritic diameter, followed by the further enhancement of apical main and basal dendrites, along with an increase in axonal diameter; (2) An enlargement in neuron size, involving: a) Extension of distal dendritic segment lengths; b) Augmentation in dendritic complexity (e.g., number of nodes and dendrites); and c) Increase in the number of dendritic spines. These morphological features likely play a pivotal role in shaping distinct regional and species-specific cell specializations, systematically contributing to the diverse cortical processing capacities within each region and species.

## ACKNOWLEDGMENTS

We would like to thank Debora Cano, Lorena Valdes, Carmen Alvarez, Miriam Marin, and Ana García for their technical assistance and Nick Guthrie for his helpful editorial assistance. This work was supported by grants from the following entities: Grant PID2021-127924NB-I00 funded by MCIN/AEI/10.13039/501100011033; Centro de Investigación en Red sobre Enfermedades Neurodegenerativas (CIBERNED, CB06/05/0066); CSIC Interdisciplinary Thematic Platform (PTI) Cajal Blue Brain (PTI-BLUEBRAIN; Spain); and the European Union’s Horizon 2020 Framework Programme for Research and Innovation under Specific grant agreement No. 785907 (Human Brain Project SGA2) and Specific grant agreement No. 945539 (Human Brain Project SGA3).

## SUPPLEMENTARY FIGURES AND TABLES

**Supplementary Figure 1.**
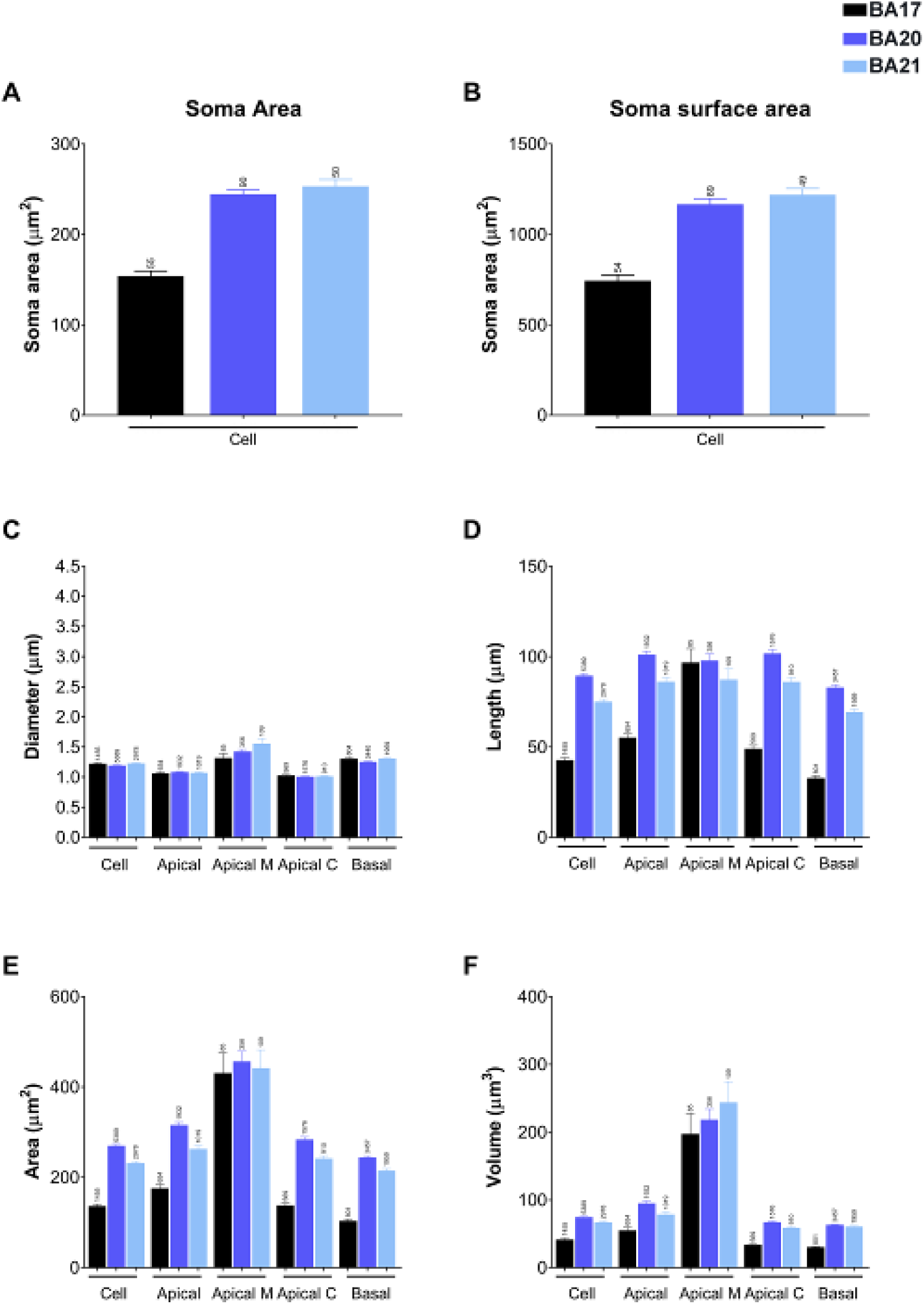
Graphs showing cell body cross-sectional area (**A**), soma surface area (**B**), dendritic segment average diameter (**C**), segment length (**D**), segment surface area (**E**), and segment volume (**F**), expressed per cell and per dendritic compartment: apical arbor (including main apical dendrite and apical collateral dendrites together); main apical dendrite alone (Apical M); apical collateral dendrites alone (Apical C); and basal dendritic arbor. Measurements are reported as mean ± SEM. Only dendritic segments that were complete, and thus excluding incomplete endings, were included in this analysis. The statistical significance of the differences is shown in Supplementary Table 1.

**Supplementary Figure 2.**
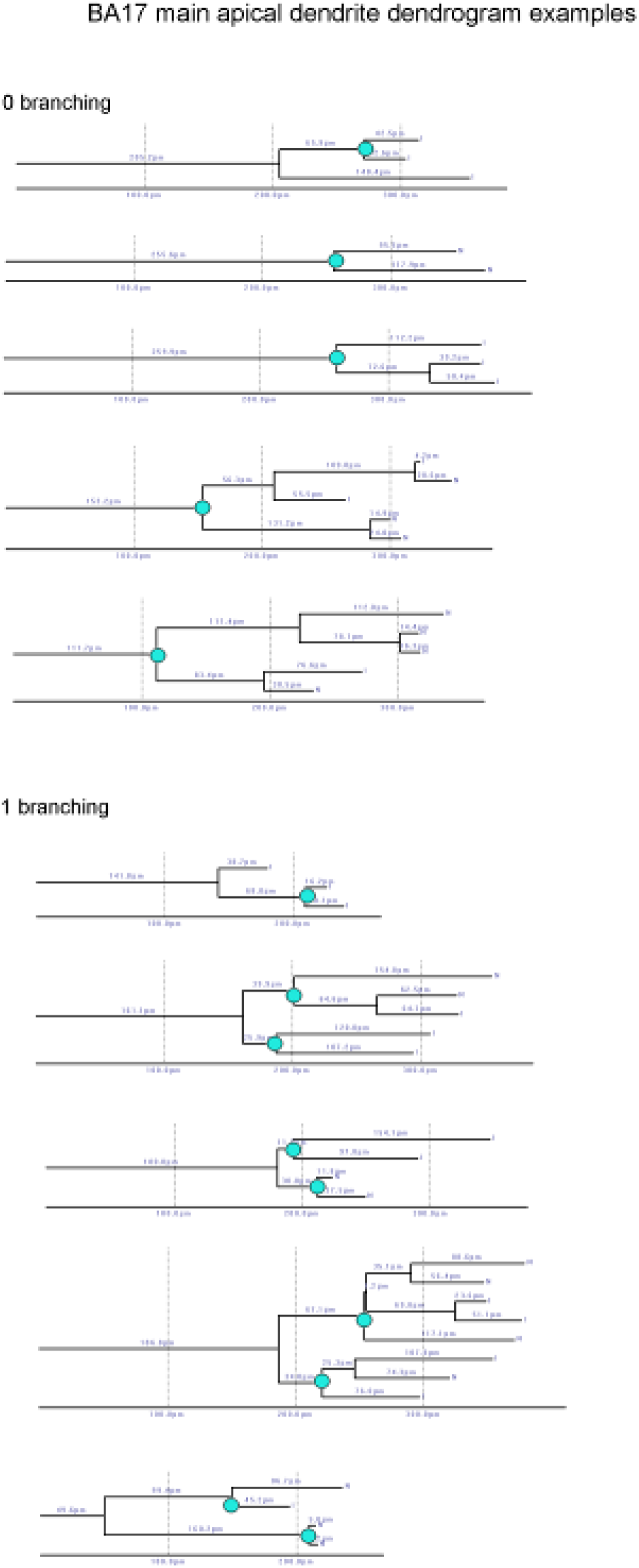
Dendrogram examples of main apical dendrites that had 0 bifurcations and 1 bifurcation (within the first 200 µm) from the 3D reconstructed dataset of human BA17 pyramidal neurons. Two and three bifurcations were absent.

**Supplementary Figure 3.**
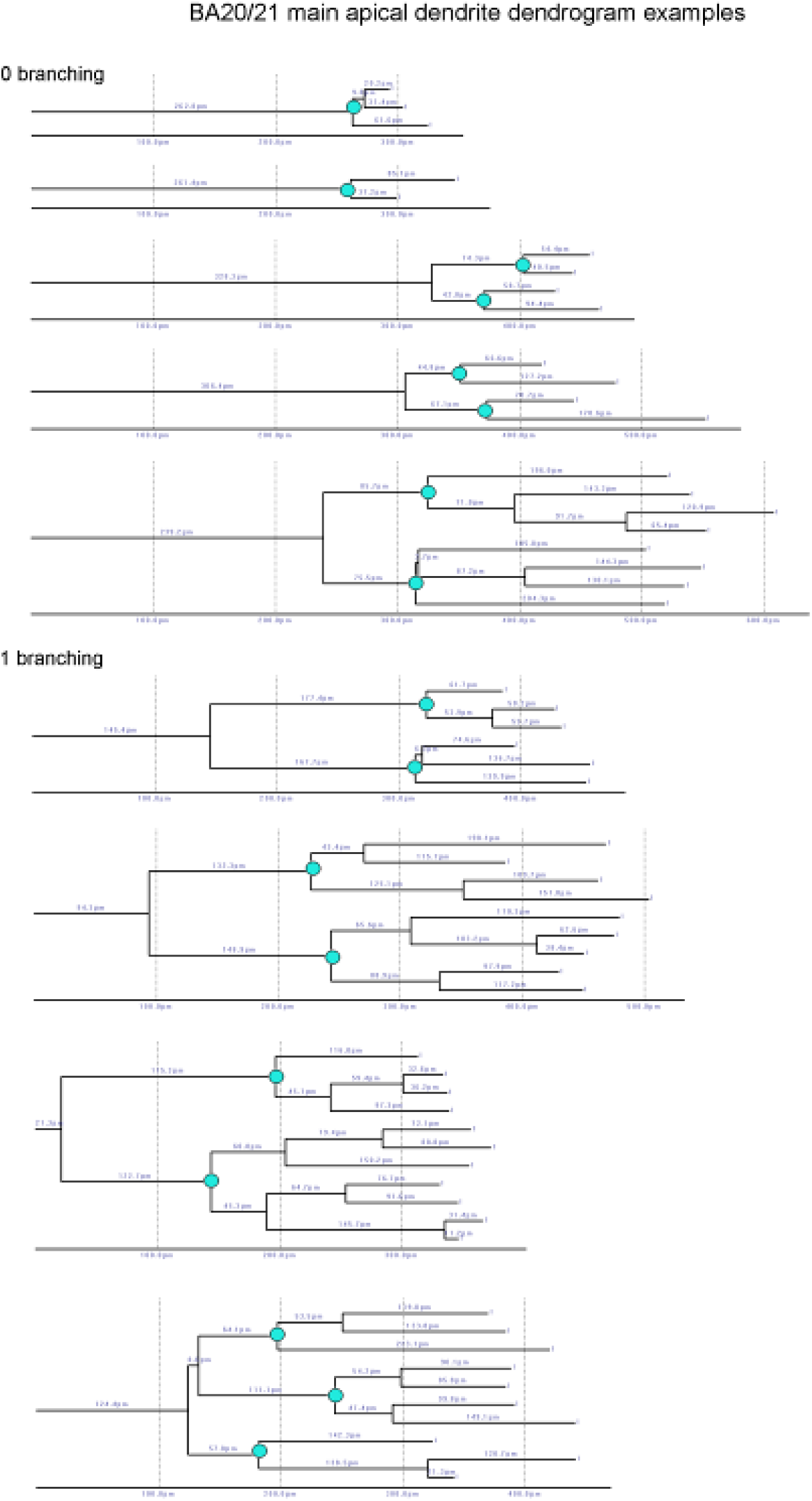
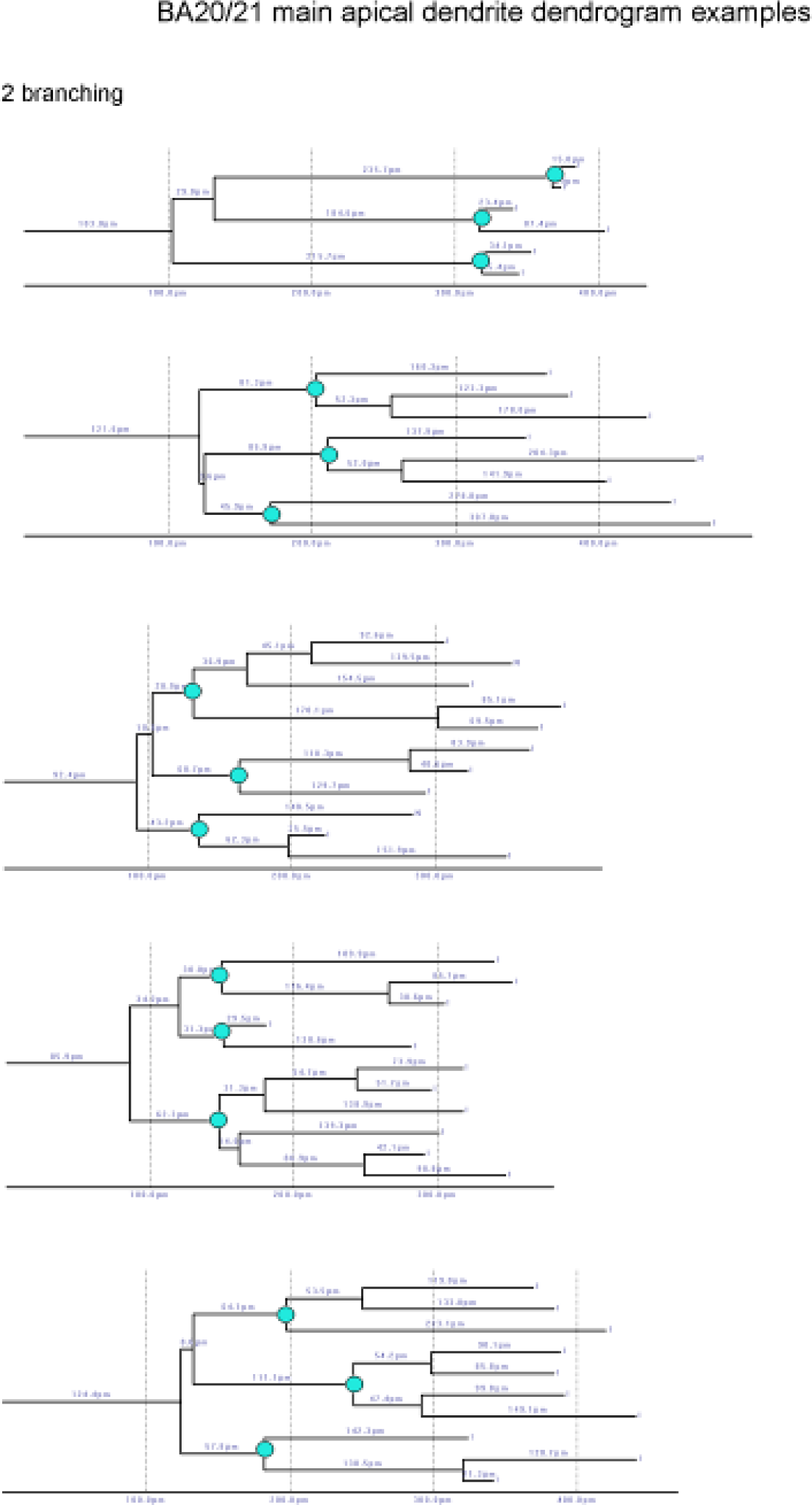
Dendrogram examples of main apical dendrites that had 0 bifurcations, 1 bifurcation and 2 bifurcations (within the first 200 µm) from the 3D reconstructed dataset of human BA20 and BA21 pyramidal neurons.

**Supplementary Figure 4.**
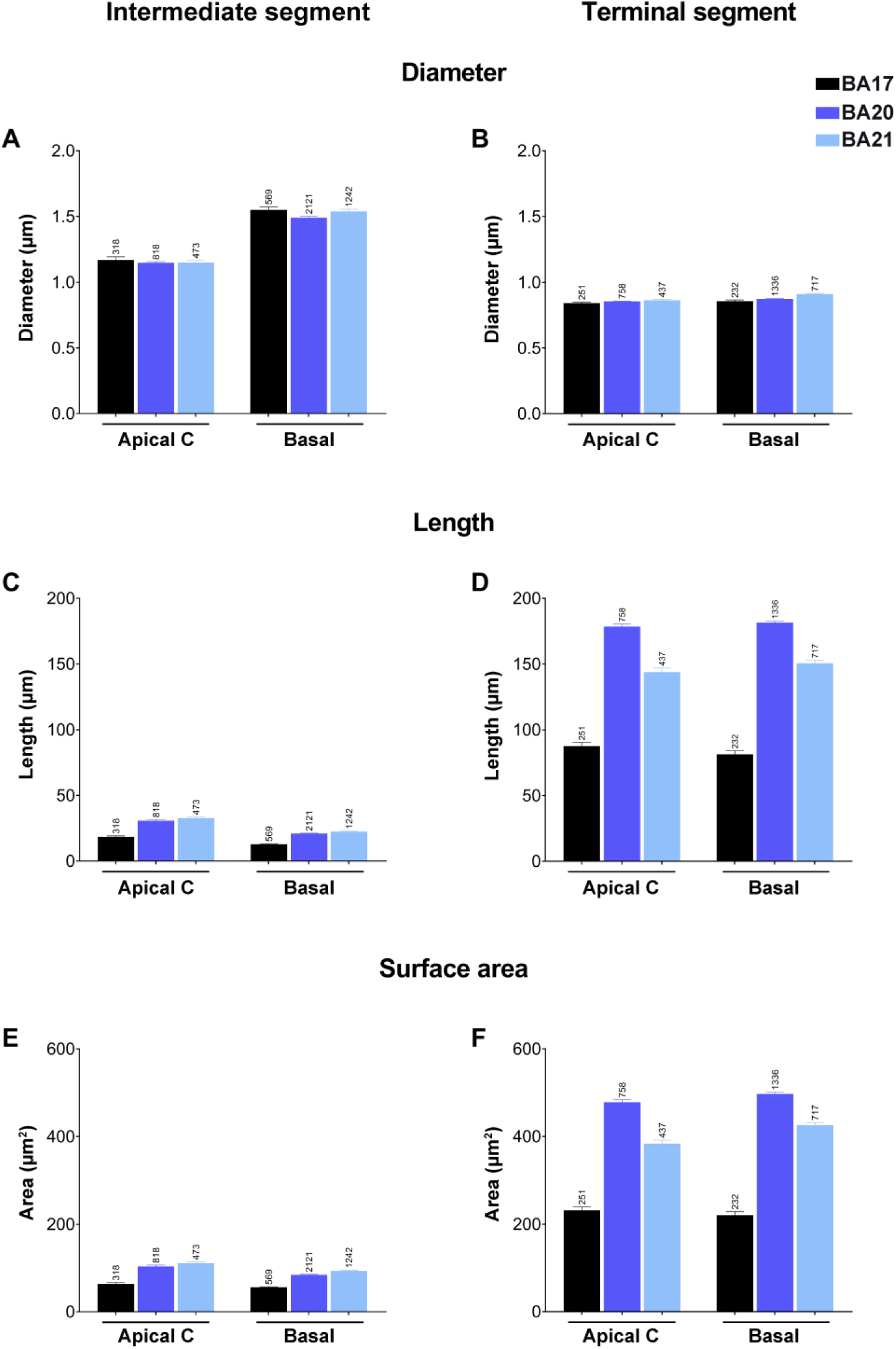
Graphs showing intermediate (**A, C, E**) versus terminal segment (**B, D, F**) diameters (A, B), lengths (C, D) and surface area (E, F) for apical collateral (apical C) and basal dendrites from human BA17 (black), BA20 (dark blue) and BA21 (light blue) cortical areas. Measurements are reported as mean ± SEM. Only dendritic segments that were complete, and thus excluding incomplete endings, were included in this analysis. Additional graphs showing intermediate and terminal segment volume are shown in Supplementary Figure 5B, C. The statistical significance of the differences is shown in Supplementary Table 4.

**Supplementary Figure 5.**
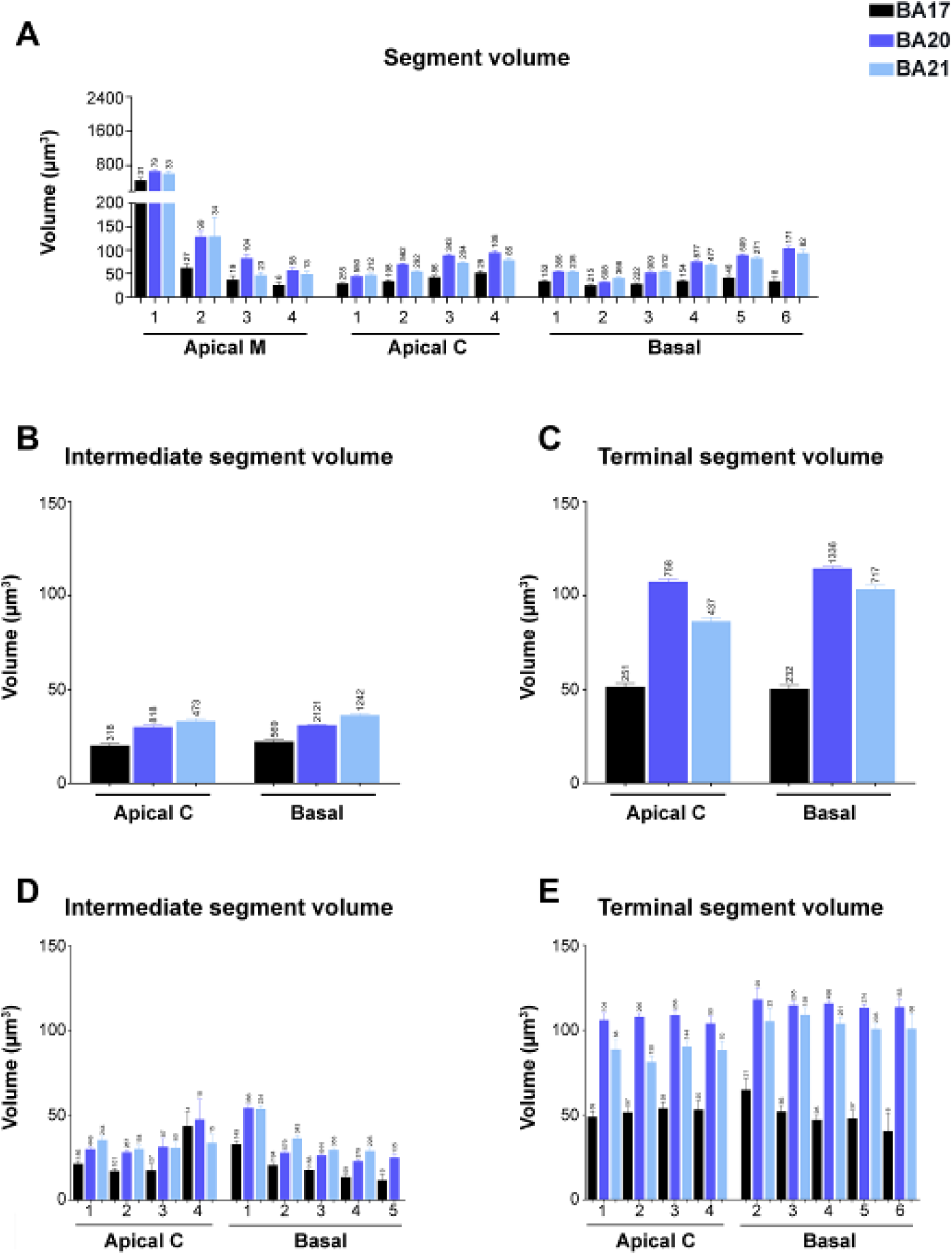
(**A**) Graph showing dendritic segment volume, expressed per branch order (1, 2, 3, etc.) and per dendritic compartment: main apical dendrite (Apical M), apical collateral dendrites (Apical C), and basal arbor (Basal) from human BA17 (black), BA20 (dark blue) and BA21 (light blue) cortical areas. (**B**, **C**) Graphs showing intermediate (B) versus terminal segment (C) volume for apical C and basal dendrites from the same 3 cortical areas as in A. (**D, E**) Graphs showing —per branch order— intermediate (D) versus terminal (E) segment volume for the same compartments and cortical areas as in the previous graphs. Measurements are reported as mean ± SEM. Only dendritic segments that were complete, and thus excluding incomplete endings, were included in this analysis. The statistical significance of the differences is shown in Supplementary Tables 3, 4.

**Supplementary Figure 6.**
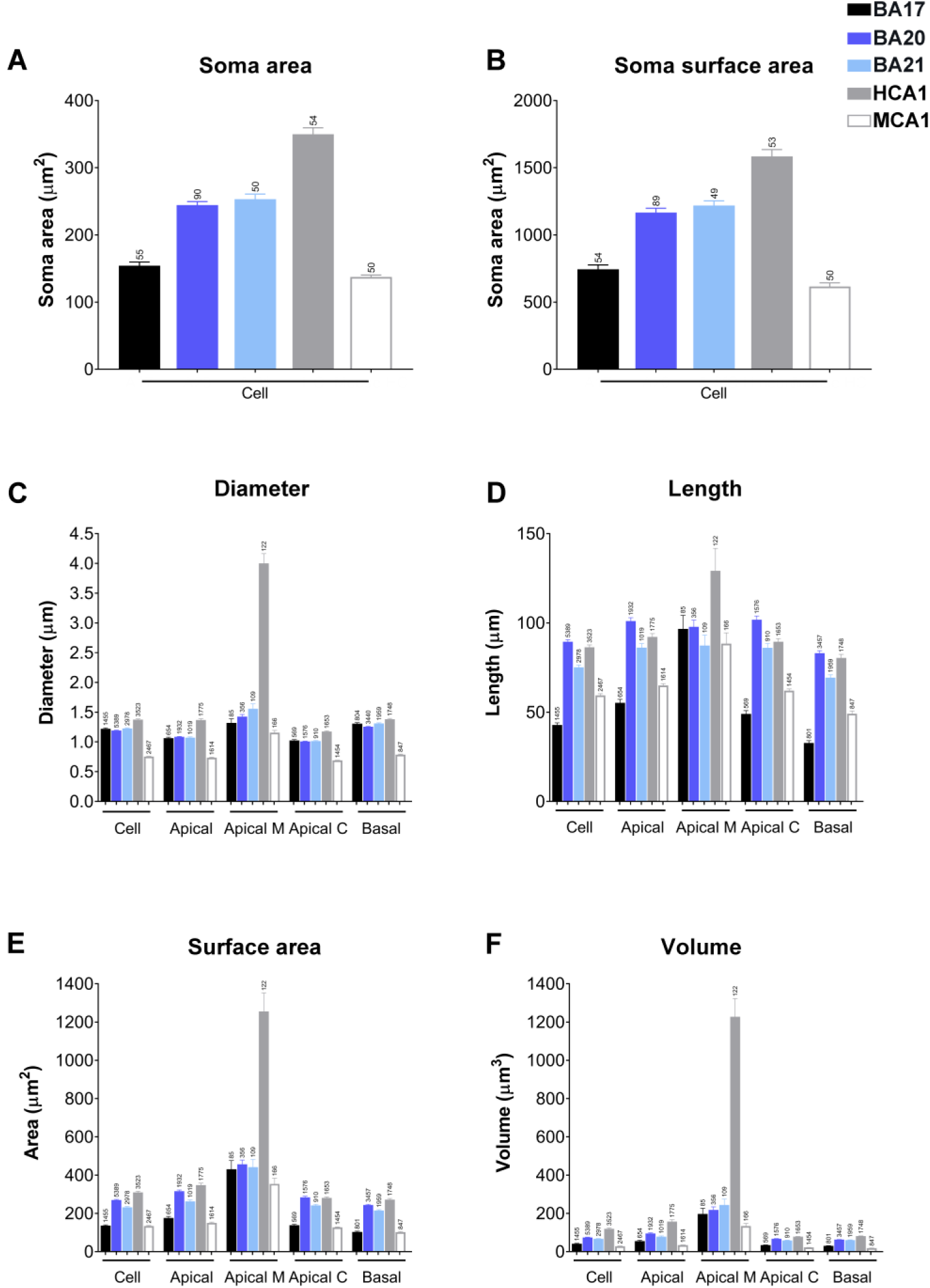
Graphs showing soma area (**A**), soma surface area (**B**), dendritic segment average diameter (**C**), segment length (**D**), segment surface area (**E**), and segment volume (**F**), expressed per cell and per dendritic compartment: apical arbor (including main apical dendrite and apical collateral dendrites together); main apical dendrite alone (Apical M); apical collateral dendrites alone (Apical C); and basal dendritic arbor from human BA17 (black), BA20 (dark blue), BA21 (light blue) cortical areas and human CA1 (grey) and mouse CA1 (white) regions. Measurements are reported as mean ± SEM. Only dendritic segments that were complete, and thus excluding incomplete endings, were included in this analysis. The statistical significance of the differences is shown in Supplementary Tables 1, 6.

**Supplementary Figure 7.**
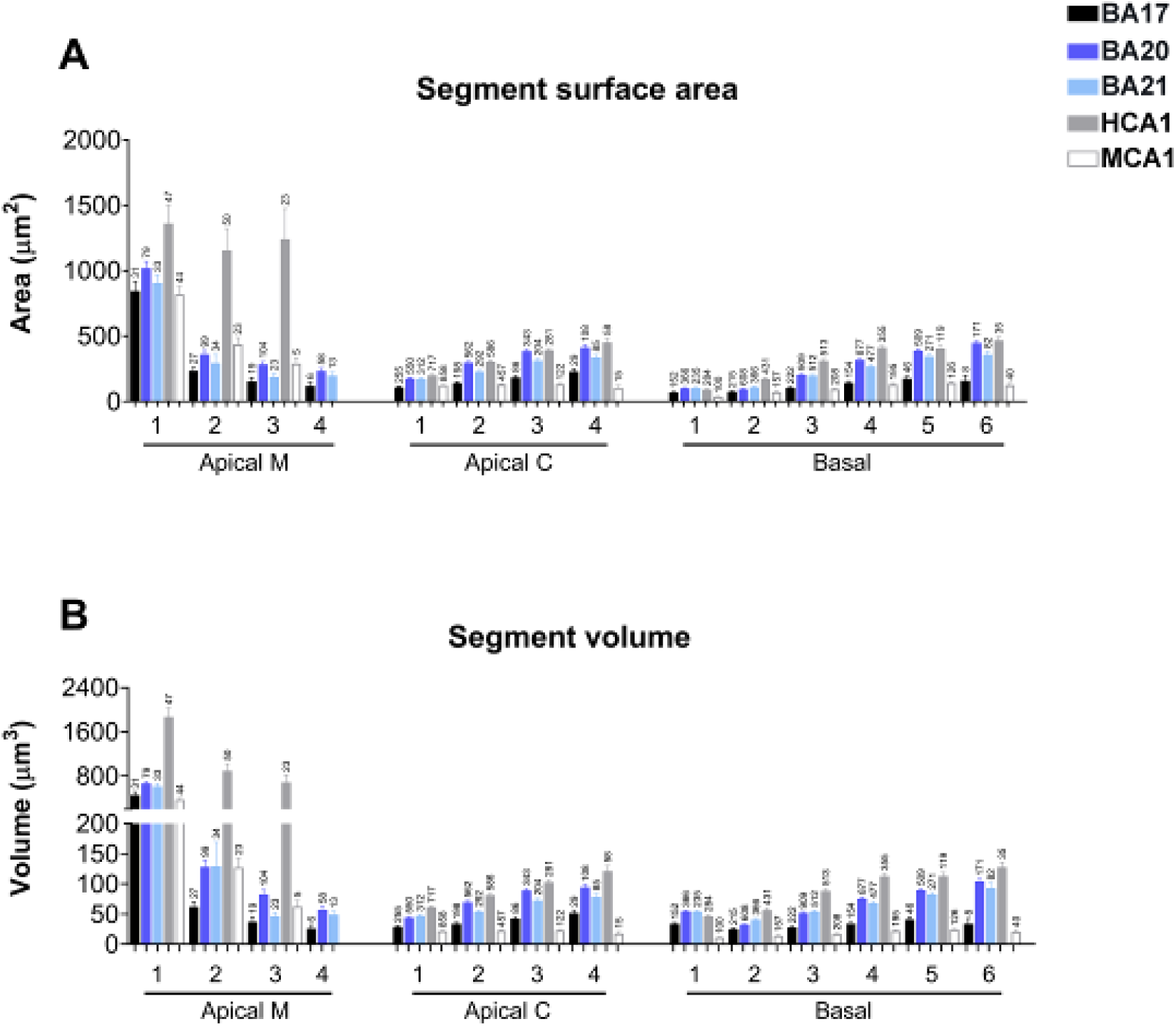
Graphs —per branch order— showing dendritic segment average surface area (**A**) and volume (**B**) per dendritic compartment: main apical dendrite (Apical M), apical collateral dendrites (Apical C), and basal arbor (Basal) from human BA17 (black), BA20 (dark blue), BA21 (light blue) cortical areas and human CA1 (grey) and mouse CA1 (white)regions. Measurements are reported as mean ± SEM. Only dendritic segments that were complete, and thus excluding incomplete endings, were included in this analysis. The statistical significance of the differences is shown in Supplementary Table 8.

**Supplementary Figure 8.**
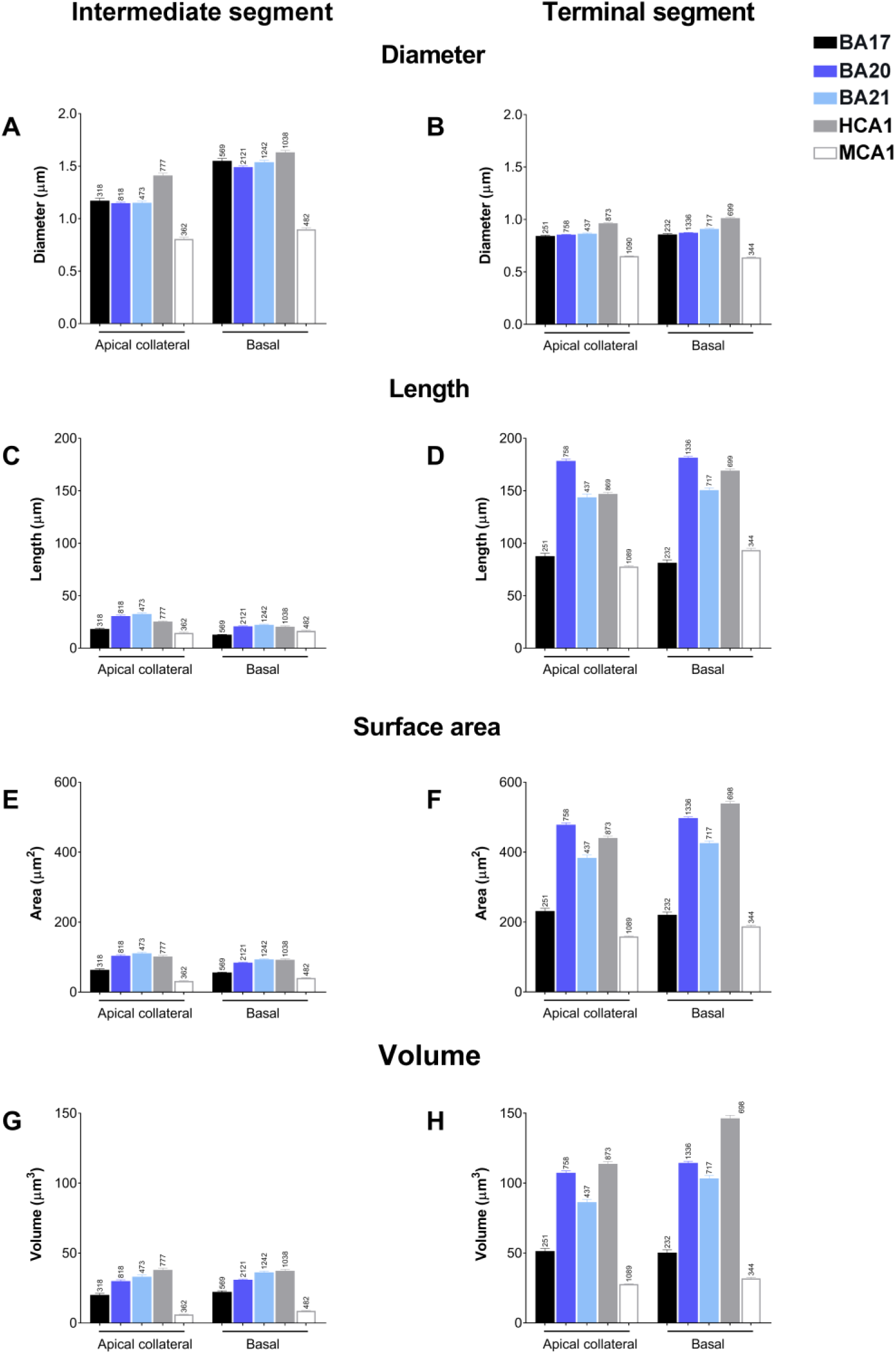
Graphs showing intermediate (**A, C, E, G**) versus terminal segment (**B, D, F, H**) diameters (A, B), lengths (C, D), surface area (E, F) and volume (G, H) for apical collateral (apical C) and basal dendrites from human BA17 (black), BA20 (dark blue), BA21 (light blue) cortical areas and human CA1 (grey) and mouse CA1 (white) regions. Measurements are reported as mean ± SEM. Only dendritic segments that were complete, and thus excluding incomplete endings, were included in this analysis.

**Supplementary Figure 9.**
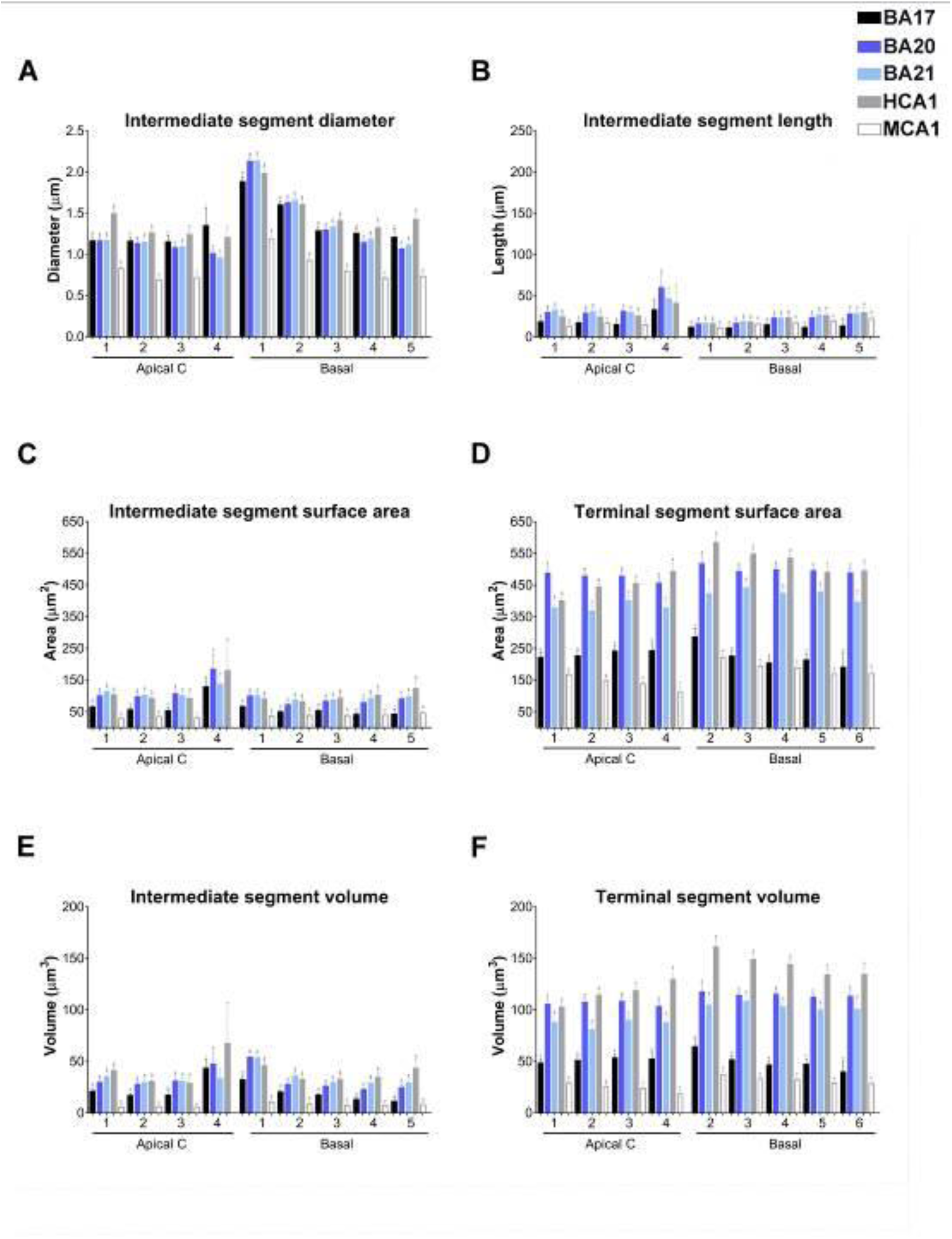
(**A, B**) Graphs showing —per branch order— intermediate segment diameter (A) and length (B) for apical collateral (apical C) and basal dendrites from human BA17 (black), BA20 (dark blue), BA21 (light blue) cortical areas and human CA1 (grey) and mouse CA1 (white) regions. (**C–F**) Graphs showing —per branch order— intermediate (C, E) versus terminal (D, F) segment surface area (C, D) and volume (E, F) for the same compartments and regions as above. Measurements are reported as mean ± SEM. Only dendritic segments that were complete, and thus excluding incomplete endings, were included in this analysis.

**Supplementary Figure 10.**
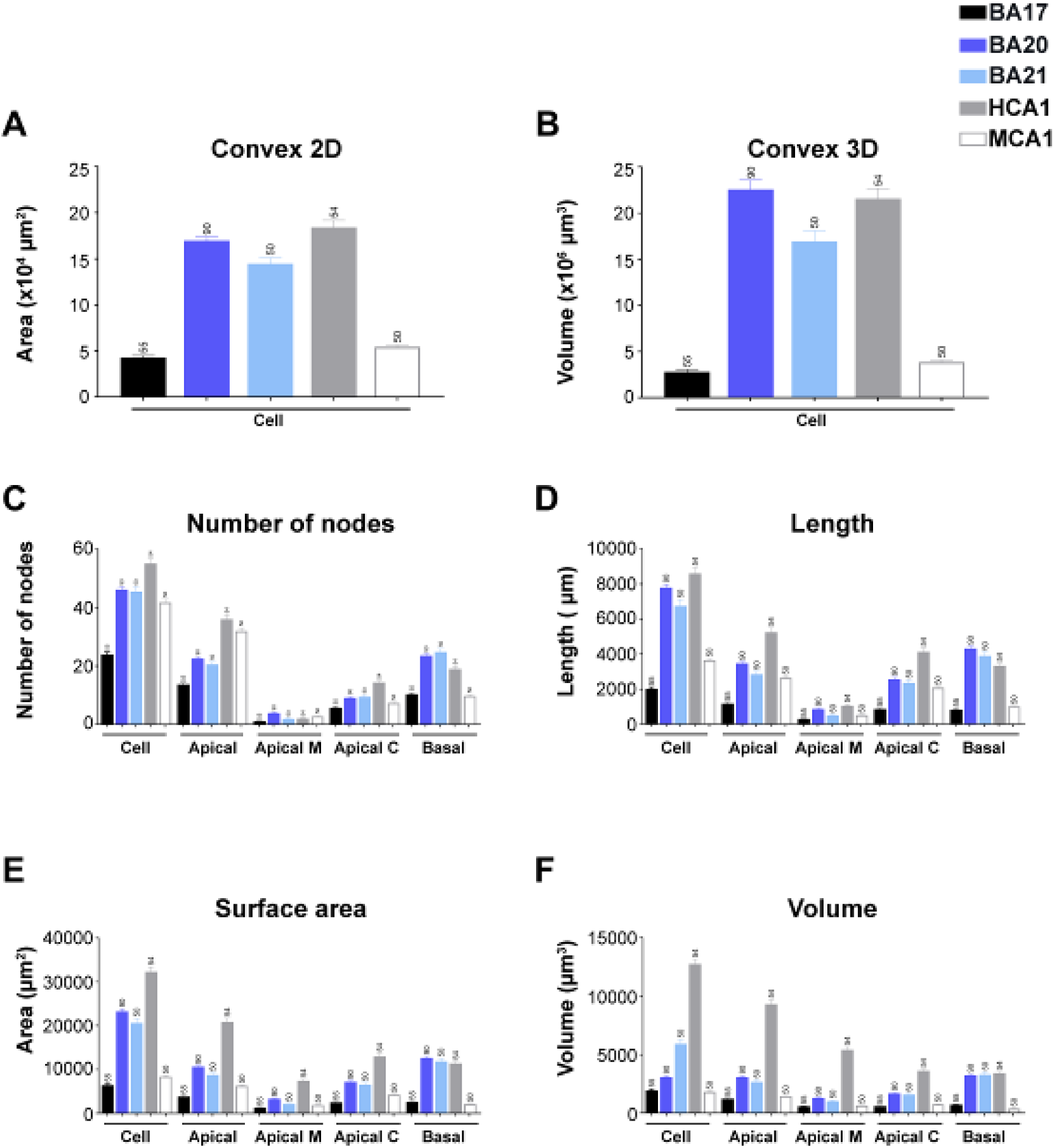
Graphs showing non-full measurements: convex 2D (**A**), convex 3D (**B**), number of nodes (**C**), dendritic length (**D**), dendritic surface area (**E**) and dendritic volume (**F**) from human BA17 (black), BA20 (dark blue), BA21 (light blue) cortical areas and human CA1 (grey) and mouse CA1 (white) regions.

**Supplementary Figure 11.**
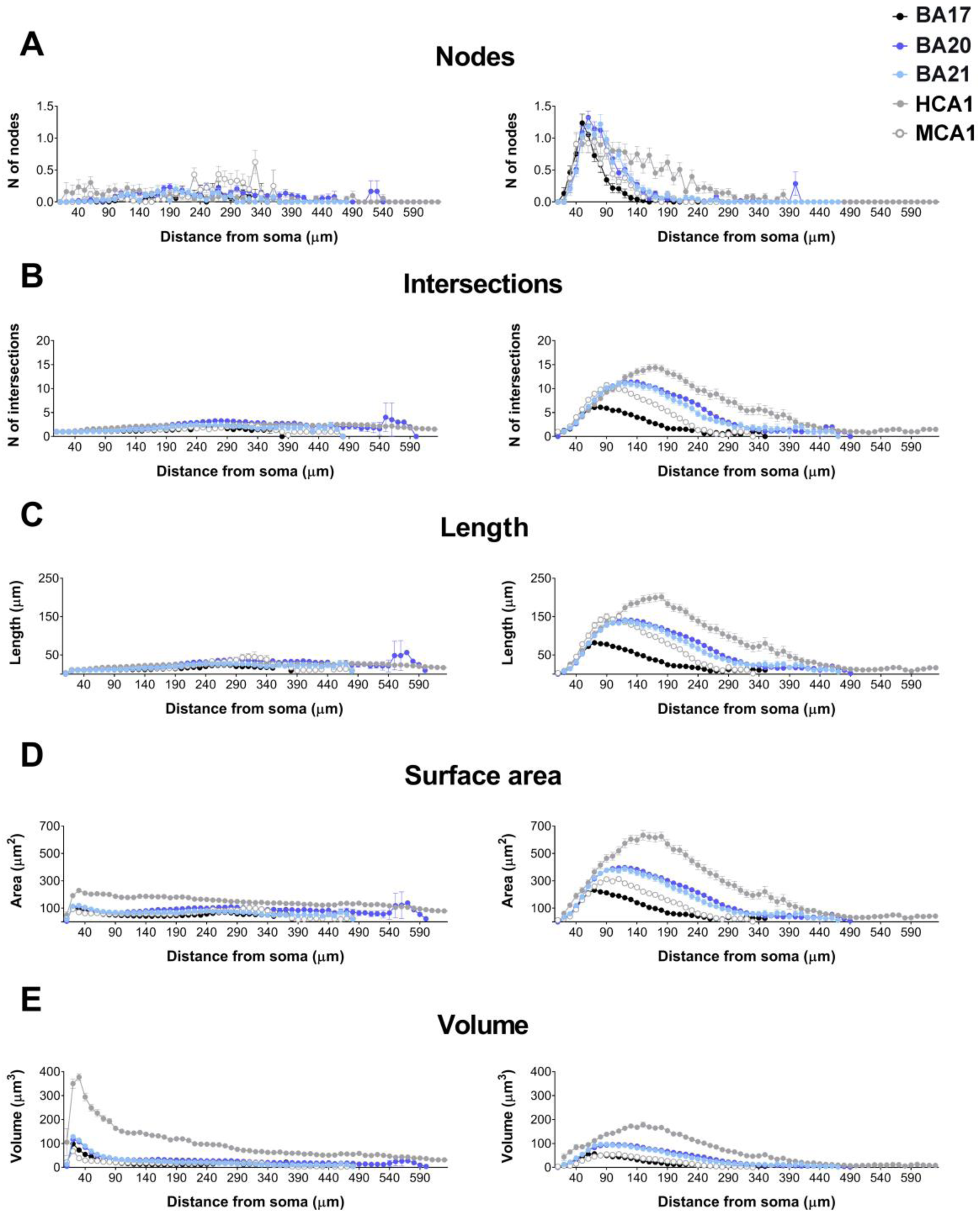
Graphs showing the number of nodes (**A**), dendritic intersections (**B**), dendritic length (**C**), dendritic surface area (**D**), and dendritic volume (**E**) distribution as a function of the distance from the soma in Apical M and Apical C compartments from human BA17 (black), BA20 (dark blue), BA21 (light blue), CA1human (grey) and CA1mouse (white) pyramidal neurons. Measurements are reported as mean ± SEM.

**Supplementary Figure 12.**
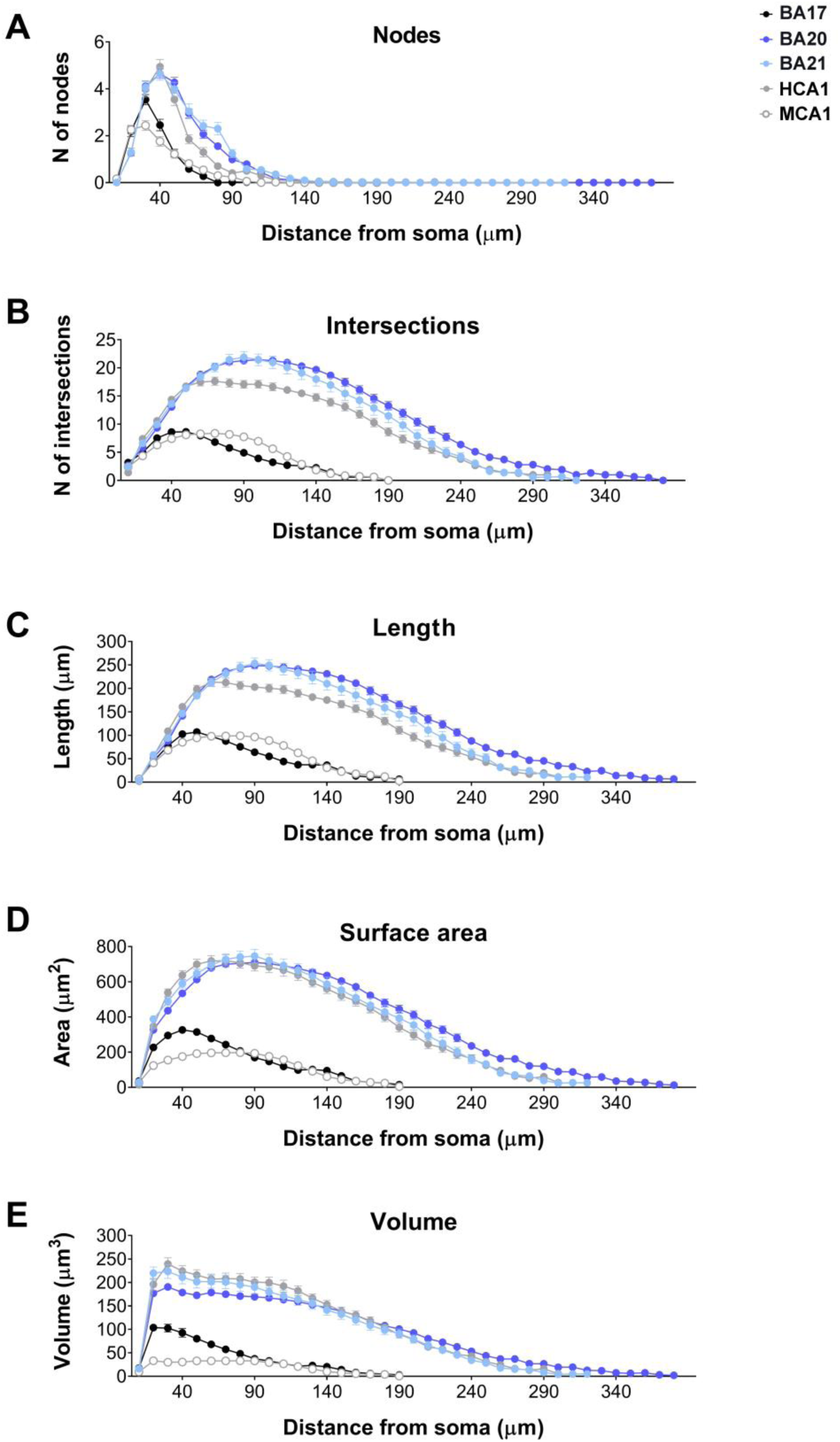
Graphs showing the number of nodes (**A**), dendritic intersections (**B**), dendritic length (**C**), dendritic surface area (**D**), and dendritic volume (**E**) distribution as a function of the distance from soma in basal compartments from human BA17 (black), BA20 (dark blue), BA21 (light blue), CA1human (grey) and CA1mouse (white) pyramidal neurons. Measurements are reported as mean ± SEM.’

**Supplementary Figure 13.**
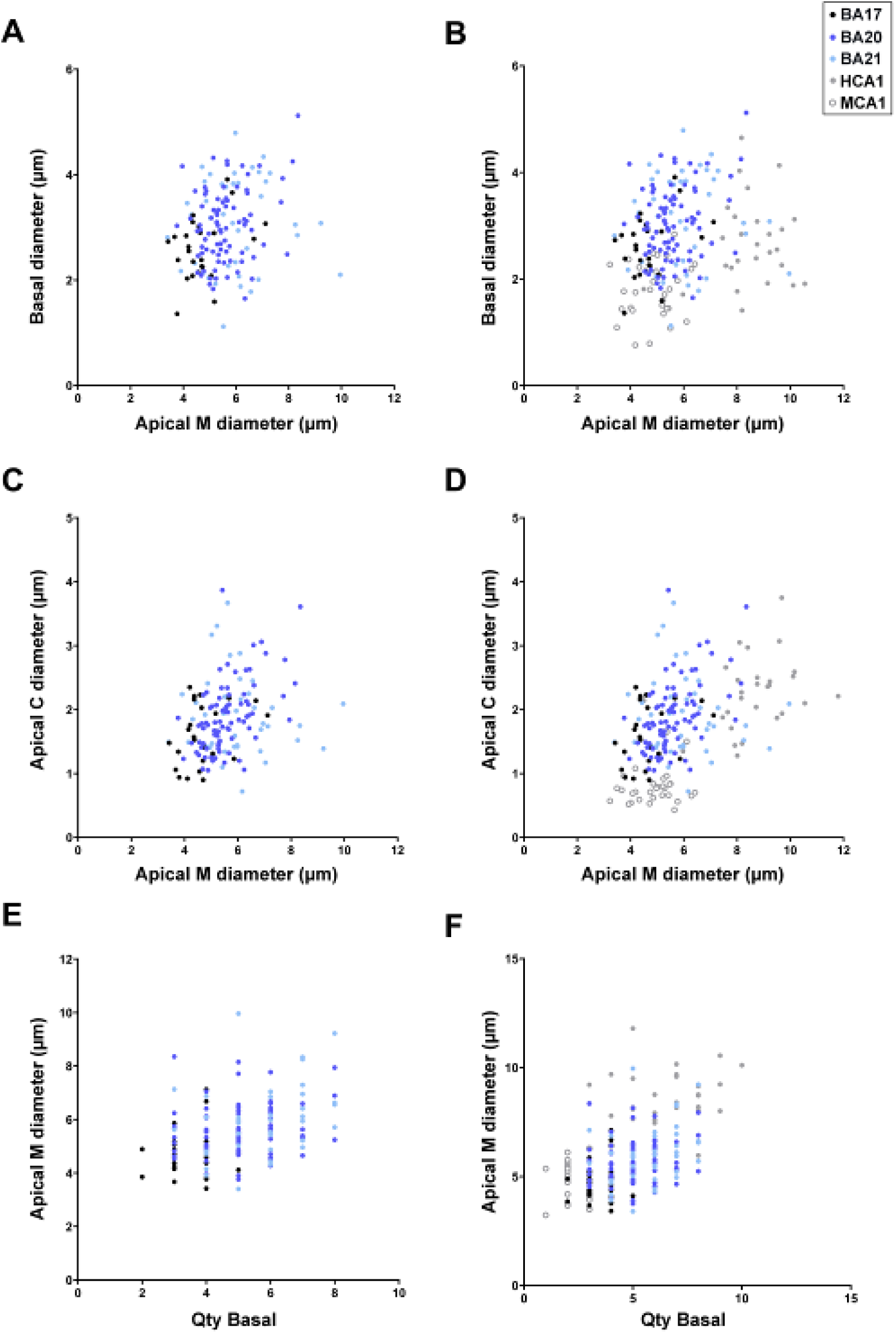
Correlation analyses between various morphological parameters analyzed. Each point represents the values obtained in one cell from human BA17 (black), BA20 (dark blue), BA21 (light blue) cortical areas (left column). Additional correlations, including human CA1 (grey) and mouse CA1 (open circles) regions, are shown (right column). Significant correlations were classified as weak [Spearman’s rho (r) value lower than 0.40], moderate (0.4<r<0.7), and strong (r>0.7). See also Supplementary Table 11.

**Supplementary Figure 14.**
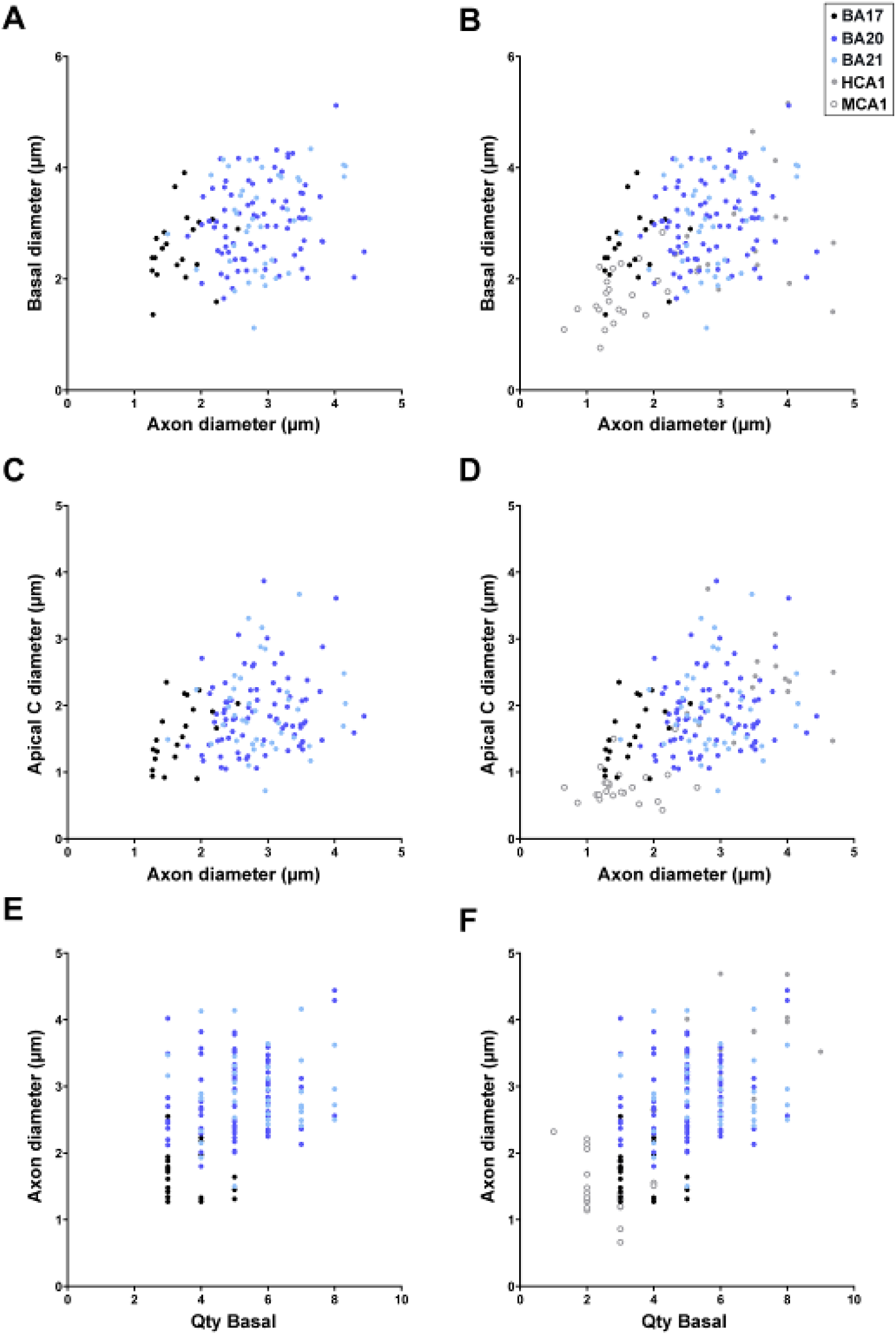
Correlation analyses between various morphological parameters analyzed. Each point represents the values obtained in one cell from human BA17 (black), BA20 (dark blue), BA21 (light blue) cortical areas (left column). Additional correlations, including human CA1 (grey) and mouse CA1 (open circles) regions, are shown (right column). Significant correlations were classified as weak [Spearman’s rho (r) value lower than 0.40], moderate (0.4<r<0.7), and strong (r>0.7). See also Supplementary Table 11.

**Supplementary Figure 15.**
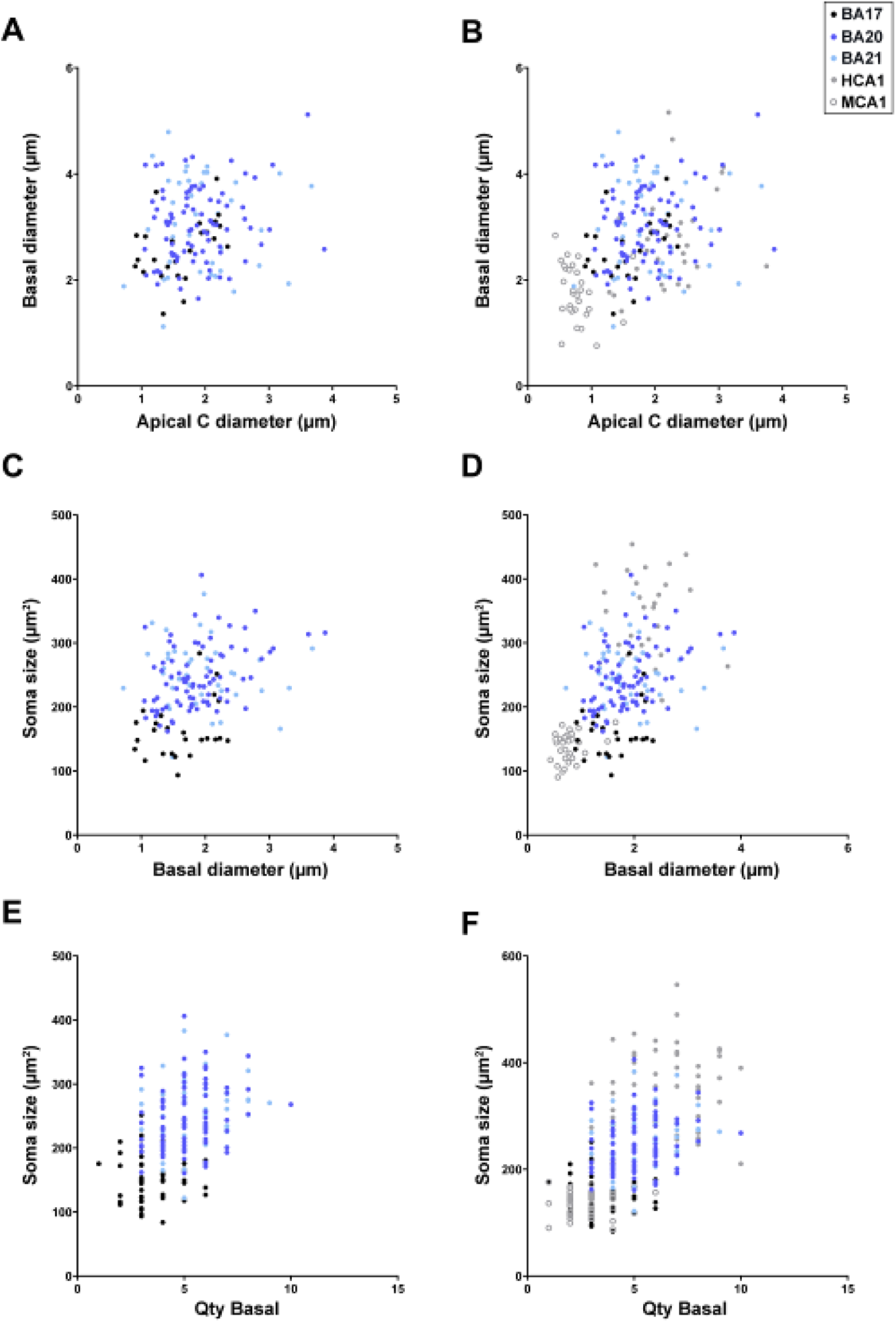
Correlation analyses between various morphological parameters analyzed. Each point represents the values obtained in one cell from human BA17 (black), BA20 (dark blue), BA21 (light blue) cortical areas (left column). Additional correlations, including human CA1 (grey) and mouse CA1 (open circles) regions, are shown (right column). Significant correlations were classified as weak [Spearman’s rho (r) value lower than 0.40], moderate (0.4<r<0.7), and strong (r>0.7). See also Supplementary Table 11.

**Supplementary Figure 16.**
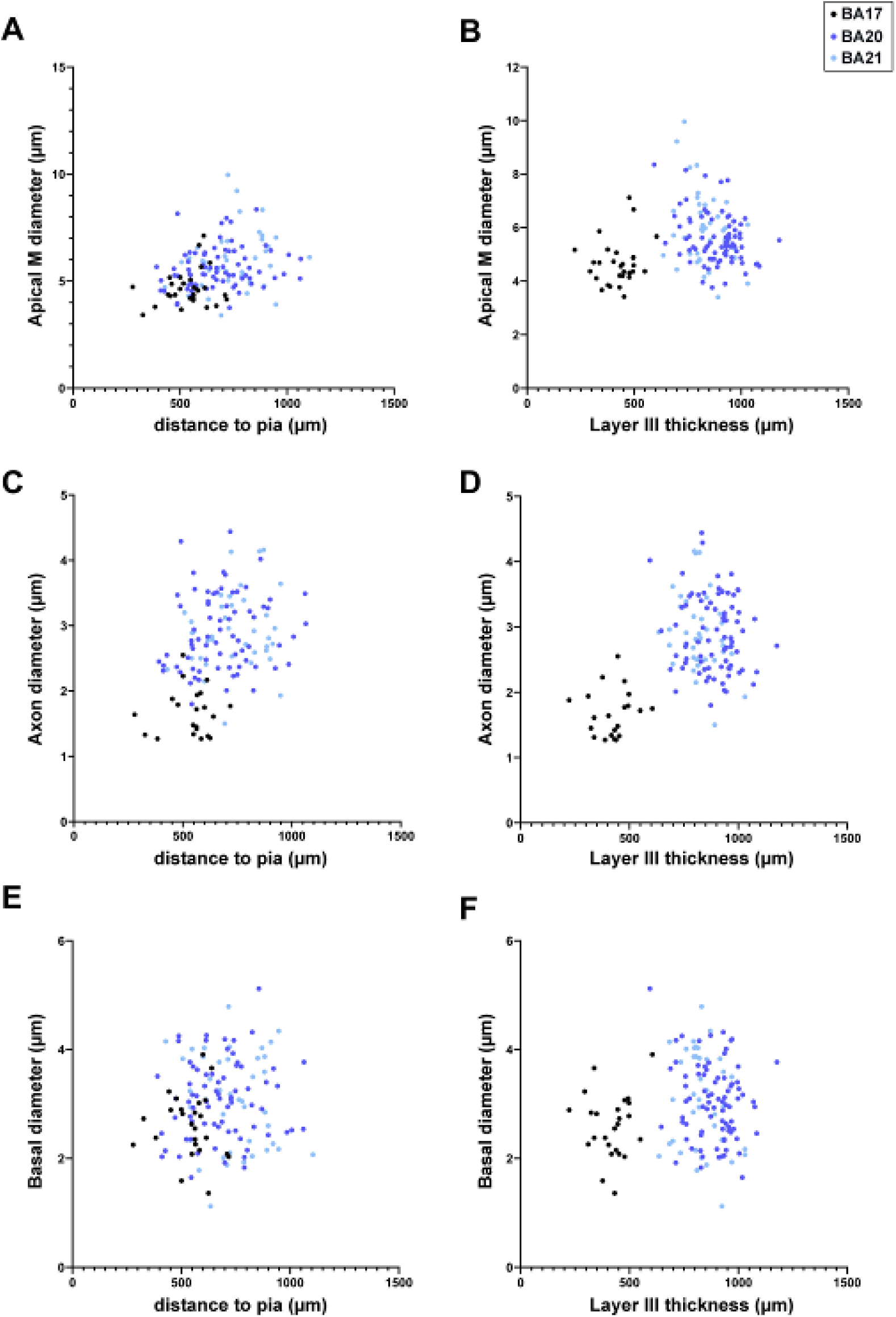
Correlation analyses between various morphological parameters analyzed. Each point represents the values obtained in one cell from human BA17 (black), BA20 (dark blue), BA21 (light blue) cortical areas and human CA1 (grey) and mouse CA1 (open circles) regions. Significant correlations were classified as weak [Spearman’s rho (r) value lower than 0.40], moderate (0.4<r<0.7), and strong (r>0.7). See also Supplementary Table 11.

**Supplementary Figure 17.**
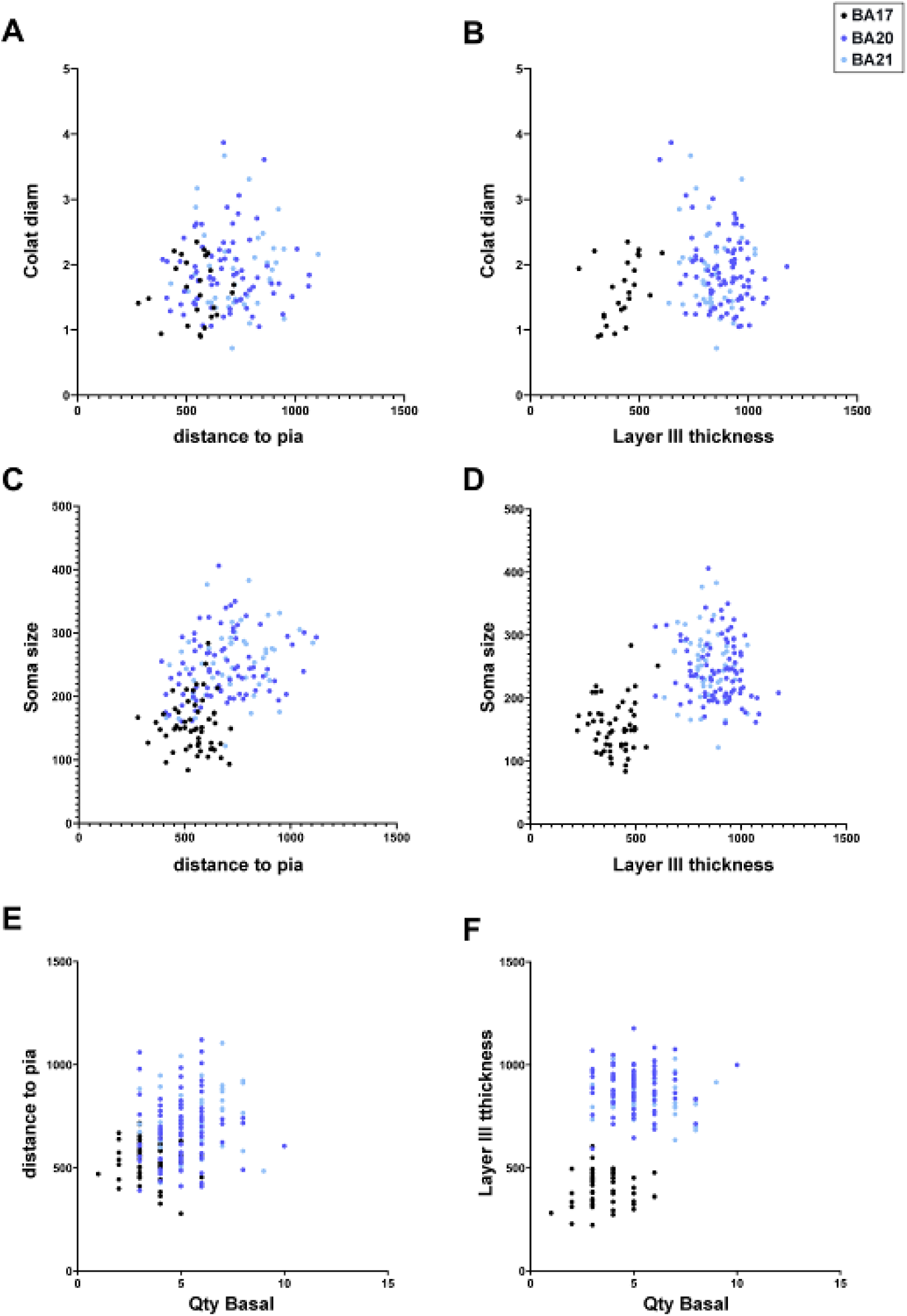
Correlation analyses between various morphological parameters analyzed. Each point represents the values obtained in one cell from human BA17 (black), BA20 (dark blue), BA21 (light blue) cortical areas and human CA1 (grey) and mouse CA1 (open circles) regions. Significant correlations were classified as weak [Spearman’s rho (r) value lower than 0.40], moderate (0.4<r<0.7), and strong (r>0.7). See also Supplementary Table 11.

**Supplementary Table 1:** Statistical comparisons (Kruskal-Wallis test) of mean values of morphological variables per region shown in Figure 3 A, B (**a**) and Supplementary figure 1 (**a**, **b**).

|  | BA17<br>vs<br>BA20 | BA17<br>vs<br>BA21 | BA20<br>vs<br>BA21 |
| --- | --- | --- | --- |
| Soma volume (Fig. 3A) | **** | **** | ns |
| Number of primary basal dendrites (Fig. 3B) | **** | **** | ns |
| Soma area(Supl. fig 1A) | **** | **** | ns |
| Soma surface area (Supl. fig 1B) | **** | **** | ns |

**b.**
|  | Segment diameter |  |  | Segment length |  |  | Segment surface area |  |  | Segment volume |  |  |
| --- | --- | --- | --- | --- | --- | --- | --- | --- | --- | --- | --- | --- |
|  | BA17<br>vs<br>BA20 | BA17<br>vs<br>BA21 | BA20<br>vs<br>BA21 | BA17<br>vs<br>BA20 | BA17<br>vs<br>BA21 | BA20<br>vs<br>BA21 | BA17<br>vs<br>BA20 | BA17<br>vs<br>BA21 | BA20<br>vs<br>BA21 | BA17<br>vs<br>BA20 | BA17<br>vs<br>BA21 | BA20<br>vs<br>BA21 |
| Cell | ns | ns | ns | **** | **** | **** | **** | **** | *** | **** | **** | *** |
| Apical (M+C) | *** | ns | ** | **** | **** | ** | **** | **** | *** | **** | **** | **** |
| Apical M | ns | ns | ns | ns | ns | ns | ns | ns | ns | ns | ns | ns |
| Apical C | * | ns | ns | **** | **** | * | **** | **** | ** | **** | **** | ** |
| Basal | ns | ns | ** | **** | **** | * | **** | **** | ns | **** | **** | ns |

**Supplementary Table 2:** Statistical comparisons of dendritic diameter values per distance from soma from Fig 3 C-F.

| | $\mu\text{m}$ | Apical M | | | Apical C | | | Basal | | | Axon | | |
| --- | --- | --- | --- | --- | --- | --- | --- | --- | --- | --- | --- | --- | --- |
|  |  | BA17<br>vs.<br>BA20 | BA17<br>vs.<br>BA21 | BA20<br>vs.<br>BA21 | BA17<br>vs.<br>BA20 | BA17<br>vs.<br>BA21 | BA20<br>vs.<br>BA21 | BA17<br>vs.<br>BA20 | BA17<br>vs.<br>BA21 | BA20<br>vs.<br>BA21 | BA17<br>vs.<br>BA20 | BA17<br>vs.<br>BA21 | BA20<br>vs.<br>BA21 |
| Dendritic<br>Diameter | 20 | **** | **** | ns | ns | ns | ns | *** | **** | ns | **** | **** | ns |
|  | 30 | **** | **** | ns | ns | ns | ns | *** | **** | ns | **** | **** | ns |
|  | 40 | **** | **** | ns | ns | ns | ns | ** | **** | ns | * | ** | ns |
|  | 50 | *** | ** | ns | * | ns | ns | *** | **** | ns | ns | ns | ns |
|  | 60 | *** | *** | ns | * | * | ns | *** | *** | ns | ns | ns | ns |
|  | 70 | **** | **** | ns | *** | ** | ns | ** | *** | ns | ns | ns | ns |
|  | 80 | **** | **** | ns | ** | ** | ns | ** | ** | ns | ns | ns | ns |
|  | 90 | **** | *** | ns | *** | * | ns | *** | *** | ns | - | - | - |
|  | 100 | **** | ** | ns | ** | ns | ns | **** | ** | ns | - | - | - |
|  | 110 | **** | *** | ns | * | ns | ns | **** | ** | ns | - | - | - |
|  | 120 | **** | *** | ns | ns | ns | ns | *** | ** | ns | - | - | - |
|  | 130 | **** | ** | ns | ns | ns | ns | * | ns | ns | - | - | - |
|  | 140 | **** | ** | ns | ns | ns | ns | ** | * | ns | - | - | - |
|  | 150 | **** | ** | ns | ns | ns | ns | ** | ns | ns | - | - | - |
|  | 160 | **** | ** | ns | ns | ns | ns | ns | ns | ns | - | - | - |
|  | 170 | **** | ** | ns | ns | ns | ns | - | - | - | - | - | - |
|  | 180 | **** | ** | ns | ns | ns | ns | - | - | - | - | - | - |
|  | 190 | *** | ns | ns | ns | ns | ns | - | - | - | - | - | - |
|  | 200 | ** | ns | ns | ns | ns | ns | - | - | - | - | - | - |
|  | 210 | *** | ns | * | ns | ns | ns | - | - | - | - | - | - |
|  | 220 | *** | ns | ns | ns | ns | ns | - | - | - | - | - | - |
|  | 230 | * | ns | ns | ns | ns | ns | - | - | - | - | - | - |
|  | 240 | * | ns | ns | ns | ns | ns | - | - | - | - | - | - |
|  | 250 | ns | ns | ns | ns | ns | ns | - | - | - | - | - | - |
|  | 260 | ns | ns | ns | ns | ns | ns | - | - | - | - | - | - |
|  | 270 | ns | ns | ns | ns | ns | ns | - | - | - | - | - | - |

**Supplementary Table 3:**
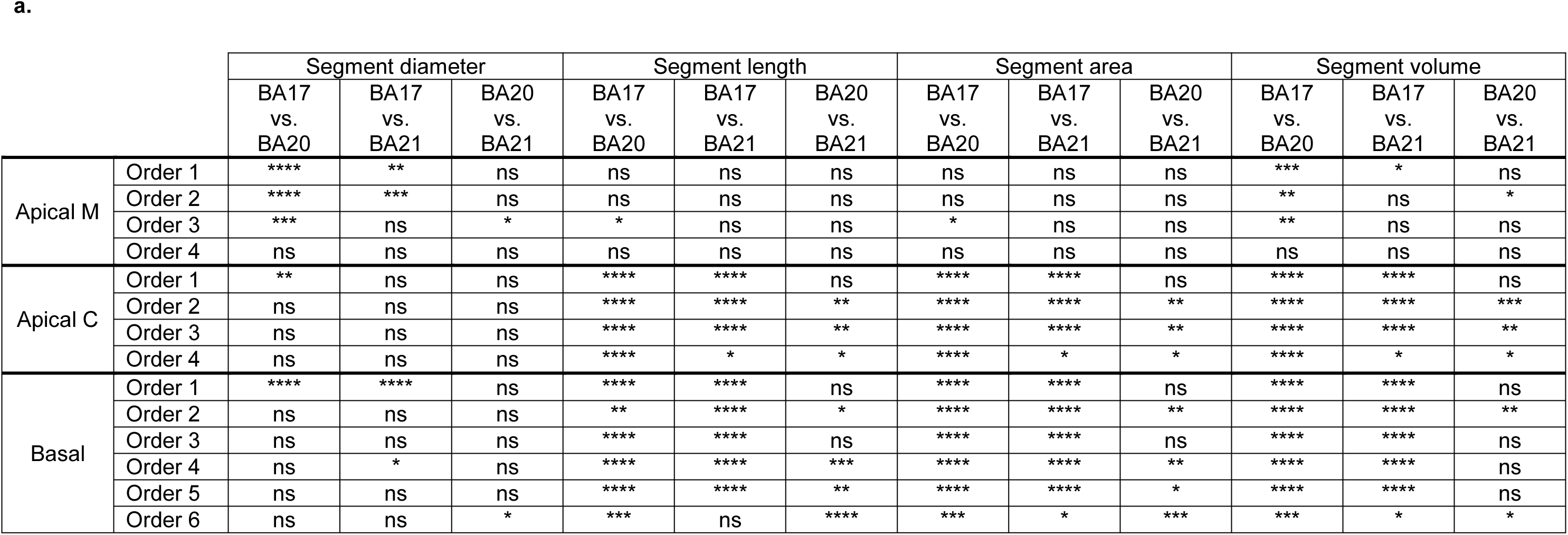
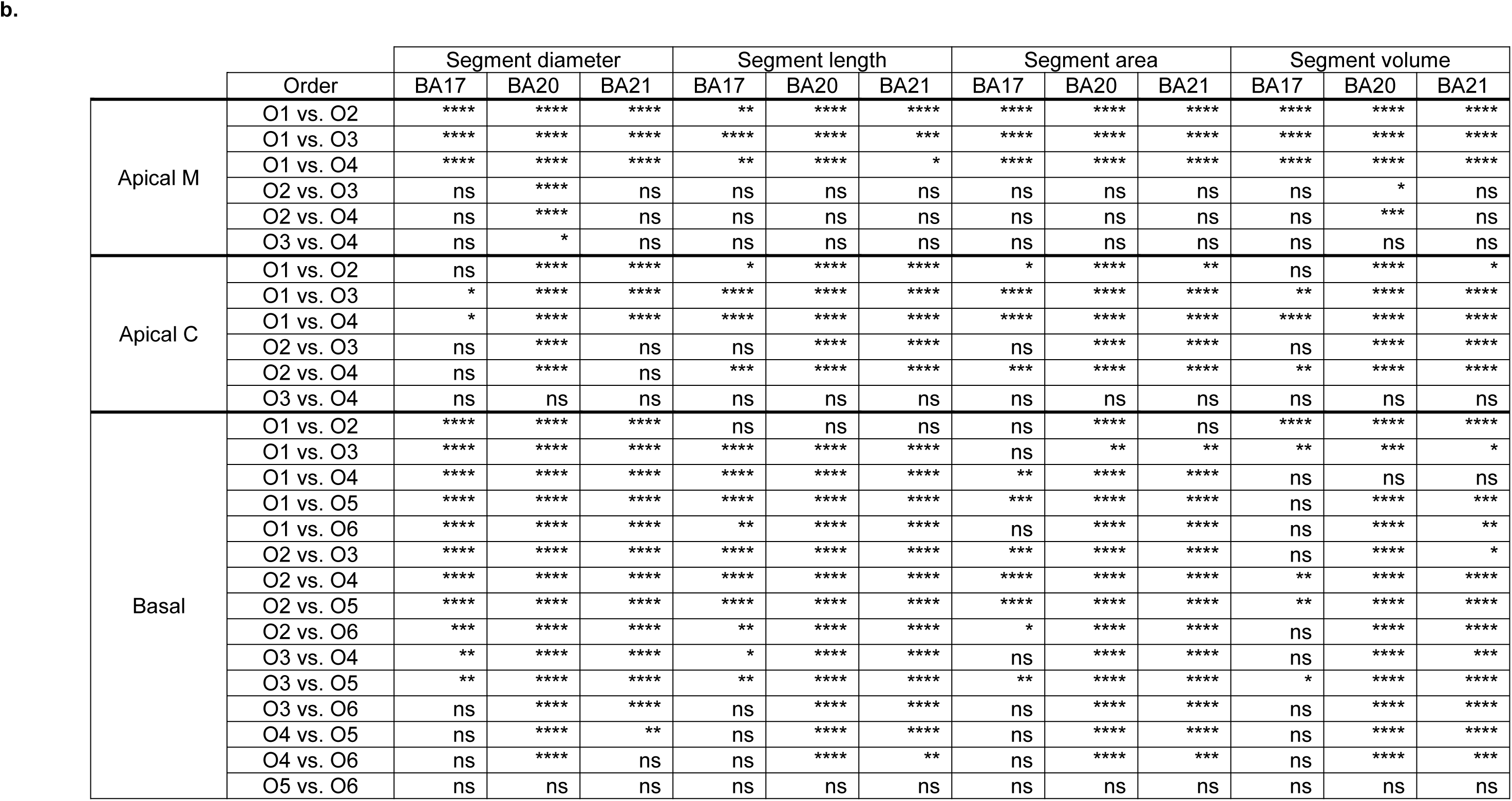
Statistical comparisons of segment diameter, length, surface area and volume values per branch order, between compartments (**a**) and orders (**b**) from graphs shown in Figure 4 and Supplementary Figure 5A.

**Supplementary Table 4:**
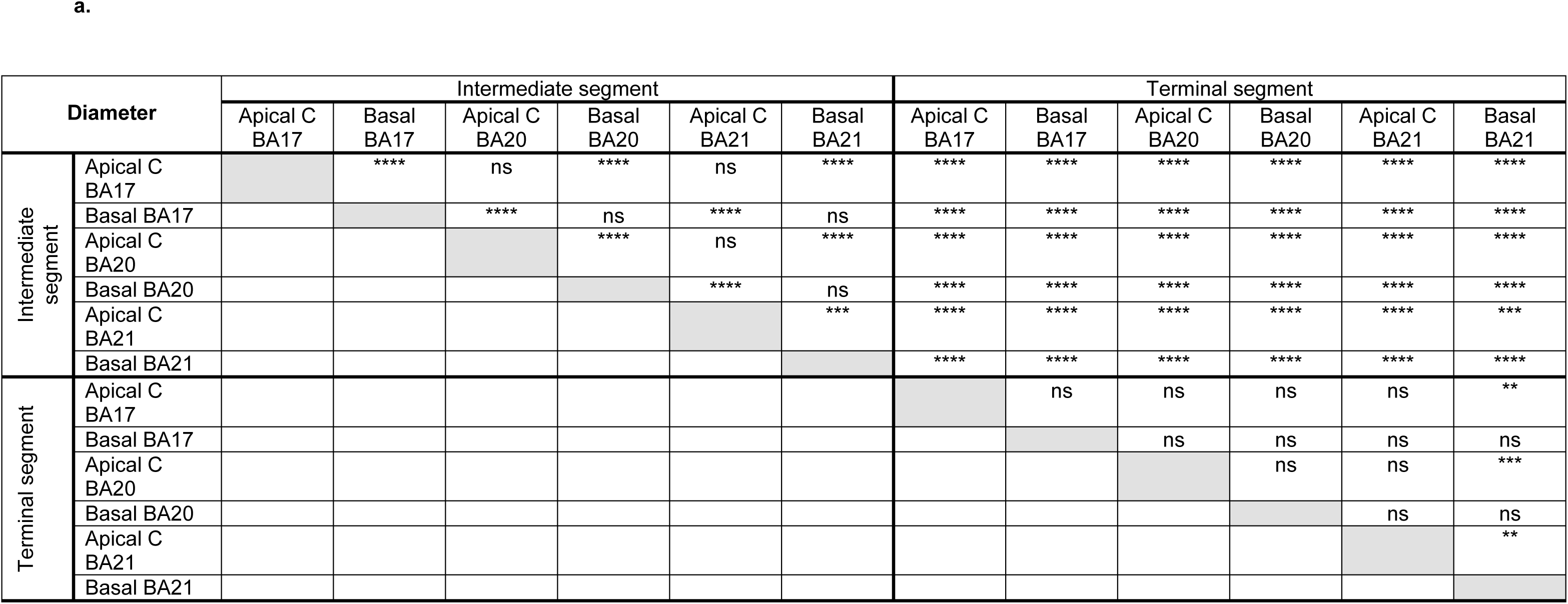
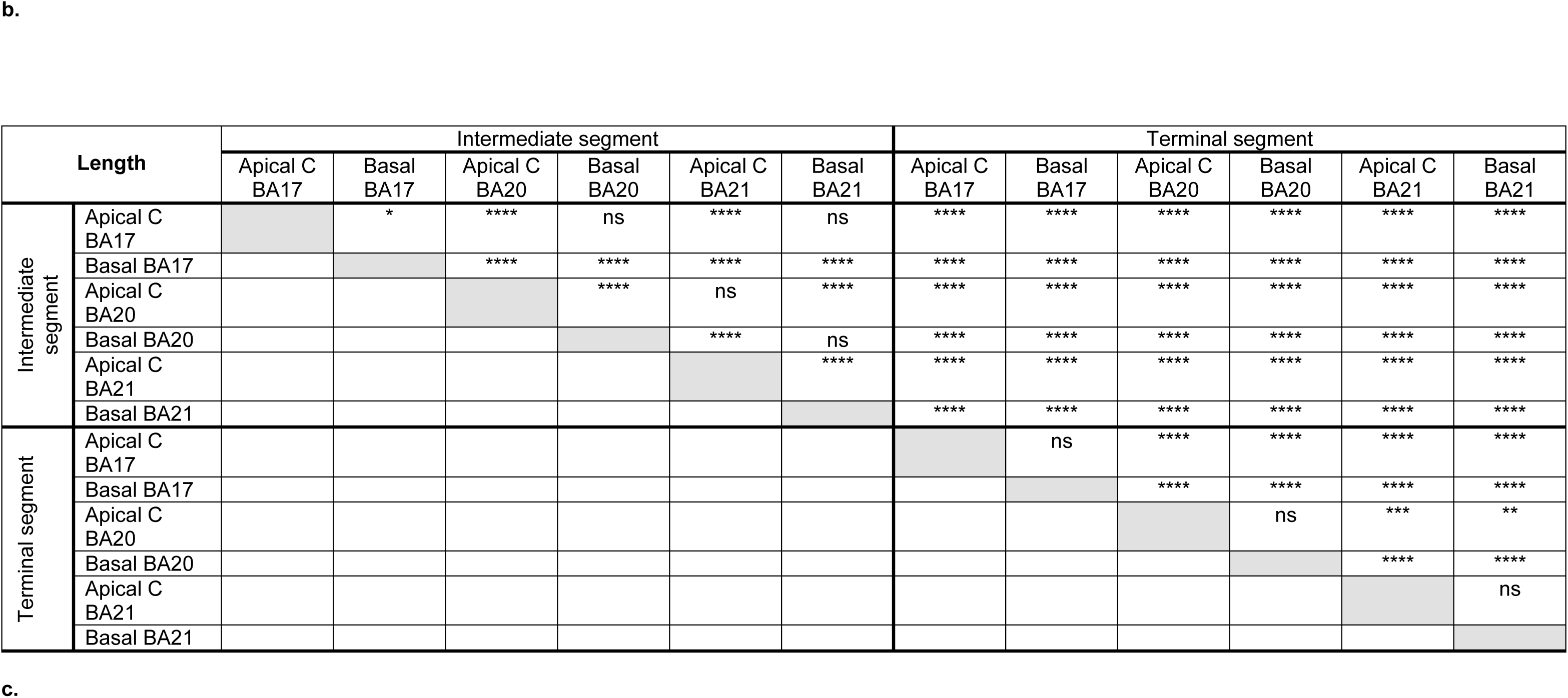
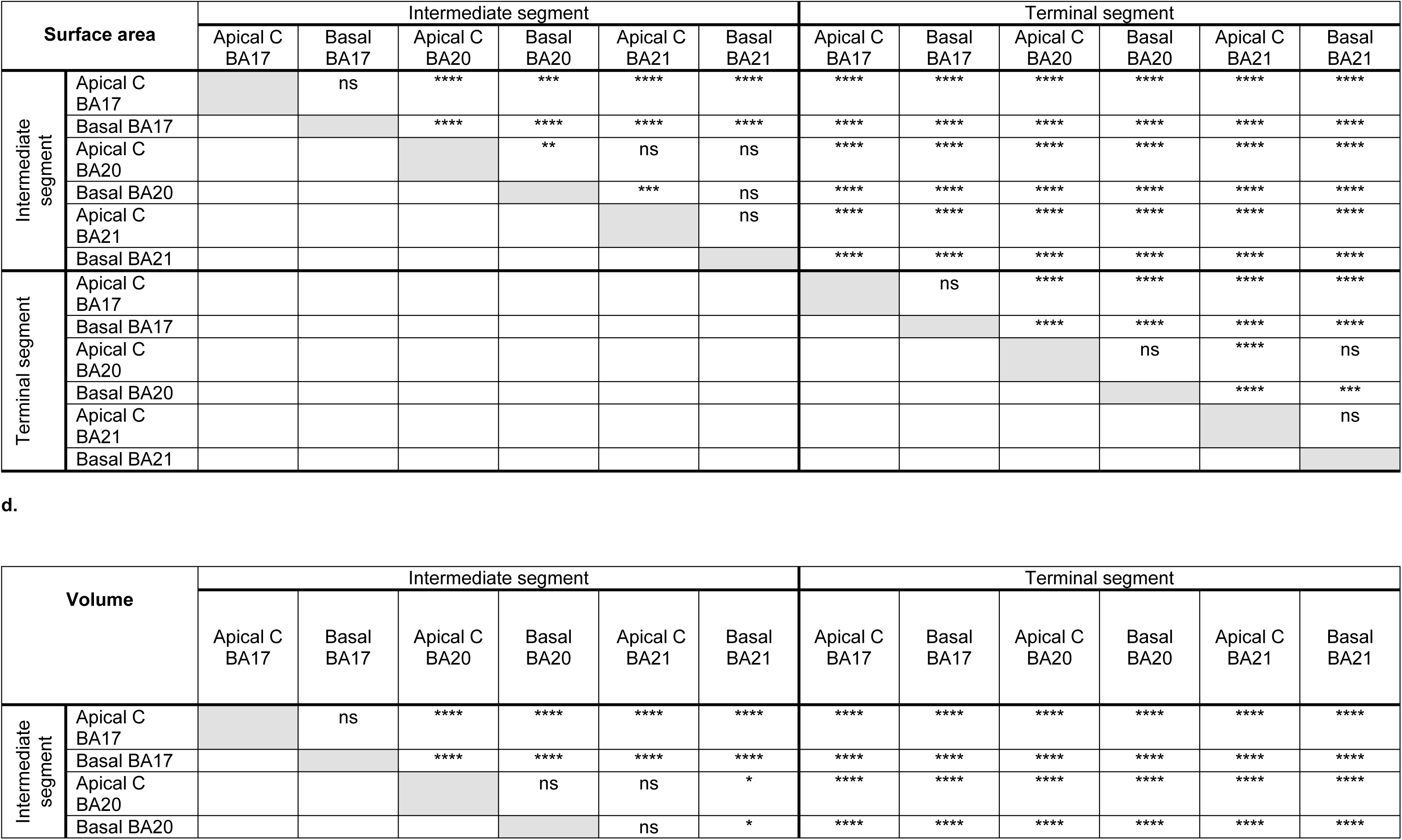
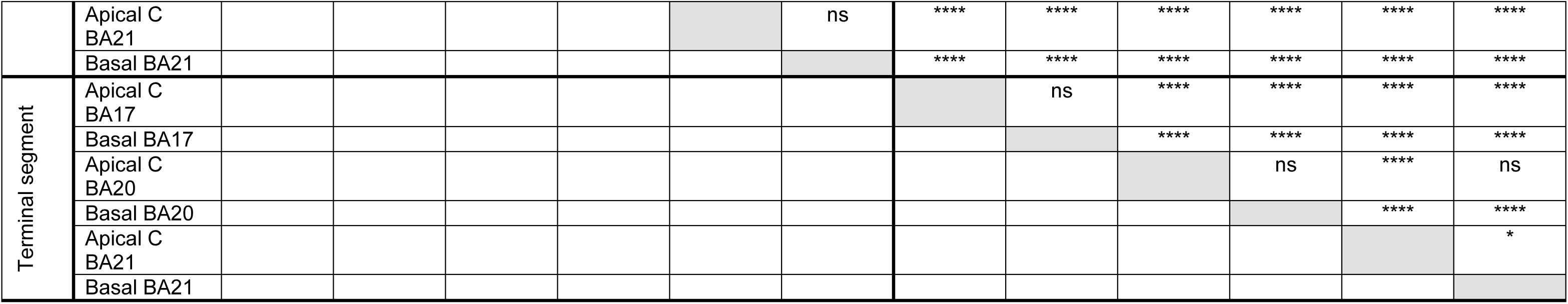
Statistical comparisons of segment diameter (**a**), length (**b**), surface area (**c**) and volume (**d**) values sorted per intermediate and terminal segments, from graphs shown in Supplementary Figures 4 and 5 B, C.

**Supplementary Table 5:**
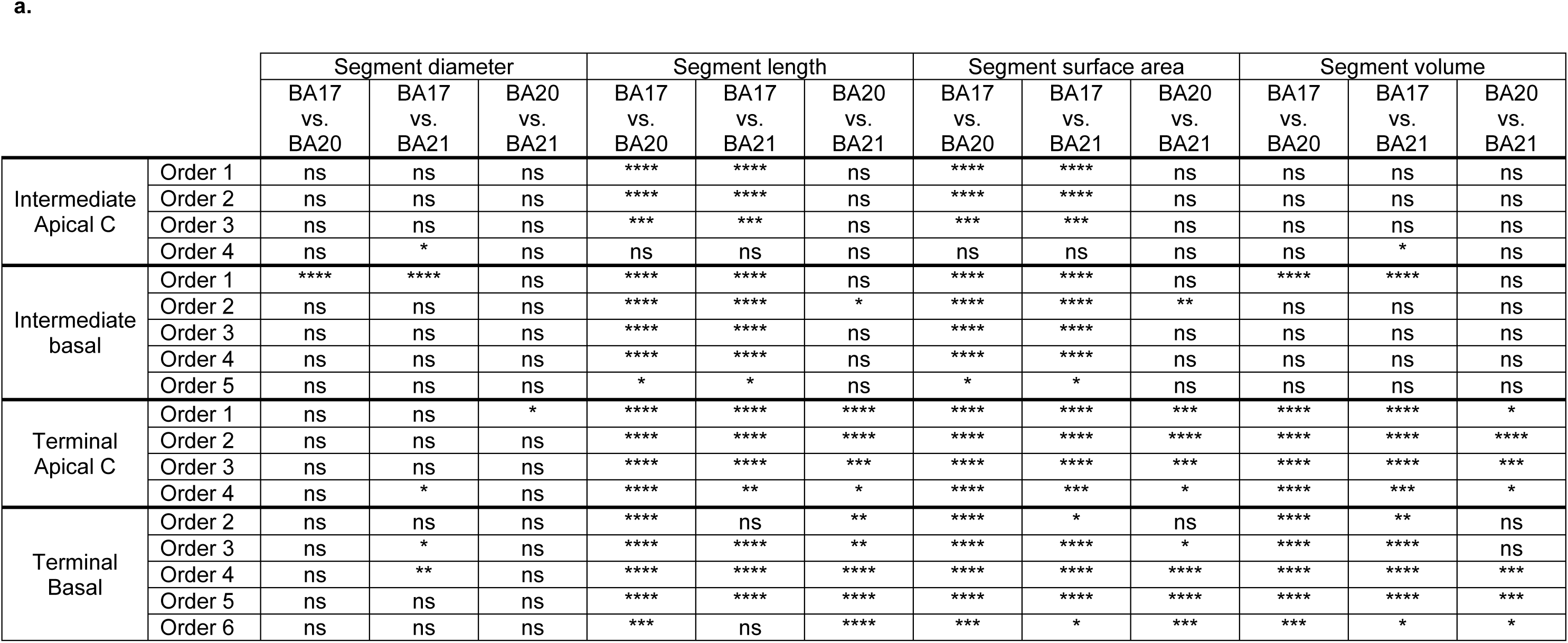
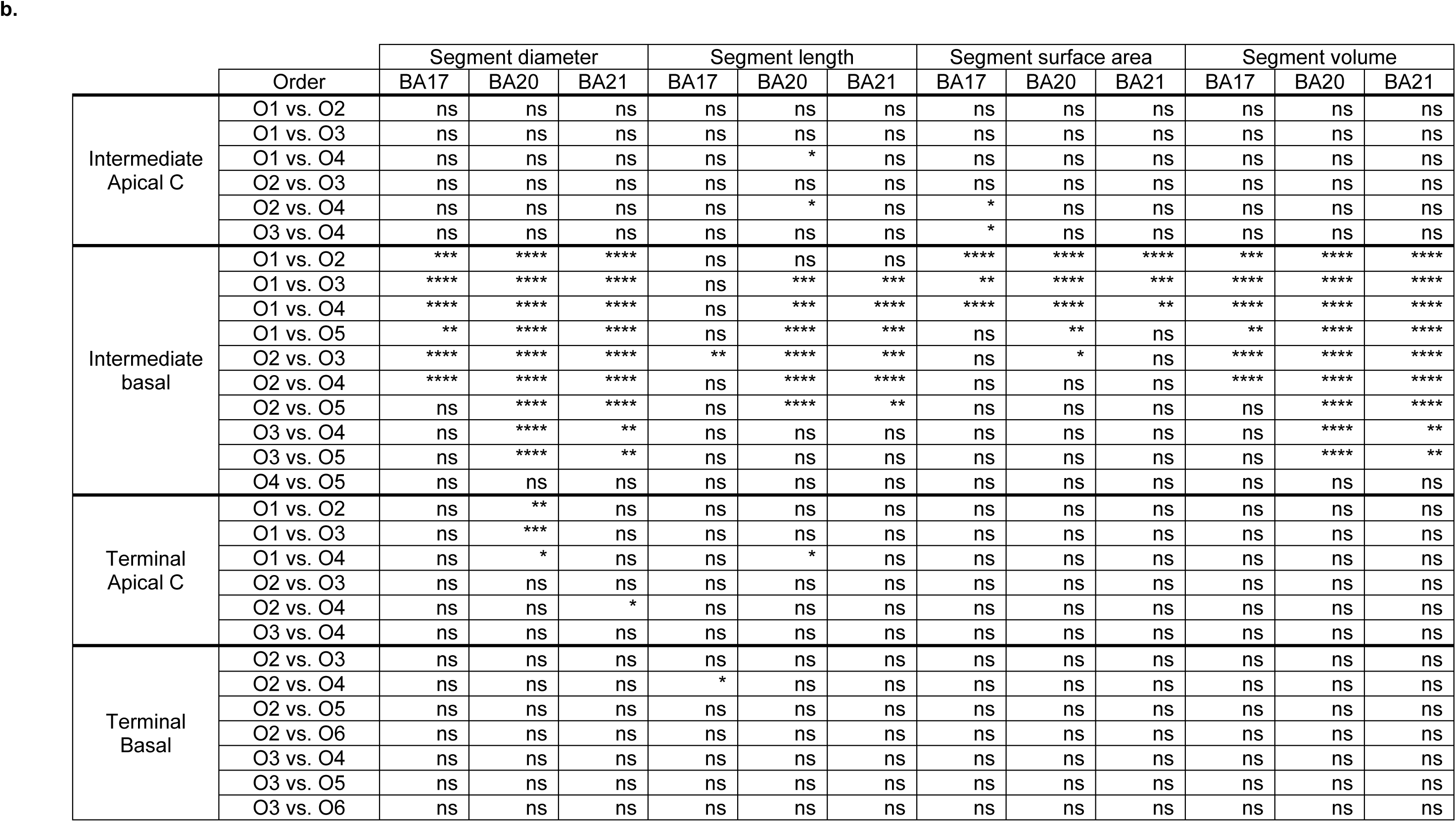
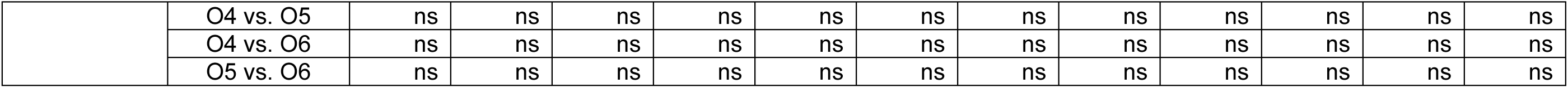
Statistical comparisons of segment diameters/length/surface area and volume values per branch order and dendritic segment type (intermediate/terminal), between compartments (**a**) and orders (**b**) from graphs shown in Figure 5, Supplementary Figure 5 D, E.

**Supplementary Table 6:** Statistical comparisons (Kruskal-Wallis test) of mean values of morphological variables per region shown in Figure 7 A, B (**a**) and Supplementary figures 6 and 10 A, B (**a, b**).

|  | BA17<br>vs<br>BA20 | BA17<br>vs<br>BA21 | BA17<br>vs<br>HCA1 | BA17<br>vs<br>MCA1 | BA20<br>vs<br>BA21 | BA20<br>vs<br>HCA1 | BA20<br>vs<br>MCA1 | BA21<br>vs<br>HCA1 | BA21<br>vs<br>MCA1 | HCA1<br>vs<br>MCA1 |
| --- | --- | --- | --- | --- | --- | --- | --- | --- | --- | --- |
| Soma volume (Fig. 7A) | **** | **** | **** | ns | ns | *** | **** | * | **** | **** |
| Number of primary basal dendrites (Fig. 7B) | **** | **** | **** | ns | ns | * | **** | ns | **** | **** |
| Soma area (Supl. Fig. 6A) | **** | **** | **** | ns | ns | **** | **** | *** | **** | **** |
| Soma surface area (Supl. Fig. 6B) | **** | **** | **** | ns | ns | **** | **** | * | **** | **** |
| Cell area (Convex 2D) (Supl. Fig. 10A) | **** | **** | **** | ns | ns | ns | **** | ns | **** | **** |
| Cell volume (Convex 3D) (Supl. Fig. 10B) | **** | **** | **** | ns | ns | ns | **** | ns | **** | **** |

**b.**
| Segment<br>diameter | BA17<br>vs<br>BA20 | BA17<br>vs<br>BA21 | BA17<br>vs<br>HCA1 | BA17<br>vs<br>MCA1 | BA20<br>vs<br>BA21 | BA20<br>vs<br>HCA1 | BA20<br>vs<br>MCA1 | BA21<br>vs<br>HCA1 | BA21<br>vs<br>MCA1 | HCA1<br>vs<br>MCA1 |
| --- | --- | --- | --- | --- | --- | --- | --- | --- | --- | --- |
| Cell | ns | ns | **** | **** | ns | **** | **** | **** | **** | **** |
| Apical (M+C) | * | ns | **** | **** | ns | **** | **** | **** | **** | **** |
| Apical M | ns | ns | **** | ns | ns | **** | **** | **** | *** | **** |
| Apical C | ns | ns | **** | **** | ns | **** | **** | **** | **** | **** |
| Basal | ns | ns | **** | **** | * | **** | **** | **** | **** | **** |

| Segment length | BA17 vs BA20 | BA17 vs BA21 | BA17 vs HCA1 | BA17 vs MCA1 | BA20 vs BA21 | BA20 vs HCA1 | BA20 vs MCA1 | BA21 vs HCA1 | BA21 vs MCA1 | HCA1 vs MCA1 |
| --- | --- | --- | --- | --- | --- | --- | --- | --- | --- | --- |
| Cell | **** | **** | **** | **** | **** | ns | **** | ** | ** | **** |
| Apical (M+C) | **** | **** | **** | **** | ** | ** | **** | ns | **** | **** |
| Apical M | ns | ns | ns | ns | ns | ns | ns | ns | ns | ns |
| Apical C | **** | **** | **** | **** | * | * | **** | ns | **** | **** |
| Basal | **** | **** | **** | **** | ns | ns | **** | ns | **** | **** |

| Segment surface area | BA17 vs BA20 | BA17 vs BA21 | BA17 vs HCA1 | BA17 vs MCA1 | BA20 vs BA21 | BA20 vs HCA1 | BA20 vs MCA1 | BA21 vs HCA1 | BA21 vs MCA1 | HCA1 vs MCA1 |
| --- | --- | --- | --- | --- | --- | --- | --- | --- | --- | --- |
| Cell | **** | **** | **** | ** | *** | ** | **** | **** | **** | **** |
| Apical (M+C) | **** | **** | **** | ns | *** | ns | **** | *** | **** | **** |
| Apical M | ns | ns | **** | ns | ns | **** | ** | **** | ns | **** |
| Apical C | **** | **** | **** | ns | * | ns | **** | * | **** | **** |
| Basal | **** | **** | **** | ns | ns | ns | **** | *** | **** | **** |

| Segment volume | BA17 vs BA20 | BA17 vs BA21 | BA17 vs HCA1 | BA17 vs MCA1 | BA20 vs BA21 | BA20 vs HCA1 | BA20 vs MCA1 | BA21 vs HCA1 | BA21 vs MCA1 | HCA1 vs MCA1 |
| --- | --- | --- | --- | --- | --- | --- | --- | --- | --- | --- |
| Cell | **** | **** | **** | **** | * | **** | **** | **** | **** | **** |
| Apical (M+C) | **** | **** | **** | **** | *** | *** | **** | **** | **** | **** |
| Apical M | ns | ns | **** | ns | ns | **** | **** | **** | * | **** |
| Apical C | **** | **** | **** | **** | ns | *** | **** | **** | **** | **** |
| Basal | **** | **** | **** | **** | ns | **** | **** | **** | **** | **** |

**Supplementary Table 7:** Statistical comparisons of dendritic diameter values per distance from soma between regions in the apical M (**a**), apical C (**b**), basal (**c**) and axonal (**d**) compartments from graphs shown in Figure 7 C-F.

**a.**
| $\mu\text{m}$ | Apical M dendritic diameter | | | | | | | | | |
| --- | --- | --- | --- | --- | --- | --- | --- | --- | --- | --- |
|  | BA17<br>vs<br>BA20 | BA17<br>vs<br>BA21 | BA17<br>vs<br>HCA1 | BA17<br>vs<br>MCA1 | BA20<br>vs<br>BA21 | BA20<br>vs<br>HCA1 | BA20<br>vs<br>MCA1 | BA21<br>vs<br>HCA1 | BA21<br>vs<br>MCA1 | HCA1<br>vs<br>MCA1 |
| 20 | - | - | - | - | - | - | - | - | - | - |
| 30 | **** | *** | **** | **** | ns | **** | **** | **** | **** | **** |
| 40 | **** | *** | **** | *** | ns | **** | **** | **** | **** | **** |
| 50 | *** | ** | **** | * | ns | **** | **** | **** | **** | **** |
| 60 | *** | ** | **** | ns | ns | **** | **** | **** | **** | **** |
| 70 | **** | **** | **** | ns | ns | **** | **** | **** | **** | **** |
| 80 | **** | *** | **** | ns | ns | **** | **** | **** | **** | **** |
| 90 | **** | ** | **** | ns | ns | **** | **** | **** | *** | **** |
| 100 | **** | ** | **** | ns | ns | **** | **** | **** | *** | **** |
| 110 | **** | ** | **** | ns | ns | **** | **** | **** | ** | **** |
| 120 | **** | *** | **** | ns | ns | **** | **** | **** | * | **** |
| 130 | **** | * | **** | ns | ns | **** | **** | **** | ns | **** |
| 140 | **** | * | **** | ns | ns | **** | **** | **** | ns | **** |
| 150 | **** | ** | **** | * | ns | **** | ** | **** | ns | **** |
| 160 | **** | * | **** | * | ns | **** | ** | **** | ns | **** |
| 170 | **** | ** | **** | ** | ns | **** | * | **** | ns | **** |
| 180 | **** | * | **** | ** | ns | **** | * | **** | ns | **** |
| 190 | ** | ns | **** | ns | ns | **** | ns | **** | ns | **** |
| 200 | ** | ns | **** | ns | ns | **** | ns | **** | ns | **** |
| 210 | *** | ns | **** | ns | ns | **** | ns | **** | ns | **** |
| 220 | *** | ns | **** | * | ns | **** | ns | **** | ns | **** |
| 230 | * | ns | **** | ns | ns | **** | ns | **** | ns | **** |
| 240 | ns | ns | **** | ns | ns | **** | ns | **** | ns | **** |
| 250 | ns | ns | **** | ns | ns | **** | * | **** | ns | **** |
| 260 | ns | ns | **** | ns | ns | **** | * | **** | ns | **** |
| 270 | ns | ns | **** | ns | ns | **** | ** | **** | ns | **** |

b.
| $\mu\text{m}$ | Apical C dendritic diameter | | | | | | | | | |
| --- | --- | --- | --- | --- | --- | --- | --- | --- | --- | --- |
|  | BA17<br>vs<br>BA20 | BA17<br>vs<br>BA21 | BA17<br>vs<br>HCA1 | BA17<br>vs<br>MCA1 | BA20<br>vs<br>BA21 | BA20<br>vs<br>HCA1 | BA20<br>vs<br>MCA1 | BA21<br>vs<br>HCA1 | BA21<br>vs<br>MCA1 | HCA1<br>vs<br>MCA1 |
| 20 | ns | ns | ns | ** | ns | ns | **** | ns | ** | * |
| 30 | ns | ns | ns | **** | ns | ns | **** | ns | **** | *** |
| 40 | ns | ns | ns | **** | ns | ns | **** | ns | **** | **** |
| 50 | * | ns | ** | **** | ns | ns | **** | ns | **** | **** |
| 60 | ns | ns | ** | **** | ns | ns | **** | ns | **** | **** |
| 70 | *** | ** | **** | **** | ns | ns | **** | ns | **** | **** |
| 80 | ** | * | **** | **** | ns | *** | **** | ns | **** | **** |
| 90 | ** | ns | **** | **** | ns | **** | **** | ** | **** | **** |
| 100 | ** | ns | **** | **** | ns | **** | **** | **** | **** | **** |
| 110 | ns | ns | **** | **** | ns | **** | **** | **** | **** | **** |
| 120 | ns | ns | **** | **** | ns | **** | **** | **** | **** | **** |
| 130 | ns | ns | **** | **** | ns | **** | **** | **** | **** | **** |
| 140 | ns | ns | **** | **** | ns | **** | **** | **** | **** | **** |
| 150 | ns | ns | **** | **** | ns | **** | **** | **** | **** | **** |
| 160 | ns | ns | **** | ** | ns | **** | **** | **** | **** | **** |
| 170 | ns | ns | **** | **** | ns | **** | **** | **** | **** | **** |
| 180 | ns | ns | **** | **** | ns | **** | **** | **** | **** | **** |
| 190 | ns | ns | *** | *** | ns | **** | **** | **** | **** | **** |
| 200 | ns | ns | *** | **** | ns | **** | **** | **** | **** | **** |
| 210 | ns | ns | * | **** | ns | *** | **** | **** | **** | **** |
| 220 | ns | ns | ns | **** | ns | ** | **** | **** | **** | **** |
| 230 | ns | ns | ns | ** | ns | *** | **** | **** | **** | **** |
| 240 | ns | ns | * | * | ns | ** | **** | **** | **** | **** |
| 250 | ns | ns | ns | ns | ns | **** | **** | **** | *** | **** |
| 260 | ns | ns | ns | ns | ns | **** | **** | **** | **** | **** |
| 270 | ns | ns | ns | ns | ns | *** | ** | *** | ** | **** |

C.
| μm | Basal dendritic diameter |  |  |  |  |  |  |  |  |  |
| --- | --- | --- | --- | --- | --- | --- | --- | --- | --- | --- |
|  | BA17<br>vs<br>BA20 | BA17<br>vs<br>BA21 | BA17<br>vs<br>HCA1 | BA17<br>vs<br>MCA1 | BA20<br>vs<br>BA21 | BA20<br>vs<br>HCA1 | BA20<br>vs<br>MCA1 | BA21<br>vs<br>HCA1 | BA21<br>vs<br>MCA1 | HCA1<br>vs<br>MCA1 |
| 20 | ** | **** | ** | **** | ns | ns | **** | ns | **** | **** |
| 30 | *** | **** | **** | **** | ns | ns | **** | ns | **** | **** |
| 40 | ** | **** | **** | **** | ns | ns | **** | ns | **** | **** |
| 50 | ** | **** | **** | **** | ns | ns | **** | ns | **** | **** |
| 60 | ** | *** | **** | **** | ns | ns | **** | ns | **** | **** |
| 70 | * | ** | **** | **** | ns | *** | **** | ns | **** | **** |
| 80 | ** | * | **** | **** | ns | **** | **** | *** | **** | **** |
| 90 | *** | ** | **** | **** | ns | **** | **** | **** | **** | **** |
| 100 | *** | * | **** | **** | ns | *** | **** | **** | **** | **** |
| 110 | *** | * | **** | **** | ns | *** | **** | **** | **** | **** |
| 120 | ** | ** | **** | **** | ns | **** | **** | **** | **** | **** |
| 130 | * | ns | **** | **** | ns | **** | **** | **** | **** | **** |
| 140 | * | ns | **** | **** | ns | *** | **** | **** | **** | **** |
| 150 | * | ns | **** | **** | ns | ** | **** | **** | **** | **** |
| 160 | ns | ns | **** | **** | ns | *** | **** | **** | **** | **** |

d.
| $\mu\text{m}$ | Axon dendritic diameter | | | | | | | | | |
| --- | --- | --- | --- | --- | --- | --- | --- | --- | --- | --- |
|  | BA17<br>vs<br>BA20 | BA17<br>vs<br>BA21 | BA17<br>vs<br>HCA1 | BA17<br>vs<br>MCA1 | BA20<br>vs<br>BA21 | BA20<br>vs<br>HCA1 | BA20<br>vs<br>MCA1 | BA21<br>vs<br>HCA1 | BA21<br>vs<br>MCA1 | HCA1<br>vs<br>MCA1 |
| 20 | **** | **** | **** | * | ns | *** | **** | ** | **** | **** |
| 30 | **** | **** | **** | ns | ns | **** | **** | **** | **** | **** |
| 40 | * | * | **** | ns | ns | **** | **** | **** | **** | **** |
| 50 | ns | ns | *** | ns | ns | **** | **** | **** | **** | **** |
| 60 | ns | ns | **** | ns | ns | **** | **** | **** | ** | **** |
| 70 | ns | ns | ns | ns | ns | **** | **** | **** | **** | **** |
| 80 | ns | ns | ns | ns | ns | *** | **** | ** | **** | **** |

**Supplementary Table 8:** Statistical comparisons of segment diameter/length/surface area and volume values per branch order in the different dendritic compartments, from graphs shown in Figure 8 A, B and Supplementary Figure 7.

|  |  | Segment diameter |  |  |  |  |  |  |  |  |  | Segment length |  |  |  |  |  |  |  |  |  |
| --- | --- | --- | --- | --- | --- | --- | --- | --- | --- | --- | --- | --- | --- | --- | --- | --- | --- | --- | --- | --- | --- |
|  |  | BA17<br>vs<br>BA20 | BA17<br>vs<br>BA21 | BA17<br>vs<br>HCA1 | BA17<br>vs<br>MCA1 | BA20<br>vs<br>BA21 | BA20<br>vs<br>HCA1 | BA20<br>vs<br>MCA1 | BA21<br>vs<br>HCA1 | BA21<br>vs<br>MCA1 | HCA1<br>vs<br>MCA1 | BA17<br>vs<br>BA20 | BA17<br>vs<br>BA21 | BA17<br>vs<br>HCA1 | BA17<br>vs<br>MCA1 | BA20<br>vs<br>BA21 | BA20<br>vs<br>HCA1 | BA20<br>vs<br>MCA1 | BA21<br>vs<br>HCA1 | BA21<br>vs<br>MCA1 | HCA1<br>vs<br>MCA1 |
| Apical<br>main<br>shaft | O1 | ** | * | **** | ns | ns | **** | **** | **** | ** | **** | ns | ns | *** | ns | ns | **** | ns | * | **** | ns |
|  | O2 | *** | * | **** | ns | ns | **** | ns | **** | ns | **** | ns | ns | ns | ns | ns | ns | ns | ns | ns | * |
|  | O3 | ** | ns | **** | ns | ns | **** | * | **** | ns | **** | ns | ns | *** | ns | ns | * | ns | * | ns | ns |
| Apical<br>collateral | O1 | ns | ns | **** | **** | ns | **** | **** | **** | **** | **** | **** | **** | **** | **** | ns | ns | **** | ns | * | *** |
|  | O2 | ns | ns | **** | **** | ns | **** | **** | **** | **** | **** | **** | **** | **** | *** | ** | ns | **** | * | ns | **** |
|  | O3 | ns | ns | **** | **** | ns | **** | **** | **** | **** | **** | **** | **** | **** | ns | ** | ns | **** | ns | **** | **** |
|  | O4 | ns | ns | *** | *** | ns | **** | **** | ** | **** | **** | **** | ns | *** | ns | ns | ns | **** | ns | ** | **** |
| Basal<br>dendrite | O1 | **** | *** | ns | **** | ns | *** | **** | ** | **** | **** | **** | **** | ** | ns | ns | ns | **** | ns | **** | **** |
|  | O2 | ns | ns | ns | **** | ns | ** | **** | * | **** | **** | * | **** | **** | ** | ns | **** | ns | ns | ns | ns |
|  | O3 | ns | ns | ** | **** | ns | ns | **** | ns | **** | **** | **** | **** | **** | ns | ns | **** | ** | *** | ** | **** |
|  | O4 | ns | ns | **** | **** | ns | **** | **** | **** | **** | **** | **** | **** | **** | ns | ** | ** | **** | **** | *** | **** |
|  | O5 | ns | ns | **** | **** | ns | **** | **** | **** | **** | **** | **** | **** | **** | ns | * | ns | **** | ns | **** | **** |
|  | O6 | ns | ns | ** | ns | ns | **** | **** | ** | **** | **** | *** | ns | * | ns | **** | ns | **** | ns | *** | **** |

|  |  | Segment area |  |  |  |  |  |  |  |  |  | Segment volume |  |  |  |  |  |  |  |  |  |
| --- | --- | --- | --- | --- | --- | --- | --- | --- | --- | --- | --- | --- | --- | --- | --- | --- | --- | --- | --- | --- | --- |
|  |  | BA17<br>vs<br>BA20 | BA17<br>vs<br>BA21 | BA17<br>vs<br>HCA1 | BA17<br>vs<br>MCA1 | BA20<br>vs<br>BA21 | BA20<br>vs<br>HCA1 | BA20<br>vs<br>MCA1 | BA21<br>vs<br>HCA1 | BA21<br>vs<br>MCA1 | HCA1<br>vs<br>MCA1 | BA17<br>vs<br>BA20 | BA17<br>vs<br>BA21 | BA17<br>vs<br>HCA1 | BA17<br>vs<br>MCA1 | BA20<br>vs<br>BA21 | BA20<br>vs<br>HCA1 | BA20<br>vs<br>MCA1 | BA21<br>vs<br>HCA1 | BA21<br>vs<br>MCA1 | HCA1<br>vs<br>MCA1 |
| Apical<br>main<br>shaft | O1 | ns | ns | ns | ns | ns | ns | ns | ns | ns | * | * | ns | **** | ns | ns | **** | **** | **** | ** | **** |
|  | O2 | ns | ns | **** | ns | ns | **** | ns | **** | ns | ns | ns | ns | **** | ns | ns | **** | ns | **** | ns | *** |
|  | O3 | ns | ns | **** | ns | ns | **** | ns | **** | ns | ns | * | ns | **** | ns | ns | **** | ns | **** | ns | * |
| Apical<br>collateral | O1 | **** | **** | **** | **** | ns | * | ns | ns | ns | **** | **** | **** | **** | ns | ns | **** | **** | *** | **** | **** |
|  | O2 | **** | **** | **** | ns | ** | ns | **** | *** | **** | **** | **** | **** | **** | ns | * | ns | **** | **** | **** | **** |
|  | O3 | **** | **** | **** | ns | ** | ns | **** | ** | **** | **** | **** | **** | **** | ns | * | ns | **** | **** | **** | **** |
|  | O4 | *** | ns | **** | ns | ns | ns | **** | * | *** | **** | *** | ns | **** | ns | ns | ns | **** | *** | **** | **** |
| Basal<br>dendrite | O1 | **** | **** | **** | **** | ns | * | **** | * | **** | **** | **** | **** | **** | **** | ns | *** | **** | ** | **** | **** |
|  | O2 | *** | **** | **** | ns | * | **** | **** | ns | **** | **** | **** | **** | **** | **** | * | *** | **** | ns | **** | **** |
|  | O3 | **** | **** | **** | ns | ns | **** | **** | **** | **** | **** | **** | **** | **** | *** | ns | **** | **** | **** | **** | **** |
|  | O4 | **** | **** | **** | ns | * | **** | **** | **** | **** | **** | **** | **** | **** | ns | ns | **** | **** | **** | **** | **** |
|  | O5 | **** | **** | **** | ns | ns | ns | **** | ns | **** | **** | **** | **** | **** | ns | ns | * | **** | *** | **** | **** |
|  | O6 | *** | ns | *** | ns | ** | ns | **** | * | **** | **** | ** | ns | **** | ns | ns | ns | **** | *** | **** | **** |

**Supplementary Table 9:** Statistical comparisons of segment diameters and length values per branch order and dendritic segment type (intermediate/terminal), from graphs shown in Figure 8 C-D.

|  |  | Terminal segment diameter |  |  |  |  |  |  |  |  |  | Terminal segment length |  |  |  |  |  |  |  |  |  |
| --- | --- | --- | --- | --- | --- | --- | --- | --- | --- | --- | --- | --- | --- | --- | --- | --- | --- | --- | --- | --- | --- |
|  |  | BA17<br>vs<br>BA20 | BA17<br>vs<br>BA21 | BA17<br>vs<br>HCA1 | BA17<br>vs<br>MCA1 | BA20<br>vs<br>BA21 | BA20<br>vs<br>HCA1 | BA20<br>vs<br>MCA1 | BA21<br>vs<br>HCA1 | BA21<br>vs<br>MCA1 | HCA1<br>vs<br>MCA1 | BA17<br>vs<br>BA20 | BA17<br>vs<br>BA21 | BA17<br>vs<br>HCA1 | BA17<br>vs<br>MCA1 | BA20<br>vs<br>BA21 | BA20<br>vs<br>HCA1 | BA20<br>vs<br>MCA1 | BA21<br>vs<br>HCA1 | BA21<br>vs<br>MCA1 | HCA1<br>vs<br>MCA1 |
| Terminal<br>Apical C | O1 | ns | ns | **** | **** | ns | **** | **** | ns | **** | **** | **** | **** | **** | ns | **** | **** | **** | ns | **** | **** |
|  | O2 | ns | ns | **** | **** | ns | **** | **** | **** | **** | **** | **** | **** | **** | ns | **** | **** | **** | ns | **** | **** |
|  | O3 | ns | ns | **** | **** | ns | **** | **** | **** | **** | **** | **** | **** | **** | ns | *** | **** | **** | ns | **** | **** |
|  | O4 | ns | ns | **** | ** | ns | **** | **** | ** | **** | **** | **** | ** | **** | ns | ns | ns | **** | ns | *** | **** |
| Terminal<br>Basal | O2 | ns | ns | *** | ** | ns | **** | ** | ns | **** | **** | **** | ns | **** | ns | * | ns | **** | * | ns | **** |
|  | O3 | ns | ns | **** | **** | ns | **** | **** | **** | **** | **** | **** | **** | **** | ns | ** | ns | **** | ns | **** | **** |
|  | O4 | ns | * | **** | **** | ns | **** | **** | **** | **** | **** | **** | **** | **** | ns | **** | ns | **** | ** | **** | **** |
|  | O5 | ns | ns | **** | **** | ns | **** | **** | **** | **** | **** | **** | **** | **** | ns | **** | **** | **** | ns | **** | **** |
|  | O6 | ns | ns | ** | ns | ns | **** | **** | *** | **** | **** | *** | ns | ns | ns | **** | ns | **** | ns | ** | *** |

**Supplementary Table 10:**
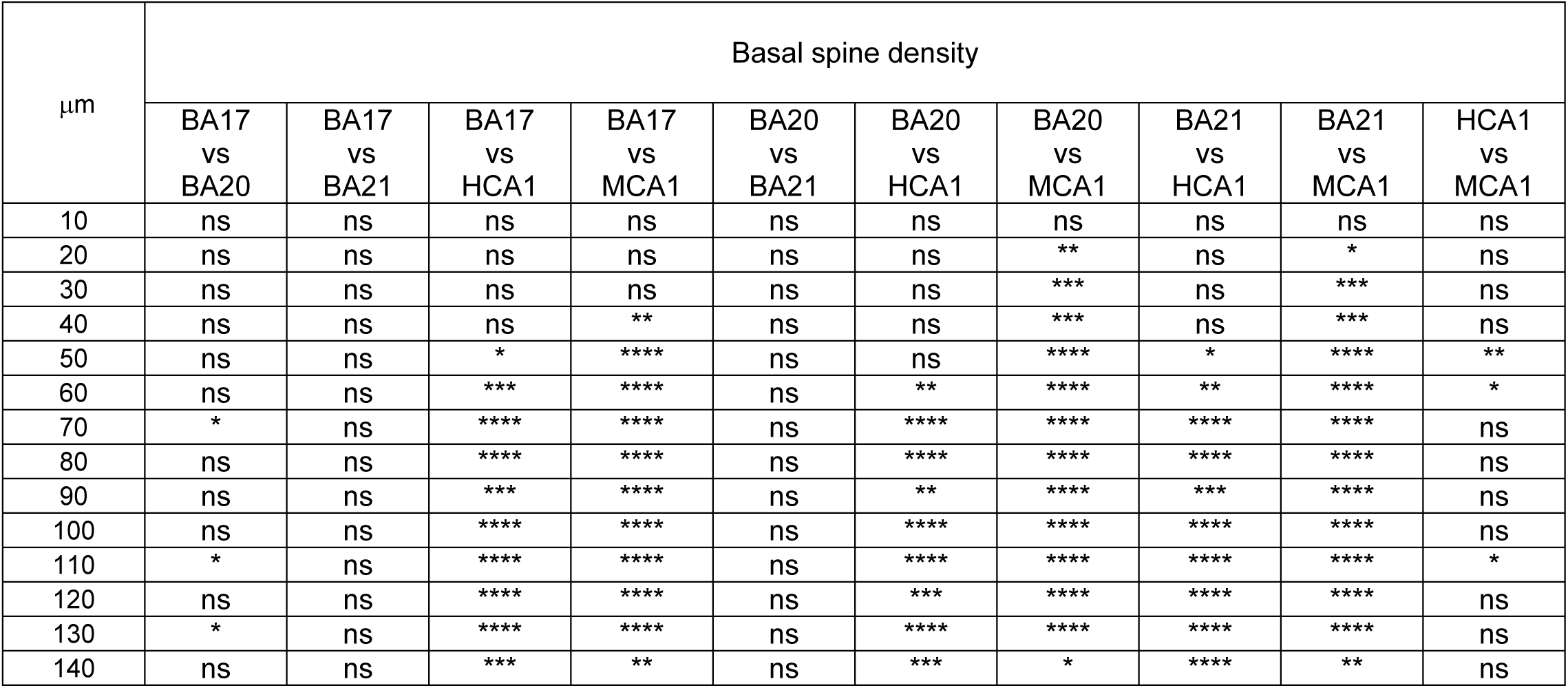
Statistical comparisons of basal spine density values per distance from soma from graph shown in Figure 9C.

**Supplementary Table 11:**
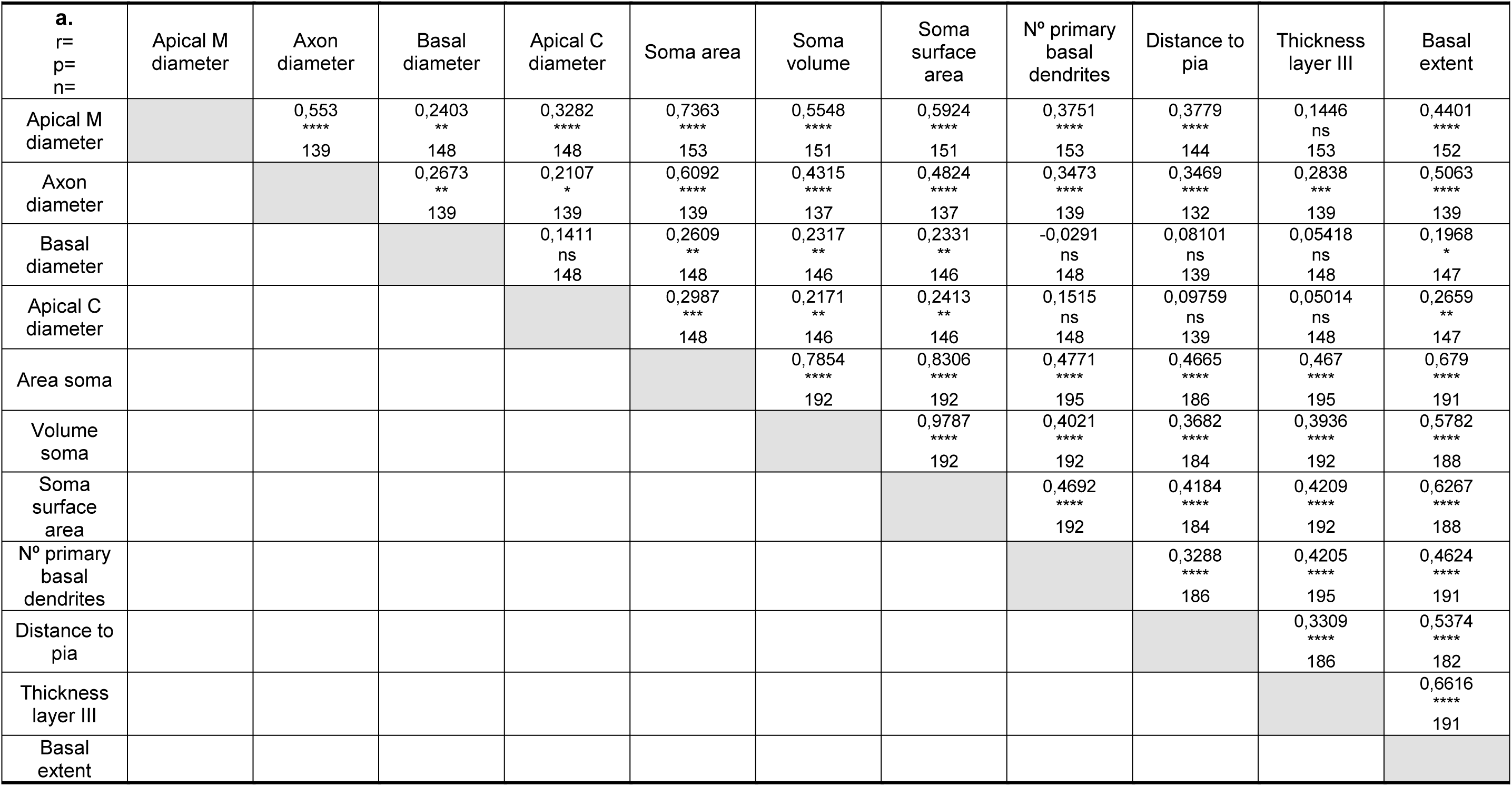
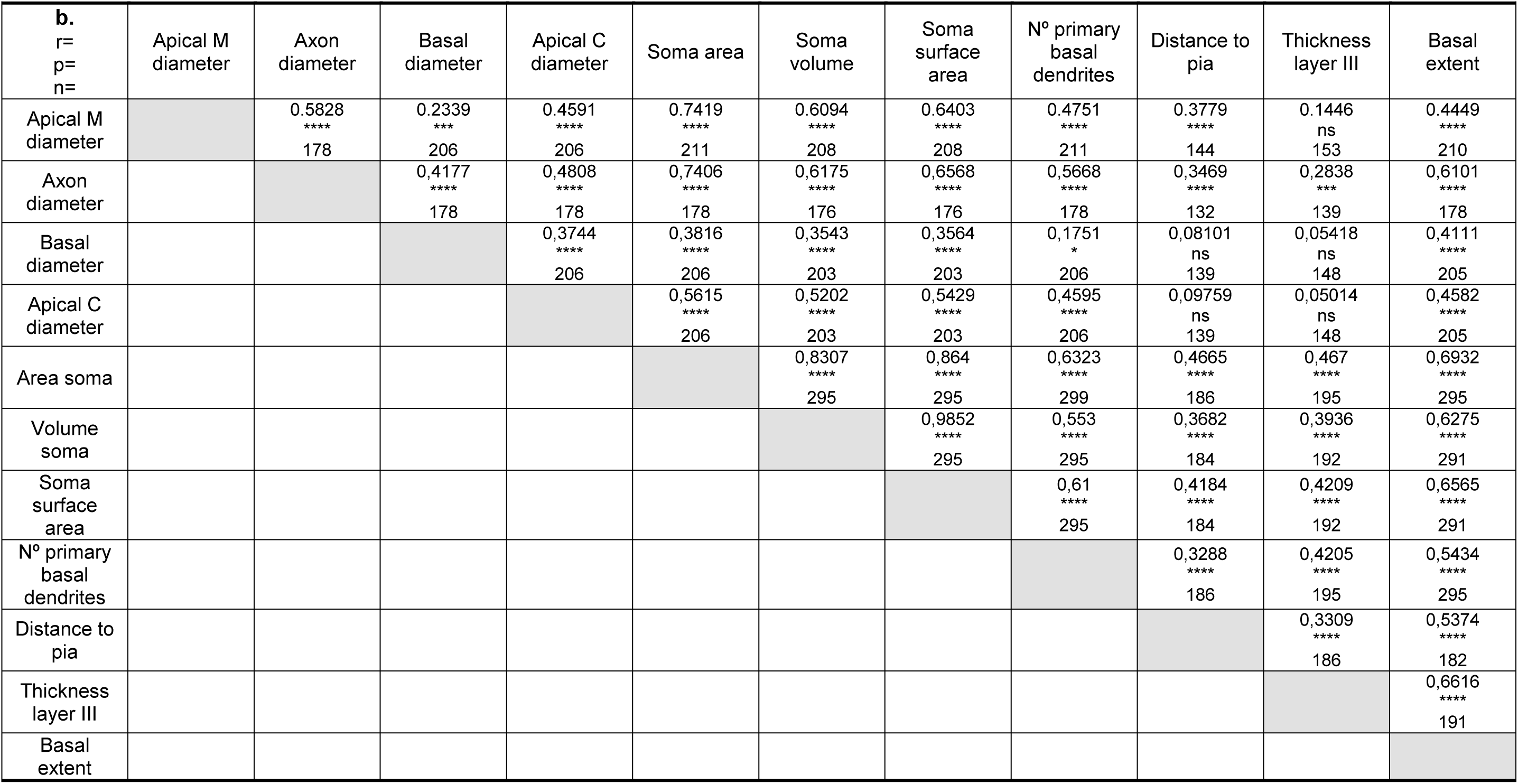
Spearman correlation analyses between various morphological parameters analyzed from human BA17, BA20 and BA21 (**a**) and human BA17, BA20, BA21, HCA1 and MCA1 (**b**) regions from graphs shown in Figure 10 and Supplementary figures 13-17. r= spearman correlation coeficient, p= signifficance, n= number of values.

